# Automated segmentation of renal structures using brightfield images of unstained kidney biopsies

**DOI:** 10.64898/2026.11.16.766951

**Authors:** Mariana Carvalho, Isabel Brandão, Roberto Silva, João P. Oliveira, Inês S. Alencastre, Jaime S. Cardoso

## Abstract

The automated segmentation of unstained kidney brightfield microscopy images remains a significant challenge in medical imaging. In conventional histopathology, staining provides crucial structural and molecular contrast, allowing renal compartments and structures to be readily distinguished. The absence of staining markedly reduces this contrast, obscuring the fine architectural details that are essential for diagnosis. Addressing this gap would reduce reliance on staining protocols, reduce technical and interpretive variability, and preserve valuable tissue for downstream molecular or multi-omics analyses. Ultimately, automation of unstained image segmentation could enable faster, more objective tissue analysis while preserving samples for further molecular testing.

In this work, we provide a comprehensive review of state-of-the-art segmentation methods applied to renal brightfield microscopy images. Current findings highlight a predominance of Convolutional Neural Network (CNN) based architectures for semantic segmentation approaches while recent trends point to the growing application of transformer-based and instance-segmentation strategies. However, the field is still hampered by the scarcity and poor annotation of publicly available renal datasets. Aiming to further contribute to the enhancement of the field, a dataset of 18 Whole Slide Images (WSIs) of unstained human kidney tissue is made publicly available. Additionally, in the dataset, we provide 543 annotated tiles extracted from the WSIs, facilitating further research into stain-independent segmentation approaches.

As a third contribution, we present a preliminary case study, evaluating state-of-the-art deep learning segmentation architectures, including U-Net and Segment Anything Model (SAM). Our preliminary results indicate that deep learning models demonstrate promising segmentation performance in unstained kidney biopsies. In alignment with these findings, advanced customised loss functions and data augmentation were integrated into the supervised training of the promising U-Net, DeepLabv3+, and TransUNet models, achieving significant improvements compared to the baseline experiments. To exploit the abundance of unlabelled data, we further tested FixMatchSeg, a semi-supervised learning method; however, its segmentation performance remained slightly behind the supervised methods.

Taken together, these contributions establish a foundation for future research in stain-independent Computational Pathology, demonstrating that unstained brightfield microscopy images can reliably support downstream tissue analysis tasks. Our dataset, validation experiments, and methodological insights provide the community with both resources and benchmarks to advance robust, data-efficient, and generalisable kidney tissue segmentation.

## 1. Introduction

The kidney biopsy is considered the gold standard for diagnosing, staging, and assessing the prognosis of Chronic Kidney Disease (CKD) [1]. Biopsies provide direct histological insight into the extent of chronic damage, including glomerulosclerosis, interstitial fibrosis, and tubular atrophy, offering critical information about the likely trajectory of kidney function and overall organ survival [2].

Renal biopsies are typically obtained using a 14-gauge needle, the largest needle size commonly used for this purpose, which extracts a core of tissue with a diameter of approximately 2.11 mm [3]. To diagnose kidney disease, pathologists must carefully assess structural changes in the glomeruli, tubules, vessels, and interstitium, as well as the presence and distribution of specific substances in the nephron, often using histological stains to highlight specific tissue components and pathological features. Figure 1a presents a brightfield image, with brightfield microscopy being the most widely used method in renal pathology. This technique relies on transmitted light to illuminate the biopsy sample, allowing for detailed visualisation of the kidney tissue architecture. Staining relies on chemical reactions that facilitate the microscopic visualisation of “structures, cells, organelles, minerals, microorganisms, and carbohydrates” [4]. The Hematoxylin and Eosin (H&E) staining technique, which has been shown to be insufficient for certain applications, is complemented with special stains such as Periodic Acid-Schiff (PAS) and Masson’s trichrome (TRI) [5]. The PAS is the best stain to evaluate glomeruli, and the TRI stain is particularly helpful in evaluating the amount of fibrous connective tissue [6].

**Figure 1:**
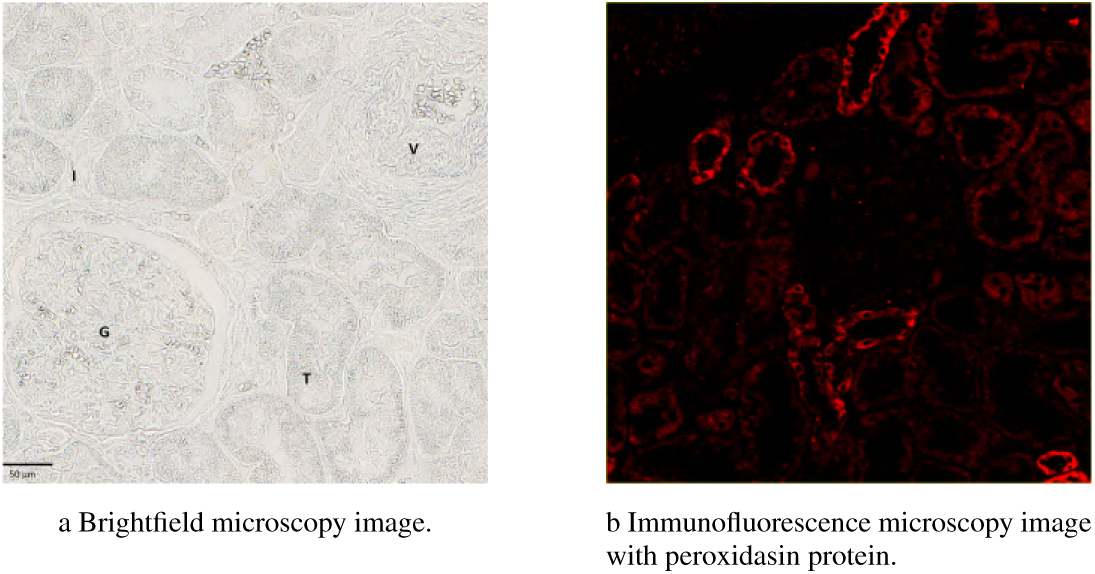
Representative micrographs of human kidney biopsy tissue acquired with brightfield and immunofluorescence microscopy. Images were obtained using a NanoZoomer S60 Digital Slide Scanner (Hamamatsu Photonics, Hamamatsu, Japan) at 40× magnification. Scale bar: 50 µm. Legend: G, glomerulus; T, tubule; V, vessel; I, interstitium.

Recognising kidney structures is essential for accurate analysis of renal biopsies [7]. Automatic segmentation of renal samples can aid in biomedical research and in routine clinical tasks such as quantification and morphology assessment of the tissue structures. Besides that, it can significantly accelerate the diagnostic process of renal disease that would otherwise rely on the labour-intensive work of pathologists. However, developing reliable models for automatic segmentation presents several challenges. Renal structures affected by disease can exhibit a wide range of appearances, with variations in both the type and severity of lesions. For example, glomerular lesions can affect different regions within the same glomerulus, resulting in significant variability even among adjacent glomeruli. This structural diversity introduces substantial complexity when designing algorithms capable of accurately segmenting kidney biopsy samples.

Current histopathologic evaluation of kidney tissue samples relies heavily on the manual assessment of structures by pathologists [5]. This approach is not only time-consuming but also prone to significant intra-operator variability, potentially affecting diagnostic consistency. Additionally, tissue staining adds another layer of complexity, increasing both the time and cost of the diagnostic process. Pathologists often use multiple staining techniques, each optimised to highlight specific tissue components. While this approach provides a more comprehensive assessment, it also introduces challenges for developing stain-independent models for automatic tissue segmentation, as the resulting images can vary significantly depending on the stain used [5]. Moreover, continuous staining and staining removal processes produce hazardous chemical waste, which is damaging to the environment.

Immunofluorescence microscopy is a technique primarily used to detect antigens, such as proteins, within tissue samples, and serves as a valuable supplement to standard histopathological assessment [8]. For instance, the peroxidasin (PXDN) protein has recently been identified as being ubiquitously expressed in all compartments of human kidney tissue from the earliest stages of nephrogenesis, with particularly high expression in the tubular compartment [9], as can be observed in Figure 1b. When using multiple microscopy techniques, such as brightfield and immunofluorescence, pathologists often rely on adjacent slides, which are pairs of slides prepared from the same tissue section. One slide is typically stained for conventional histopathology, while the other is labelled with fluorescent markers for protein localisation. This approach allows for a comprehensive assessment but introduces a significant technical challenge: aligning the microscopic structures between adjacent slides. This mapping process is both time-consuming and prone to error, as it requires precise spatial correlation at a microscopic scale.

Given this context, developing a model to automatically segment a brightfield image of an unstained kidney biopsy would be highly relevant. This would provide pathologists with a faster, less expensive, and more accurate method for the correct assessment of histopathological alteration of kidney structures, therefore impacting disease diagnosis. However, to date, little research has been published concerning the segmentation of kidney biopsy images, and to our knowledge, no studies target unstained tissue.

This work distinguishes itself by evaluating segmentation methodologies through the lens of their applicability to unstained kidney microscopy images. We conduct a comprehensive survey of foundational segmentation research and specialised architectures tailored to renal brightfield imaging. This includes an analysis of commonly employed pre-processing and post-processing techniques, as well as the performance metrics and datasets used for model development and evaluation. Our review highlights continuous advancements in the field of Computer Vision (CV), with models becoming increasingly efficient, robust, and accurate in segmenting Regions of Interest (RoIs). Despite these developments, research focused on unstained renal tissue remains limited. To help close this gap, we publicly release HUKID, a dataset comprising 18 Whole Slide Images (WSIs) of unstained kidney tissue. From these WSIs, we extract and annotate 543 tiles of 1024×1024 pixels, providing a valuable resource for the medical imaging community. Given the scarcity and annotation demands of such datasets, this contribution is intended to support further exploration of stain-independent segmentation techniques. Additionally, we conduct preliminary experiments to assess the feasibility of segmenting unstained renal tissue using state-of-the-art deep learning models. These initial investigations lay the groundwork for the advanced experiments conducted with new loss functions that incorporate boundary supervision into the training of specialised segmentation networks. We also delve into semi-supervised learning to increase our training data samples while avoiding the need for further annotation effort. The findings of this study form the foundation for for future research into automated analysis of unstained histological samples.

## 2. Overview of segmentation methodologies, datasets and performance metrics

The following section provides an overview of traditional and current methodologies used for the segmentation task of renal structures present in histopathological images. As mentioned above, no research was found on the segmentation of unstained kidney tissue. Consequently, most explored methodologies were tested on stained renal datasets. However, the architecture of these models, which is promising for the automated segmentation of unstained samples, can be further enhanced with pre- and post-processing techniques, which are also discussed in this section.

### 2.1. Traditional methodologies

Barisoni et al. introduced the Nephrotic Syndrome Study Network’s digital pathology scoring system [10]. The authors describe a descriptor-based methodology to guide pathologists in profiling renal biopsy morphology and validating these descriptors as potential predictors of diagnosis, prognosis, and response to treatment. The study involved twelve experienced pathologists who assessed fifty-one glomerular descriptors and achieved moderate to excellent concordance.

A more automated approach is the work of Marée et al. [11], which explored a bio-image workflow that aimed to facilitate a large-scale analysis of renal biopsy slides. Their method utilised a combination of Icy and Cytomine software to automate glomeruli detection through geometric invariance based on this renal structure’s morphology. On the other hand, Kline et al. used the Chan-Vese optimisation and a region-based approach to segment kidney biopsy images of Nile Red-stained samples [12]. Their semi-automated method allows users to define starting points for glomeruli detection. The authors propose their strategy as a valuable input for deep learning models since it demonstrated excellent agreement with manual segmentation, specifically for glomeruli, but also for tubules and interstitium.

Despite the advancements, early kidney structure segmentation methods, such as those in previous studies, face significant challenges. These approaches produce highly variable results between research groups, due to differences in staining and imaging protocols. Moreover, these methods rely heavily on specialised pathologists’ manual work, requiring arduous labour and resulting in slow and expensive processes. These issues led to the exploration of deep learning techniques.

### 2.2. Deep learning methodologies

The development and spread of deep learning techniques have enabled their rapid adaptation to medical imaging applications. This boom has provoked the extensive study of how deep learning models can solve biomedical segmentation tasks, aiming to extract RoIs from medical images to later be used for the assessment of patients’ health.

Recent studies show that semantic and instance segmentation are the most common segmentation methods used for the automated analysis of histopathologic images. Semantic segmentation classifies each pixel into a class label, whereas the instance method involves detecting and segmenting each object in an image. Despite having different approaches, both segmentation methods employ downsampling and upsampling techniques.

Downsampling reduces the spatial dimensions of the model’s input image while retaining essential features and capturing high-level content so that the model can differentiate between classes or identify individual object instances. This technique effectively reduces the computational complexity of these deep learning models since, by nature, their training data sizes, number of layers and parameters are very high. The upsampling technique reconstructs the spatial resolution of altered images to their original size. This step is crucial for the models to predict at the pixel level or to precisely outline object boundaries, ensuring instances are well-separated. Consequently, the models’ architecture must balance computational efficiency while preserving important spatial information.

#### 2.2.1. Semantic Segmentation Models

Ronneberger et al. were the first to introduce the application of U-Net’s architecture specifically tailored for biomedical image segmentation [13]. The authors describe this architecture as a Convolutional Neural Network (CNN)-based model with a U-shape encoder-decoder architecture, consisting of a more or less symmetric contracting and expansive path. The convolutional layers in this architecture extract information from the data structures that represent images, denoted feature maps. In each convolutional layer, filters are applied to these feature maps for the model to detect patterns, such as edges or textures. Typically, U-Net employs two successive convolutional operations per group of layers. An activation function follows these operations, usually Rectified Linear Unit, ReLU, which introduces non-linearity and encourages the model to learn more complex patterns in the data. In U-Net, consecutive downsampling steps occur on the contractive path, where the spatial dimensions of feature maps are reduced. In contrast, the number of feature maps increases, capturing more detail and high-level features in an image. On the expansive path, consecutive upsampling operations are applied to the previously altered feature maps. The goal is to return the feature maps’ spatial dimensions to their original size. U-Net also incorporates skip connections to preserve high-level and low-level spatial information, enabling data to bypass intermediate computations and connecting earlier layers to later ones. This model is commonly used as a backbone for more complex methodologies, but nowadays, it is rarely used without complementary and more advanced techniques.

Following Ronneberger et al. foundational work, various studies have leveraged semantic segmentation architectures to analyse biomedical images. Hermsen et al. studied the application of U-Net to segment kidney structures [14]. The authors used an ensemble of five U-Net models and applied spatial and colour augmentation techniques. This approach aims to invest in the robustness of the architecture against variation of tissue morphology and staining while avoiding overfitting.

The output form of U-Net is a probability map where each pixel indicates the likelihood of the pixel being within an imaging object. Moreover, U-Net has an under-segmentation limitation, where blobs – distinct objects – close to each other tend to be detected as a single blob, resulting in a lower number of blobs than the true quantity. This is an issue for images that contain many overlapped or touching blobs. For instance, Xu et al. extended the U-Net architecture to address the issue of under-segmentation in overlapping regions [15]. The researchers employed optimal thresholding as a post-processing step, achieving improved detection accuracy for tightly packed blobs.

CAMEL is a weakly supervised learning framework that combines multiple instance learning with segmentation models like U-Net and DeepLabv2 to segment lymph nodes [16]. DeepLab is a family of semantic segmentation models that has iteratively produced enhanced versions such as DeepLabv2 or DeepLabv3+. Google developed these models to segment at the pixel level. DeepLab’s key feature is the introduction of atrous (or dilated) convolution, where the idea is to broaden the filters’ field of view to incorporate a larger context without increasing the number of the model’s parameters, which could worsen the model’s performance. In this case study, there are only image-level labels, such as “In this image there is a cancerous adenoma” [16], which does not provide pixel-level information. Their approach enriches these weak annotations to produce approximate pixel-level labels that are later the segmentation models’ input. CAMEL achieves comparable results to fully supervised models, demonstrating the value of using weakly labelled data, which is easier and less expensive to obtain.

Other architectures have also been explored for the segmentation of kidney tissue. SegNet has an encoder-decoder structure similar to U-Net, where the encoder extracts features and patterns through convolutional layers. At the same time, the decoder reconstructs the spatial dimensions of the feature maps, producing segmentation masks. There are, however, some key differences between the models. U-Net uses skip connections to transfer spatial information from lower to higher-level layers. In contrast, SegNet stores max-pooling indices from the encoder, which are later used in the upsampling layers, removing the need to learn the upsampling process. This alteration simplifies the model’s overall architecture but increases the chance of losing fine detail. For instance, SegNet and DeepLabv3+ were employed to assess global glomerulosclerosis in PAS-stained kidney biopsy images [17]. To enhance the performance of these semantic segmentation models, the authors incorporated pre-processing techniques, including colour normalisation and data augmentation, resulting in significant improvements over the models’ baseline accuracy. The best results on the segmentation of non-sclerotic glomeruli were obtained using DeepLabv3+, while for sclerotic glomeruli, the best-performing model was SegNet. Bueno et al. compared the performance between U-Net and SegNet for pixel-level segmentation, considering both a two- and three-class problem [18]. The two-class problem consists of classifying pixels into two labels, glomerular and non-glomerular structures. On the other hand, the three-class problem differentiates between pixels in normal glomerular, sclerotic glomerular and non-glomerular structures. SegNet was the best-performing approach for the two-class segmentation problem, with an accuracy of 98.58%. The results from this segmentation were introduced into a fine-tuned AlexNet model – a classification network – to characterise the glomeruli into sclerotic or non-sclerotic glomeruli. This study further emphasised the role of pre-processing techniques, such as Reinhard’s colour normalisation and data augmentation, in enhancing model performance. Although the sequential pipeline of this approach of applying a semantic segmentation model, followed by a classification model, achieves good results, it introduces additional computational complexity and may not be the most optimised approach.

Both Zeng et al. [19] and Hossain and Sakib [20] segmented renal cells using U-Net-based approaches. However, Zeng et al. proposed using different networks to segment different structures, selecting U-Net, DenseNet, LSTM-GCNet, and 2D V-Net [19]. Although it produces good results, it becomes a very complex pipeline. 2D V-Net is an adaptation of V-Net, which has a similar structure to U-Net but was designed and optimised for volumetric segmentation tasks. On the other hand, DenseNet and LSTM-GCNET have very different structures from U-Net. The former is a type of CNN that reuses features through its connections. In contrast, the latter combines a Long Short-Term Memory unit with graph convolutional neural networks, creating a model capable of learning temporal dependencies. Hossain and Sakib generated synthetic annotated data based on non-synthetic data references [20]. The authors used the Speed Up Robust Features (SURF) algorithm to refine the synthetic data and remove patches that are less similar to the reference patch. Finally, the refined synthetic patches were fed into U-Net. Testing the pipeline on synthetic and non-synthetic data shows that the model performs better on synthetic data, which is not ideal for real-life applications.

Kang et al. extended the U-Net architecture with curriculum learning to improve tumour segmentation in prostate cancer samples [21]. They used a coarse-to-fine approach, generating multi-scale labels and then incrementally training the model by feeding it coarse-to-fine labels. The principle of curriculum learning is training a model from easier samples to harder samples, and it should iteratively converge to the best local minima of the non-convex latent space. However, measuring the “hardness” of data samples is not easy, representing a critical bottleneck to adapting curriculum learning to the tumour segmentation task. Although the authors arguably managed to capture these concepts in their experimental datasets, it may not be possible to do so with other examples.

Spronck et al. were the first to adapt the well-known nnU-Net image segmentation framework for tasks in the field of histopathology [22]. The nnU-Net framework has a pre-processing pipeline that requires files in NIfTI format, so the extracted RoIs and annotation masks from Whole Slide Images (WSIs) were converted to this format. Moreover, the pipeline steps in this process needed to be adapted to histology data since nnU-Net was originally designed to work with radiology images. These data types differ since histology images are usually coloured and at a microscopic scale, while radiology images are in grayscale and are typically captured at a macroscopic scale.

Both Li et al. [23] and Fuster et al.[24] studied how to adapt segmentation models to different tissues. Li et al. used a self-supervised method to optimise a pre-trained U-Net on batch normalisation layers, enhancing segmentation performance across diverse tissue types [23]. On the other hand, Fuster et al. employed TRI-CNN, a domain-adaptive model, and explicitly used a multi-scale CNN backbone with VGG16 encoders [24]. Additionally, the authors used active learning, a variant of supervised learning involving human interaction during training, which enables the selective annotation of the most informative samples. Although its performance was slightly lower than models trained on fully annotated datasets, the active learning strategy enabled effective segmentation using only 59% of the training data. Nevertheless, this type of learning requires actual human interaction with the model, which may not be feasible.

Bouteldja et al. proposed a stain-augmentation technique to tackle robustness against different stains and achieve stain-independent performance for U-Net segmentation of kidney structures [25]. To accomplish this, the authors assumed there was a pre-trained CNN alongside its training data. Then, they incorporated CycleGANs for stain translation and re-trained the original CNN on the generated stain-augmented data. However, image registration in alpha-smooth muscle actin (aSMA) and collagen type III (Col3), two of the target stains, was sensitive to slide artefacts, which made manual optimisation necessary. Stain translation required training for specific stains, and it still showed poor performance for challenging structures such as vessels. The pre-trained CNN revealed large performance decreases on both the stain-translated and registered images compared with real PAS-stained data, showing the dissimilarity between original images and their synthesised translations. In contrast, stain augmentation did not show any of those limitations, demonstrating high segmentation performance in all stains and structures. Conversely, stain augmentation requires the availability of the training data from the pre-trained CNN, which could present a potential limitation of this approach if the underlying annotated training data is not shared.

One of the most recent studies on semantic segmentation of kidney structures presents a Pediatric Kidney Diagnosis (APKD) system [26]. The segmentation module of this system, which is the relevant part for this analysis, was built with an SCNet architecture, using ResNet50 as a backbone. The architecture and backbone model were chosen as they performed the best compared to other state-of-the-art models in segmenting kidney structures. ResNet50 was used to extract robust feature maps from the input images, and the additional layers in SCNet performed further processing for a more refined result. This combination demonstrated very good segmentation results for various kidney structures, such as glomeruli, proximal tubules, and distal tubules, when tested on PAS- and H&E-stained tissue samples.

Most deep learning techniques for segmenting kidney structures require extensive manual annotations and focus on single-stain domains. This problem was addressed by introducing a generative self-supervised meta-learning framework for multi-stained multi-tissue segmentation in renal biopsy WSIs, tackling domain variability [5]. Their framework comprised two main phases – the domain-specific feature representation for each stain and the domain-invariant feature representation across multiple stain domains. First, the authors performed data augmentation by employing CycleGAN-based models to generate synthetic images matching different domains’ staining characteristics. This approach increased the robustness and generalisability of the model without additional manual annotations. Then, self-supervised learning was used to pre-train the segmentation network that extracted features of multi-tissue from stain-level data. In this step, domain-specific feature representations were learned. Finally, a small number of the pixel-level data was used to construct a few-shot meta-learning scheme. This strategy was based on both the virtual images generated by the CycleGAN models for stain transformation and the self-supervised model. This meta-learning approach consisted of randomly extracting data from different domains to form various tasks and consequently learning domain-invariant feature representations of tissues under multiple stains. Their framework used pair-wise stain transformation models to address the training limitations directly with mixed samples from different stain domains, which can lead to subpar segmentation performance. In general, their approach demonstrated favourable results; however, when the network was trained on a single stain domain and tested on data from other domains, it presented poor performance, showing that it did not achieve sufficient robustness against inter-domain variability. Nevertheless, this framework offers a meaningful solution, surpassing classic advanced segmentation networks and recently proposed few-shot segmentation methods.

#### 2.2.2. Instance Segmentation Models

Instance segmentation models are characterised by their ability to identify and segment individual objects in images, often providing bounding boxes and even pixel-level masks for those instances. Generally, instance segmentation models combine object-detection and pixel-level segmentation mechanisms, enabling them to distinguish between overlapping objects, which is often encountered in tissue samples. On the contrary, semantic segmentation models may struggle to identify that an overlapped region may present different instances and, consequently, classify that whole area as being of the same class, even if it is not the case.

Faster R-CNN is an object-detection model that can be extended for segmentation tasks. This model was built on the foundational architectures of R-CNN and Fast R-CNN. R-CNN proposed R-CNN as one of the first models for object-detection based on region proposals [27]. These proposals are areas in the image suspected to contain objects and fed through a pre-trained CNN for feature extraction. These features are then put through a classification model that determines the object’s class. One year later, Fast R-CNN was developed [28]. This architecture improves the slow process of individually computing the proposed regions by processing an entire image at a time. It inputs the image into a CNN that generates feature maps, and then regions are proposed by mapping spatial coordinates into those feature maps. Finally, Faster R-CNN was created, and it introduced the Region Proposal Network (RPN), a computationally efficient algorithm that extracts region proposals from feature maps similarly to Fast R-CNN [29].

Kawazoe et al. used Faster R-CNN to detect glomeruli from tissue-stained biopsy images [30]. Their framework localises these structures by drawing bounding boxes around them. Differences in biopsy and staining techniques between pathology laboratories can cause variations in the structures’ appearance, so the model should be robust against these variations. Inception-ResNet, pre-trained on the Common Objects in Context (COCO) dataset, was chosen as the model for feature map generation, as it fulfilled the authors’ robustness requirement. The experiment revealed that the model achieved high performance for various stains. However, it is computationally slower than the You Only Look Once (YOLO) model or a single-shot multi-box detector (SSD).

These object-detection models may be complemented by others that provide pixel-level masks to obtain a new method for segmentation. One such extension is Mask R-CNN, a framework that extends Faster R-CNN by adding a branch specifically for pixel-wise segmentation. In addition to generating bounding boxes, this approach predicts binary masks for each detected object. This is achieved by an additional head that operates in parallel with the class prediction and bounding box heads. One improvement over Faster R-CNN is the Region of Interest Alignment, RoI Align. This technique aligns candidate RoIs using bilinear interpolation, ensuring that the feature maps extracted for each RoI align correctly with the original image, improving the model’s accuracy. For instance, Cascade Mask R-CNN architecture was implemented for glomeruli instance segmentation [7]. This type of structure includes multiple detection and segmentation stages, each refining and improving the results from the previous stage. Similar to Feng et al., ResNet50 was used as a backbone for feature extraction, providing efficient segmentation. Despite reaching good results, the study also has some limitations. First, the researchers show that the imbalanced classes prevent the model from accurately segmenting instances of the rarest class. Moreover, the developed model has only been trained on glomeruli that have suffered noticeable changes in their structures, so it may not perform as expected when detecting glomeruli that present less severe or no lesions.

The work of Sun et al. experimented with Deep Attention Integrated Networks, DAINets, to segment nuclei [31]. They addressed the problem of variations, caused by different staining protocols, uneven nucleus intensity, background noises, and nucleus tissue differences in biopsy specimens. Their method was mainly built on self-attention modules – spatial and channel attention – and introduced a feature fusion branch to fuse high-level representations with low-level features for multi-scale perception. The mark-based watershed algorithm was also applied to refine the predicted segmentation maps, removing fragmented regions or noise. An Adversarial Learning strategy was used to encourage the predicted mask to be iteratively refined and become more similar to the observed mask. In the training procedure, they employed pre-processing strategies such as conventional data augmentation, individual colour normalisation and random image crop. In the test procedure, they applied patch-based extraction, individual colour normalisation, and a multi-scale zoom strategy to tackle the staining variation problem in specimens. The methodology was evaluated on the Multi-Organ Nucleus Segmentation (MoNuSeg) dataset, and DAINet performed better than previous methods.

Salvi et al. proposed to combine detection and segmentation models with active shape models [32]. The chosen Softmax-Driven Active Shape Model (SD-ASM) used information from deep neural networks to initialise and evolve a dynamic deformable model that seeks to match the shape of the target objects. In practice, their detection branch used Faster R-CNN to generate bounding boxes, which were used to initialise each active shape model (ASM). Their segmentation branch uses ResUNet, an architecture that enables the network to encode multi-scale contextual information and gradually recover the spatial information by capturing object boundaries. To address the challenge of touching and partially overlapping instances, the authors integrated ASMs into the pipeline, introducing a repulsive interaction term in the mathematical formulation of each ASM. Despite ASMs’ computational simplicity and ease of control, they are sensitive to noise, initialisation, and the complexity of the segmented object. Consequently, they can get stuck in local minima and achieve suboptimal convergence. To further optimise their framework, the authors suggested a Generative Adversarial Network to address the task of stain normalisation, where the goal is to translate the colour pattern of images from one domain to another, where the origin domain exhibits a wide range of colour patterns, and the target domain has a relatively uniform colour pattern. This approach was tested on the MoNuSeg and CoNIC datasets, where it consistently outperformed reference methods. To emphasise versatility, the authors successfully applied SD-ASM to segment renal tubules, where individual structure identification is critical.

GlomNet is a network designed for glomeruli instance segmentation [33]. While previous methods have shown strong performance in glomerular segmentation tasks, the authors highlight a significant limitation: when objects are close to each other, these methods often fail to separate the individual glomerular instances. This limitation illustrates that semantic segmentation alone is insufficient to correctly count the number of glomeruli in a single image. To address this problem, the authors proposed an altered U-Net architecture with an EfficientNet as its encoder – a CNN-based network that uses a compound coefficient technique to uniformly scale the network’s depth, width, and resolution. To tackle instance separation, the network includes two decoders, one that produces a binary segmentation mask and the other that generates horizontal and vertical distance maps, representing the distance of each pixel to its respective centre of mass. These outputted distance maps were used to determine object boundaries and, consequently, to separate glomerular instances. To perform this operation, the authors extracted glomeruli boundaries using a Sobel operator that computes horizontal and vertical derivatives. These boundaries were subtracted from the binary mask obtained from the first decoder, and the result was used to calculate the Euclidean Distance Transformed map. Finally, the peaks in this map were extracted and used as markers for the watershed algorithm to split the mask into separate instances. This method was trained and tested on 176 images collected from two clinical centres, outperforming two state-of-the-art methods – EfficientUNet and Cascade Mask R-CNN. These results demonstrate that strong performances can be achieved with less intricate solutions, compared to a more complex approach such as Mask R-CNN, which requires extensive computation to resolve instance overlaps.

#### 2.2.3. From CNN-based networks to transformers and foundation segmentation models

There is a family of transformer models whose structure differs from that of CNNs. This architecture works with a self-attention mechanism, which allows the model to focus on the most relevant parts of the data during processing. This mechanism computes scores for elements in the data, learning relationships and dependencies between them. Although there is some research on the use of transformer-based models for medical imaging segmentation, it is far scarcer than the literature on CNN-based models.

Xie et al. proposed a novel hierarchically structured transformer encoder, which outputs multi-scale features, denoted by the authors as SegFormer, Simple and Efficient Design for Semantic Segmentation with Transformers [34]. This architecture is known for its simplicity compared to traditional CNNs. One key advantage of SegFormer is that it does not need positional encoding, a mechanism usually used in transformer-based models to inject information about the position of elements. This characteristic avoids the need to interpolate positional codes when the testing resolution differs from the training resolution, which may lead to decreased performance. The SegFormer’s decoder is a Multi-Layer Perceptron (MLP) that aggregates information from different layers, thus combining both local attention and global attention to render powerful representations. However, this framework presented a significant computational limitation. Although the smallest SegFormer model has 3.7 million parameters and is smaller than the standard CNN model, it is unclear whether it can perform effectively on an edge device with only one hundred kilobytes of memory.

Kawazoe et al. refined their previous approach using SegFormer [35]. They designed a computational pipeline to detect kidney structures from WSIs of PAS-stained tissue samples. The first step in their method was to draw bounding boxes surrounding renal structures in a WSI using Faster R-CNN. Then, SegFormer was used to segment the structures, classifying the pixels in bounding boxes into five classes – Bowman’s space, glomerular tuft, crescentic region, sclerotic region, and background. Data augmentation techniques, such as flipping, Gaussian blurring, and sharpening, were applied to train the network and improve its robustness for variations in morphology and staining. Although the authors performed colour normalisation and data augmentation techniques, extended methods were required to compensate for the minute differences in WSIs between pathology laboratories.

The Segment Anything Model (SAM) is a recently developed foundation model for image segmentation, achieving an outstanding performance on an extensive range of segmentation tasks. This model is built on Vision Transformer (ViT) models and comprises three components: “an image encoder, a flexible prompt encoder, and a fast mask decoder” [36]. The image encoder is a pre-trained ViT Masked Autoencoder (MAE) [37] and runs once per image, creating an image embedding. The model considers sparse prompts such as points, boxes, and text, as well as dense prompts in the form of masks. The second block, the prompt encoder, represents prompts through different strategies. Points and boxes are encoded by positional encodings combined with learned embeddings. In contrast, text prompts are encoded using a pre-trained Contrastive Language-Image Pre-training (CLIP) text encoder [38]. Masks are incorporated through convolutions, which are then directly added to the image embedding. The mask decoder is the last block of SAM’s architecture, and it maps the image and prompt embeddings with an output token to a segmentation mask. This is built with a modified Transformer decoder block that uses prompt self-attention and bi-directional attention between embeddings (prompt-to-image and image-to-prompt). This block runs two times to refine the embeddings. Then, the image representation is upsampled, and a multilayer perceptron (MLP) maps the output token to a dynamic linear classifier. This classifier outputs the foreground probability at each image location. The resultant segmentation masks are ranked by estimated IoU, and the model outputs the three with the highest confidence score to deal with prompt ambiguity.

Despite demonstrating an overall good performance, SAM shows varying accuracy when applied directly to medical image segmentation tasks. Since its introduction, fine-tuning strategies have been developed to adapt the model for medical domains, such as the SAM adapter and MedSAM. In particular, MedSAM presented by [39] addresses the limitation that current segmentation foundation models face when applied to medical segmentation tasks due to differences in natural and medical images. Despite SAM being built for promptable segmentation, using it without providing prompts and accomplishing fully automatic segmentation is possible. In contrast, MedSAM requires prompts to produce an output, and consequently, the model may not be applicable depending on the task’s objectives. Furthermore, both this version and the SAM adapter require costly computational resources.

The Segment Anything Model 2 (SAM 2) was introduced by Ravi et al. [40]. It focuses on generalising the original SAM version to promptable video segmentation. Like its predecessor, SAM2 has an image encoder that is run once for the entire interaction. This block is an MAE [37] pre-trained Hiera ([41]; [42]). This hierarchical encoder allows the use of multiscale features during encoding. SAM2 generates segmentation masks for individual images and video frames, which are processed one at a time. To adapt SAM to the video domain, memory blocks were created to store contextual information from previous frames. This memory helps refine the predicted masks for the object of interest. However, when SAM2 is applied to images, the memory becomes empty. The memory mechanism is spread across three blocks. The first is memory attention, where current frame features are conditioned on new prompts and on contextual information from previous frames and predictions. Before information is gathered, encoded and stored in the memory bank, it must be collected from the newly produced segmentation masks. These are generated based on the prompt embeddings and outputted by the mask decoder. The prompt encoder works exactly like SAM’s version. In contrast, the mask decoder has “an additional head that predicts whether the object of interest is present in the current frame” [40]. Moreover, there are skip connections from the image encoder to the mask decoder that bypass the memory attention and provide high-resolution embeddings for decoding.

det-SAM is an architecture that combines object detection and instance segmentation techniques to address the challenges of histopathological analysis of kidney biopsy images [43]. This model was built based on SAM’s advantages in graphical discernibility and YOLOv8’s detection effectiveness. YOLO is an object-detection model that, unlike others, frames this task as a regression problem, achieving effective performance. Its strategy is to divide the input image into a grid and use it directly to detect objects and predict class probabilities. The authors designed a pipeline where input images were cropped first, creating small tiles. These tiles were put through YOLOv8, which detects the renal structures. Then, the Non-Maximum Suppression algorithm (NMS) is applied to eliminate redundant and overlapping bounding boxes from YOLO’s results. As mentioned above, SAM is comprised of two encoders and one decoder. In their proposed approach, one of the encoders receives the original input images, whereas the other processes NMS’s output images. The image embeddings and prompts are combined in the decoder, which outputs predictions for segmentation masks. The computational requirement of the proposed architecture is distinctively lower than that of fine-tuning SAM or SAM adapters. Moreover, det-SAM presented reliable accuracy, demonstrating a promising capability to address medical imaging segmentation tasks.

##### Discussion

Table 1 summarises the reviewed methods for segmenting stained kidney biopsy specimens. Out of the eighteen methodologies, nine are publicly accessible. The table shows that, as of recently, research groups tend to share their work, which is of great importance to the community. This enables others to gain insights from their methods and drive advancements in the field.

**Table 1:** Summary of reviewed methods for the segmentation of stained kidney biopsy specimens.

| Method | Year | Segmentation Type | Reference | Source Code |
|---|---|---|---|---|
| Glomerular Descriptors | 2016 | Manual | Barisoni et al. [10] | - |
| Geometric Invariance | 2016 | Instance | Marée et al. [11] | - |
| U-Net Ensemble | 2019 | Semantic | Hermsen et al. [14] | - |
| SegNet and DeepLabv3+ | 2020 | Semantic | Altini et al. [17] | - |
| U-Net, SegNet, and AlexNet | 2020 | Semantic | Bueno et al. [18] | - |
| U-Net, DenseNet, LSTM-GCNet, and 2D V-Net | 2020 | Semantic | Zeng et al. [19] | - |
| U-Net | 2020 | Semantic | Hossain and Sakib [20] | - |
| Chan-Vese Optimisation | 2021 | Semantic | Kline et al. [12] | https://github.com/adriennekline/renal-pathology-segmentation |
| ResNet50 | 2021 | Semantic | Yamaguchi et al. [44] | https://github.com/ryama-tokyo/glomerular_classification |
| Cascade Mask R-CNN | 2021 | Instance | Jiang et al. [7] | https://github.com/buptaicz/glomeruli-instance-segmentation |
| Faster R-CNN and SegFormer | 2022 | Instance | Kawazoe et al. [35] | https://github.com/jinseikenai/glomeruli_segmentation |
| U-Net | 2023 | Semantic | Bouteldja et al. [25] | https://github.com/NBouteldja/KidneyStainTranslation |
| U-Net | 2023 | Semantic | Li et al. [23] | https://github.com/NeuronXJTU/SFDA-CellSeg |
| det-SAM | 2023 | Instance | Ning et al. [43] | - |
| SCNet and ResNet50 | 2024 | Semantic | Feng et al. [26] | - |
| ResNet50 | 2024 | Semantic | Liu et al. [5] | https://github.com/SnowRain510/MSMTSeg |
| SD-ASM | 2024 | Instance | Salvi et al. [32] | https://data.mendeley.com/datasets/nwnfh5x5j3/1 |
| UNet | 2024 | Instance | Moreau et al. [33] | https://github.com/bozeklab/GlomNet |

Furthermore, from the summarised methodologies, it is noticeable that many of these methods focus only on glomeruli segmentation, such as [44], [7], [35], and [33]. However, Brandão et al. [9] noted that the PXDN protein is expressed throughout all compartments of kidney tissue, with particularly high levels in the tubular compartment. Therefore, all renal structures are of interest for analysis, as not only glomeruli can provide valuable insights for pathologists.

Moreover, studies [25], [32], and [5] address stain translation. In future work, their research outcomes can aid in synthetically staining our unstained samples or artificially removing the staining from other datasets to introduce them into our study.

### 2.3. Pre- and post-processing techniques

Most deep learning methodologies for tissue segmentation have a pre- and, or post-processing stage that simplifies or refines the segmentation step. Several studies have shown that introducing these techniques improves the models’ performance. Pre-processing techniques are commonly employed to prepare the input images optimally, whereas post-processing techniques usually refine the network’s output. Salvi et al. reviewed and explored the impact of these practices in digital pathology image analysis [45].

The input data for these segmentation networks are images obtained from tissue samples. To acquire digitised histological images, some steps are usually carried out to achieve a result that captures an appearance similar to the tissue’s aspect inside the living organism. The preparation steps, such as the samples’ collection, dehydration, paraffin embedding, microtomy, staining, mounting, and digitisation, can generate artefacts that potentially lower the quality of the images.

#### 2.3.1. Pre-processing techniques

The current practice to minimise sample variability is limited to procedural quality-control methods. However, these methods are insufficient to mitigate variabilities in tissue and stain appearance across diverse laboratories and different batches in the same facility. Pre-processing techniques can be included in methodologies to address the introduced artefacts.

Both Xu et al. [15] and Kawazoe et al. [35] used a Gaussian filter to remove noise in their images. Noise usually has high-frequency variations in pixel intensity, which can be smoothed by a Gaussian blur, for example.

Thresholding is a common technique used to separate tissue from the background. Many researchers analysed in previous sections, including [15], utilise Otsu’s thresholding, an algorithm that automatically determines the optimal threshold value to separate pixels into background and foreground objects.

Contrast Limited Adaptive Histogram Equalisation (CLAHE) was implemented in [16], as a pre-processing step. This technique emphasises and improves the contrast in images by dividing them into small tiles and individually applying histogram equalisation. This process transforms pixel values and redistributes them over the histogram’s range. CLAHE differentiates from standard histogram equalisation because it limits the histogram bins, highlighting some details without amplifying noise and artefacts. Moreover, CLAHE can be used to improve the results of other pre-processing techniques, such as thresholding and filters, by refining the images.

The procedure of stain colour normalisation involves transforming an input image into another image, where the intensity distribution of the original image is mapped to match the colour distribution of the target image. The Reinhard’s method is applied in [18], a global colour transformation between the target and source image that uses Principal Component Analysis. This technique involves the transfer of the target image mean colour to the source image “in a way so that the source image intensity variations are preserved, and the obtained contrast of the processed image is roughly the same as the target image contrast” [45]. Kawazoe et al. performed colour normalisation by calculating the difference between the value of a pixel and the overall mean pixel value and then dividing this result by the variance of the pixel [35]. On the other hand, Sun et al. used Individual Colour Normalisation, which normalises each colour channel individually [31]. Salvi et al. [32] explored an innovative approach described in the instance segmentation section that uses Generative Adversarial Networks to address the task of stain normalisation.

Patch selection is a technique used to extract more manageable-sized regions representative of the important features of an image. This helps reduce computational complexity and redirects the focus of the model to more relevant parts of the images. The most common patch selection strategy is to split the target WSI into a grid and adopt a sliding window approach to extract all the patches for the network training. Both [35] and [32] employed a sliding window approach, in which each windowed image is fed into the model. [35] further specified that adjacent windows should overlap by 10% to ensure complete coverage of glomeruli within the analysis. Additionally, the authors processed bounding boxes that overlapped by 35% or more by merging them into one.

#### 2.3.2. Post-processing techniques

The main challenges that current segmentation models still struggle with are the separation of overlapping structures, followed by the biological shape and size variability of renal structures. It remains difficult to precisely define object boundaries. Post-processing techniques are employed to refine the results obtained by segmentation networks.

Liang et al. used a sequence of post-processing techniques to improve their network’s original predictions [46]. First, an edge smoothing filter was applied, followed by the Overlapped Region Forecast algorithm, which merges patches to eliminate boundary errors in the original prediction. In the end, the flood fill algorithm was used to correct the predicted false negatives by internally filling the regions that the model should have segmented as an object but did not.

Hermsen et al. implemented refinements specific to renal structures [14]. The authors assigned objects smaller than three hundred pixels – the size of the smallest annotated tubule – to be of the interstitium class. Additionally, if an object consisted of multiple class labels, the most predominant class, the one exceeding at least 35% of the area, was assigned to the entire object.

Moreau et al. focused on correctly identifying separate glomeruli instances [33]. They used a Sobel operator to compute the horizontal and vertical derivatives, which were later used by a watershed algorithm that determined how to split the binary mask into instances, discriminating glomeruli and non-glomeruli. The instance segmentation section presents a more detailed explanation of their method.

### 2.4. Datasets

The segmentation of kidney structures has been explored using a variety of methodologies that have been tested on diverse datasets. These materials include in-house datasets, specifically curated for the study’s focus and typically sourced from the authors’ collaborating institutions, as well as public datasets that are widely shared and accessible to the broader scientific community. The choice of dataset can influence the generalisability and scalability of segmentation models, making it essential to analyse the datasets used by the listed methods.

#### 2.4.1. In-house datasets

In-house datasets are created by research groups and often tailored to their specific experimental needs. They typically include custom annotations, which makes them ideal for developing targeted solutions. However, their limited accessibility restricts external validation and comparative analysis.

Marée et al. created their dataset with tissue sections from two hundred slides, where one slide contained several tissue sections from the same patient [11]. The tissue was prepared using Masson’s trichrome protocol within thirteen research centres and acquired using a NanoZoomer scanner with a 40x objective.

Hermsen et al. used transplant biopsies obtained from one hundred and one patients who underwent kidney transplants in the Radboud University Medical Center, Nijmegen, the Netherlands [14]. The slides were PAS-stained and digitised using a Pannoramic 250 Flash II digital slide scanner with 20x magnification.

The Department of Emergency and Organ Transplantations (DETO) of the Bari University Hospital provided the PAS-stained kidney biopsies dataset for Altini et al. [17]. Slides were digitised using a high-resolution whole slide scanner with a 20x scanning objective. The complete dataset was composed of twenty-six kidney biopsies obtained from nineteen donors.

Zeng et al. [19] selected four hundred patients who underwent renal biopsy between 2016 and 2018 at the National Clinical Research Center of Kidney Diseases at the Jinling Hospital. A Leica Aperio ScanScope AT Turbo scanned these renal biopsies with a 40x magnification. Three experienced junior pathologists provided annotations of glomeruli, which three senior pathologists validated.

To construct their glomeruli dataset, [7] gathered kidney biopsy samples from patients diagnosed with various renal diseases from January 2018 to June 2019 in the Department of Nephrology of Peking University People’s Hospital (Beijing, China). The biopsy sections selected for model development were PASM (Periodic Schiff-Methenamine), PAS and Masson’s trichrome-stained. The digital images were obtained from two sources and were either snapshots or whole slide images. The snapshot images were saved to the researchers’ department image library and were captured at magnifications of 10×, 20×, and 40× using a Leica DFC550 digital camera (Singapore) attached to a Leica DM4000 B LED optical microscope (Weizla, Germany). The WSIs were scanned at 40x magnification using a Precie 500B scanner (UNIC Technologies Inc., Beijing, China). Due to computational limits, WSIs were cropped into overlapping 2048×2048 patches with a stride of three-fourths the window size to maximise data use and minimise glomeruli fragmentation.

Kline et al. used images of nephrectomy samples from healthy and diabetic renal pathology patients [12]. These were stained with Nile Red and captured using a Zeiss LSM 880 confocal microscope with 20x magnification. These materials were acquired from the Biobank for the Molecular Classification of Kidney Disease [47]. However, there is no public access to these data.

Yamaguchi et al. [44] collected two hundred and ninety-three PAS-stained WSIs from three hospitals: ninety-nine WSIs from the University of Tokyo Hospital from 2010 to 2017; eighty-eight WSIs from the Tazuke Kofukai Medical Research Institute at Kitano Hospital from 2014 to 2017; and one hundred and six WSIs from the University of Tsukuba Hospital from 2009 to 2016. From the two hundred and ninety-three WSIs, the glomerular image was manually extracted in a rectangular shape so that one major glomerulus was included per slide, which yielded 10202 images. Similarly to the previous study, Kawazoe et al. [35] collected PAS-stained WSIs of renal biopsy specimens from the University of Tokyo Hospital and the Kitano Hospital, gathering three hundred and fifty-three and three hundred and twenty-four WSIs, respectively. In both facilities, various kidney diseases were included in the WSIs, and the slide digitisation was conducted using a NanoZoomer C9600-12 slide scanner with 40x magnification.

Ning et al. [43] research is conducted on three Kaggle competition datasets. Despite the authors providing links to these competitions, only competitors are granted access to the data. Nevertheless, the first dataset contained human kidney micro-vasculature structures consisting of tiles extracted from fourteen WSIs, where only the tiles from five WSIs were annotated. The second dataset was dedicated to glomeruli identification. It comprised large-sized TIFF images containing annotations of glomeruli in two different formats, Run-Length Encoding (RLE)-encoded form and uncoded JavaScript Object Notation (JSON) form. The last dataset, the Functional Tissue Unit dataset, collected images of three hundred and fifty-one kidney biopsy slides, and it was used to identify the location of each functional tissue unit.

Bouteldja et al. constructed their dataset by gathering PAS-stained kidney tissue slides from different sources [25]. The obtained slides were digitised using NanoZoomer HT2 and Aperio AT2 scanners with a 20x magnification. All samples from mice, rats, and pigs came from already published studies ([48], [49], [50], [51], [52], and [53]). To complement the dataset, sixteen PAS-stained sections from human kidney specimens were anonymously obtained from the Institute for Pathology of the Rheinisch-Westfälische Technische Hochschule (RWTH) Aachen University’s archive. Eight qualified annotators generated ground truth annotations.

Moreau et al. [33] built a dataset with one hundred and seventy-six WSIs obtained from two centres. Data from Freiburg University Hospital covered forty-four WSIs from twenty-six patients. The authors thoroughly described the patients’ diagnoses: one patient had autosomal dominant polycystic kidney disease, ten had renal cancer, and fifteen had diverse nephrotic glomerular diseases. The samples were acquired with the VENTANA DP 200 software and a VENTANA DP 200 Roche Diagnostics scanner with a 20x objective. The other subgroup of the dataset consisted of one hundred and thirty-two WSIs from thirteen patients from the Cologne University Hospital, part of the ForMe registry cohort (NCT03949972). All patients from this subgroup presented a nephrotic syndrome, either Minimal Change Disease or Focal and Segmental Glomerulosclerosis. Images were acquired using NZacquire software and a Hamamatsu NanoZoomer S360 scanner with a 20x objective. Both dataset portions were PAS-stained and annotated by experts. However, the samples from Cologne University did not present precise contours since glomeruli were annotated by having an ellipse drawn around them.

Lastly, Liu et al. [5] used a total of 1008 kidney biopsy samples with four stains (PASM, PAS, H&E, and TRI) from two hundred and fifty-two patients collected from the Shanxi Provincial People’s Hospital spanning the years 2017 to 2020. All samples were digitised as WSIs using a KF-PRO-005-EX digital slide scanner with a 20x objective and a 10x eyepiece.

#### 2.4.2. Public datasets

In contrast to in-house datasets, public datasets are shared within the scientific community, enabling the comparison and validation of developed models across diverse research groups. Additionally, investigators can use these datasets in the absence of their own or even to complement their gathered samples. These characteristics drive continuous advancements in segmentation methodologies.

The digital tissue images used by Bueno et al. [18] were obtained from the Academia and Industry Collaboration for Digital Pathology (AIDPATH) Database (European project number 612471). However, no details were shared about which exact images were selected. Nonetheless, these images comprise five different PAS-stained human kidney tissue cohorts acquired and digitised from three European institutions: Castilla-La Mancha’s Healthcare Services, Andalusian Health Service and Vilnius University Hospital Santaros Klinikos. The WSIs were obtained by the Leica Aperio ScanScope CS scanner with a 20x objective.

Following a similar strategy, Hossain and Sakib [20] revealed that their Clear Cell Renal Cell Carcinoma (CCRCC) histopathology data was collected from a digital database, but, in this case, from The Cancer Image Archive (TCIA). The ten WSIs selected from the archive were digitised using Leica Aperio tools at 20x magnification scale and were utilised to generate the texture of synthetic nuclei patches. The synthesised patches were validated with the annotated Kidney Renal Clear Cell Carcinoma (KIRC) dataset [54], which was prepared from TCIA.

Li et al. [23] used three pathological image datasets in their experiments: KIRC [54], Triple Negative Breast Cancer (TNBC) [55] and a TCIA dataset [56]. As stated above, KIRC is a Kidney Renal Clear Cell Carcinoma dataset consisting of four hundred and sixty-three images of labelled pathological cells. The TNBC dataset has fifty images of Triple-Negative Breast Cancer Cells, and the last set of selected images comprises ninety-seven images of Stomach Adenocarcinoma available at TCIA. Links have been provided for the accessible download of the datasets used and are available in a repository (https://github.com/NeuronXJTU/SFDA-CellSeg).

Feng et al. [26] created a dataset from renal biopsy images obtained by the Department of Pathology, Children’s Hospital affiliated with Zhejiang University School of Medicine. They collected 2935 H&E and PAS-stained renal biopsy specimens of children diagnosed with kidney disease from 2009 to 2019. The renal biopsy tissues were scanned using a KF-PRO-400 scanner with a 10x and 40x objective. To guarantee good quality imaging for artificial intelligence development and training, the samples were manually assessed, and 988 RoIs extracted from 779 biopsy sections were selected.

##### Discussion

All the analysed datasets are summarised in Table 2 and consist of stained kidney biopsy samples. Indeed, none of the surveyed segmentation methodologies experimented with unstained kidney tissue. Staining is a standard practice in histopathology because it enhances the contrast between renal structures, which significantly aids in the analysis of tissue. Analogously, it is a helpful technique for segmentation networks since it provides clear structure boundaries. Researchers widely adopt these staining techniques despite their costly and time-consuming application process. Four of the sixteen methods used public digital image archives, whereas twelve research groups opted for in-house datasets. From the private datasets, only [5] manifested the possibility to grant access to the data upon request. The scarcity of available kidney datasets prevents the advancement and development of this digital pathology field. Collecting and preparing biopsy samples, digitising the images, and producing the corresponding annotations require specialised and meticulous work, which is very time-consuming. We aim to enrich this research field by making our unstained renal dataset publicly available. Moreover, we can complement our dataset with samples from the digital archives for future work. However, most often, what is provided in these databases are unlabelled WSIs of stained biopsy samples, meaning that to use these images, they need to undergo a process of identification of RoIs, professional annotation, and some method to address the staining.

**Table 2:** Summary of datasets of stained samples used in kidney structure segmentation studies.

| Reference | Dataset Size | Staining | Imaging | Magnification | # Classes | Availability | Annotation |
|---|---|---|---|---|---|---|---|
| Marée et al. [11] | 200 WSIs | TRI | NanoZoomer | 40x | 2 | Private | Not specified |
| Hermsen et al. [14] | 60 WSIs | PAS | Pannoramic 250 Flash II | 20x | 10 | Private | Fully annotated (at patch level) |
| Altini et al. [17] | 26 WSIs | PAS | Not specified | 20x | 2 | Private | Not specified |
| Zeng et al. [19] | 400 WSIs | PAS | Leica Aperio ScanScope AT Turbo | 40x | 4 | Private | Glomeruli annotation |
| Bueno et al. [18] | 47 WSIs | PAS | Leica Aperio ScanScope CS | 20x | 3 | https://aidpath.eu | Glomeruli annotation |
| Hossain and Sakib [20] | 10 WSIs | Not specified | Leica Aperio | 20x | Not specified | https://www.cancerimagingarchive.net/ | Nuclei annotation |
| Jiang et al. [7] | 348 WSIs | PASM, PAS<br>and Masson's trichrome | Leica DFC550 digital camera,<br>Leica DM4000 B LED optical microscope,<br>Precie 500B scanner | 10x, 20x, 40x | 3 | Private | Glomeruli annotation |
| Kline et al. [12] | 30 images | Nile Red | Zeiss LSM 880 confocal microscope | 20x | 3 | Private | Glomeruli, tubules<br>and interstitium annotation |
| Yamaguchi et al. [44] | 293 WSIs | PAS | NanoZoomer C9600-12 | 40x | 12 | Private | Glomeruli annotation |
| Kawazoe et al. [35] | 677 WSIs | PAS | NanoZoomer C9600-12 | 40x | 5 | Private | Glomeruli annotation |
| Ning et al. [43] | 365 WSIs | PAS | Not specified | Not specified | 2 | Private | Glomeruli and vessels annotation |
| Bouteldja et al. [25] | 168 WSIs | PAS | NanoZoomer HT2 and Aperio AT2 | 20x | 6 | Private | Fully annotated (at patch level) |
| Li et al. [23] | 10 WSIs | Not specified | Not specified | 40x | Not specified | https://portal.gdc.cancer.gov/ | Cell annotation |
| Moreau et al. [33] | 176 WSIs | PAS | VENTANA DP 200, Roche Diagnostics | 20x | 1 | Private | Glomeruli annotation |
| Liu et al. [5] | 1008 WSIs | PASM, PAS,<br>H&E, Masson's trichrome | KF-PRO-005-EX | 20x | 4 | Private, but access may be granted upon request | Fully annotated (at patch level) |
| Feng et al. [26] | 2935 WSIs | H&E and PAS | KF-PRO-400 | 10x, 40x | 10 | https://github.com/ChunyueFeng/Kidney-DataSet | Fully annotated (at patch level) |

### 2.5. Performance metrics

Segmentation methods should be evaluated to ensure their effectiveness and reliability. There are different metrics generally used to assess these methods’ performance, each focusing on a certain aspect to evaluate.

Intersection over Union (IoU), or Jaccard Index, is one of the most common metrics. It measures the overlap between the predicted segmentation (P) and the ground truth (G):

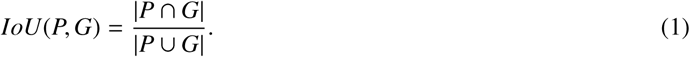

Its value ranges from zero to one and is calculated by dividing the area of overlap between the predicted and ground truth segments by the area of their union. If IoU is zero, it indicates that there was no overlap, and therefore, the segmentation was very poor. On the contrary, an IoU of one shows that there was a perfect match between the ground truth and predicted segmentation masks. Altini et al. used SegNet and DeepLab to detect and segment glomeruli, achieving a weighted IoU of 0.97, an almost-perfect glomeruli segmentation result [17]. On the other hand, Faster R-CNN and SegFormer combined demonstrated a mean IoU of 0.670 and 0.693 when experimenting with data from two facilities [35]. However, this result reflects a five-class problem, which justifies the overall lower result when compared to the previous study. The MSMTSeg framework achieved a mean IoU of 0.718 for multiple tissues under different stains, using only one annotated training sample for each stain, demonstrating promising advancements in developing stain-invariant and robust segmentation models [5].

Another common metric is the Dice Similarity Coefficient (DC), which measures the similarity between two data sets, in this case, the ground truth (G) and predicted (P) segmentation masks. DSC ranges from zero to one, and similarly to IoU, zero represents no match, and one represents a perfect match. Its value is calculated as follows:

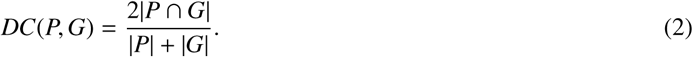

The baseline CNN approach to segmentation of kidney structures [14] achieved an overall weighted mean DSC of 0.80 and 0.84 in biopsies from two different centres. The best-segmented class was “glomeruli” in both datasets, with DCs of 0.95 and 0.94, respectively, followed by “tubulin combined” and “interstitium”. Similarly, the Chan-Vese method [12] reached an overall DSC on healthy and diabetic samples of 0.85 and 0.85, respectively. The glomeruli segmentation network, GlomNet [33], outperformed two state-of-the-art methods, EfficientUNet and Cascade Mask R-CNN, with a Dice score of 0.79. The previously mentioned framework, MSMTSeg [5], showcased a high mean Dice Similarity Coefficient of 0.836.

Precision and recall are metrics usually used to assess classification models. However, they can be borrowed to analyse segmentation at the pixel level. Precision measures the proportion of true positive pixels among all pixels classified as positive by the model:

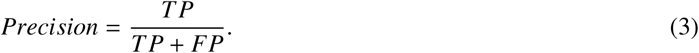

On the other hand, recall measures the proportion of true positive pixels that were correctly identified out of all observed positive pixels:

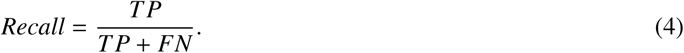

Additionally, one of the most straightforward metrics is accuracy, which calculates the ratio of correctly predicted pixels in segmentation models:

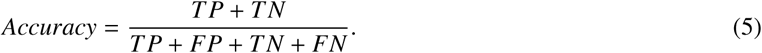

For instance, the framework proposed by Zeng et al. [19] reached an average precision of 93.1% and an average recall of 94.9% for the location of glomeruli. Moreover, this network could identify three kinds of intrinsic glomerular cells with a 92.2% accuracy, surpassing the about 5–11% average accuracy of junior pathologists. The SegNet-AlexNet approach [18] achieved a global accuracy above 92%, indicating the suitability of this CNN for detecting glomeruli in WSIs. Also, this method obtained valuable results, achieving approximately 94% and 96% accuracy for non-glomerular structures and normal glomeruli and more than 83% for sclerosed glomeruli. The authors experimented with consecutive CNNs and confirmed an increase in the performance to 98.16% accuracy, improving the previous results obtained with the three-class segmentation problem. The APKD framework [26], which is 5.5 times faster than manual detection, obtained an average accuracy of 94% for each kidney structure and 99% specifically for glomeruli using ResNet50 and SCNet. This approach is one of the most recently published studies on kidney structure segmentation and demonstrates a fast and well-performing strategy.

#### Discussion

Table 2 illustrates that most authors use datasets with varying numbers of labelled classes. This indicates their segmentation networks are designed to accommodate different numbers of target classes. For this reason, it is not possible to make a direct comparison between the methods’ performance. Nevertheless, IoU, DSC, precision, recall, and accuracy are the most commonly adopted metrics to assess segmentation performance.

Another approach to evaluate a segmentation model of renal structures is to use its output for downstream tasks. Such scenarios can include glomeruli morphometric analysis, assistance to pathologists in identifying histomorphological alterations in the structures and enabling a clinical diagnosis. Sometimes, focusing on achieving segmentation results that can fulfil these downstream tasks is more important than slightly improving the quantitative performance metrics if those would not contribute to significantly better results in clinical scenarios.

### 2.6. Summary

The analysed methodologies and respective experimental results show a continuous improvement in the field as models get faster, more robust and more accurate at segmenting renal structures, specifically glomeruli. These advancements are driven by the development and refinement of deep learning models and the introduction of novel pre- and post-processing techniques that improve the results outputted by the segmentation networks. Using diverse and multiple datasets on the models’ training aids in building a robust and generalisable method. Nevertheless, the designed methods still greatly depend on datasets that comprise stained samples. Researchers employ pre-processing techniques to address variability and artefacts introduced by the staining process, which fluctuates between facilities. However, to this day, no research has been published that concerns unstained kidney biopsy samples. Exploring this subject may show the opportunity to discard the costly and time-consuming staining process, potentially reducing variations and noise present in digitised tissue samples.

## 3. HUKID

As emphasised in the previous section, no public datasets are currently available that include samples of unstained kidney biopsies. As a contribution of this work, we make publicly available the HUKID (*HUman KIDney*) dataset (https://to_be_disclosed_upon_paper_acceptance) to encourage further work in segmenting kidney structures.

### 3.1. Samples

Formalin-fixed paraffin-embedded (FFPE) specimens of mature kidney tissue with normal histomorphological appearance and containing at least one whole glomerulus were retrieved from the archives of the Pathology Department of São João Local Health Unit and University Hospital Centre (CHUSJ), Porto, Portugal. From these samples, fourteen were recovered from kidney biopsy samples acquired for diagnostic purposes, and four were from non-tumoral kidney parenchyma of nephrectomy specimens. A margin of at least 1 cm from the tumour margins or the injured kidney parenchyma was a prerequisite for the collection of adult kidney specimens. Exclusion of developmental genitourinary anomalies and cause of death not imputable to kidney disease were conditions for preselecting the postmortem fetal and neonatal kidney specimens.

### 3.2. Brightfield microscopy

For brightfield examination, three-micrometer (3-*µ*m) thick sections were cut from each of the FFPE blocks, mounted onto pseudonymised glass microscope slides, and prepared from sequential sections according to standard histopathology techniques. In brief, the tissue sections were deparaffinised by washing in fresh xylene (3 x 10 min) and rehydrated to water by sequential exposure to decreasing concentrations of ethanol (1 x 10 min, at 100%, 96%, 70%, and 50%), before immersion in double-distilled water. The unstained slides were then used for manual segmentation of the kidney tissue compartments. PAS-stained slides, prepared according to standard techniques from sequential sections of each of the selected FFPE blocks, were used as a topographical reference for the manual segmentation and for double-checking the normal histomorphology of the study specimens (data not shown).

For indirect immunofluorescence analysis, after the deparaffinisation and rehydration steps, the slides were prepared for a specific protein immunodetection following the established protocol. The primary antibody was a rabbit polyclonal antibody (not disclosed), and the fluorophore-conjugated secondary antibody was the Donkey Anti-Rabbit IgG (H+L) Highly Cross-Adsorbed Secondary Antibody, Alexa Fluor™ Plus 647 (Invitrogen, Thermo Fisher Scientific; Waltham, MA, USA; catalog #A32795).

### 3.3. Whole slide imaging and digital image analysis

The glass microscope slides processed for brightfield or fluorescence microscopy were converted into high-resolution digital images by high-speed scanning using a Nanozoomer S60 Digital Slide Scanner (Hamamatsu Photonics; Hamamatsu, Japan) at 40x magnification. A total of 18 WSIs were produced, and 543 tiles of 1024×1024 dimensions were extracted from them. These tiles were fully annotated using the open-source QuPath software. The annotations distinguish five classes: glomeruli, tubules, vessels, interstitium, and empty space. As a contribution of this work, the 543 annotated tiles and their respective original patches, along with the 18 WSIs, are made publicly available. Each tile is identified by its coordinates and anonymized WSI ID, which helps locate the corresponding patch within the entire image.

The annotation process was extremely time-consuming, since it required experts to analyse each image and the individual structures within it. Additionally, the absence of staining resulted in low-contrast images, increasing the complexity of the analysis as the morphology of structures appeared more similar. As a result, labelling each square millimetre (*mm*^2^) took about six hours to complete manually. This effort totalled around 85 hours. Table 3 summarises information regarding the obtained WSIs, selected patches, and annotated areas. Figure 2a shows an example of a micrograph of a brightfield image, and Figure 2b demonstrates the annotation of a different micrograph, where each pixel was assigned one of the five classes. Table 4 describes information about each class and its respective annotation in the dataset.

**Figure 2:**
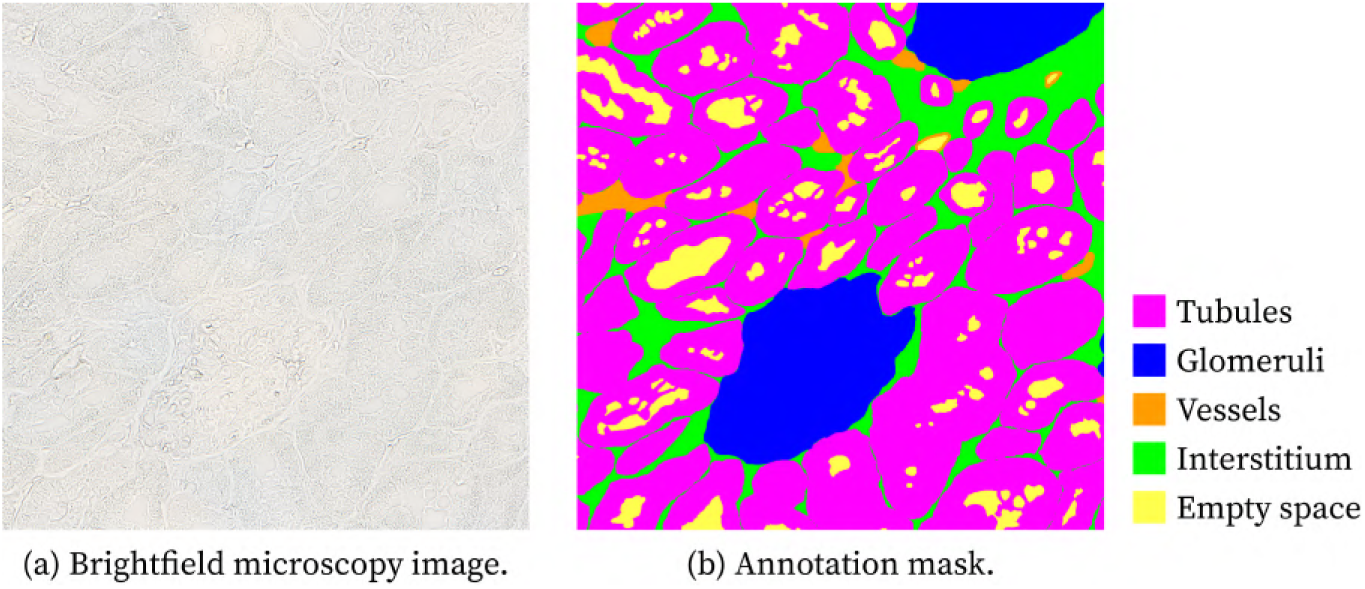
Exemplary micrographs of unstained human kidney tissue samples from histologically normal kidney tissue. After image acquisition (a), the different kidney compartments were manually segmented in unstained brightfield images using the QuPath software, presented here as a filled map of annotations (b). *Colour code: tubules (magenta), glomeruli (blue), vessels (orange), interstitium (green), and empty space (yellow)*.

**Table 3:** Dataset information.

| **WSI ID**<br>(anonymised) | **Image Size**<br>(px x px) | **Total Image Area**<br>(*µm*<sup>2</sup>) | **Total Tissue Area**<br>(*µm*<sup>2</sup>) | **Total Annotated Area**<br>(*µm*<sup>2</sup>) | **# Annotated Tiles** | **Annotated Area**<br>(% according to tissue area) | **Annotated Area**<br>(% according to whole image area) |
|---|---|---|---|---|---|---|---|
| 1 | 55552 x 48384 | 523457.778 x 10<sup>3</sup> | 275016.502 x 10<sup>3</sup> | 711.694 x 10<sup>3</sup> | 12 | 0.259 | 0.136 |
| 2 | 39424 x 26880 | 51593.024 x 10<sup>3</sup> | 8807.811 x 10<sup>3</sup> | 573.254 x 10<sup>3</sup> | 42 | 6.508 | 1.111 |
| 3 | 28672 x 32256 | 45026.639 x 10<sup>3</sup> | 3725.672 x 10<sup>3</sup> | 375.388 x 10<sup>3</sup> | 20 | 10.076 | 0.834 |
| 4 | 39424 x 37632 | 72230.233 x 10<sup>3</sup> | 7172.388 x 10<sup>3</sup> | 594.411 x 10<sup>3</sup> | 33 | 8.287 | 0.823 |
| 5 | 10752 x 48384 | 25327.484 x 10<sup>3</sup> | 5954.621 x 10<sup>3</sup> | 483.492 x 10<sup>3</sup> | 37 | 8.120 | 1.909 |
| 6 | 35840 x 21504 | 37522.199 x 10<sup>3</sup> | 1349.125 x 10<sup>3</sup> | 193.532 x 10<sup>3</sup> | 12 | 14.345 | 0.516 |
| 7 | 32256 x 37632 | 59097.464 x 10<sup>3</sup> | 4402.512 x 10<sup>3</sup> | 620.501 x 10<sup>3</sup> | 28 | 14.094 | 1.050 |
| 8 | 39424 x 26880 | 51593.024 x 10<sup>3</sup> | 2855.747 x 10<sup>3</sup> | 252.260 x 10<sup>3</sup> | 14 | 8.833 | 0.489 |
| 9 | 25088 x 53760 | 65663.849 x 10<sup>3</sup> | 14208.615 x 10<sup>3</sup> | 552.066 x 10<sup>3</sup> | 26 | 3.885 | 0.841 |
| 10 | 43008 x 21504 | 45026.639 x 10<sup>3</sup> | 6097.610 x 10<sup>3</sup> | 236.293 x 10<sup>3</sup> | 18 | 3.875 | 0,525 |
| 11 | 28672 x 21504 | 30017.759 x 10<sup>3</sup> | 3771.783 x 10<sup>3</sup> | 165.964 x 10<sup>3</sup> | 12 | 4.400 | 0.553 |
| 12 | 14336 x 48384 | 33769.979 x 10<sup>3</sup> | 7910.583 x 10<sup>3</sup> | 369.651 x 10<sup>3</sup> | 24 | 4.673 | 1.095 |
| 13 | 32256 x 26880 | 42212.474 x 10<sup>3</sup> | 7259.269 x 10<sup>3</sup> | 504.930 x 10<sup>3</sup> | 59 | 6.956 | 1.196 |
| 14 | 19712 x 16128 | 61914.360 x 10<sup>3</sup> | 5113.707 x 10<sup>3</sup> | 782.456 x 10<sup>3</sup> | 36 | 15.301 | 1.264 |
| 15 | 43008 x 40320 | 337714.696 x 10<sup>3</sup> | 166270.957 x 10<sup>3</sup> | 1297.226 x 10<sup>3</sup> | 25 | 0.780 | 0.384 |
| 16 | 46592 x 37632 | 341467.081 x 10<sup>3</sup> | 230127.560 x 10<sup>3</sup> | 1354.644 x 10<sup>3</sup> | 42 | 0.589 | 0.397 |
| 17 | 39424 x 26880 | 51593.024 x 10<sup>3</sup> | 11504.679 x 10<sup>3</sup> | 1749.959 x 10<sup>3</sup> | 63 | 15.211 | 3.392 |
| 18 | 51968 x 32256 | 326457.539 x 10<sup>3</sup> | 183584.022 x 10<sup>3</sup> | 3267.384 x 10<sup>3</sup> | 40 | 1.780 | 1.001 |

**Table 4:** Dataset information by class.

| **WSI ID**<br>(anonymised) | **Glomerulus Area**<br>(*μm*<sup>2</sup>) | **Glomerulus**<br>(%) | **Tubules Area**<br>(*μm*<sup>2</sup>) | **Tubules**<br>(%) | **Interstitium Area**<br>(*μm*<sup>2</sup>) | **Interstitium**<br>(%) | **Vessels Area**<br>(*μm*<sup>2</sup>) | **Vessels**<br>(%) | **Empty Space**<br>(*μm*<sup>2</sup>) | **Empty Space**<br>(%) |
|---|---|---|---|---|---|---|---|---|---|---|
| 1 | 62660.073 | 8.804 | 453426.110 | 63.711 | 83692.447 | 11.760 | 43550.500 | 6.119 | 68501.101 | 9.625 |
| 2 | 130412.611 | 22.750 | 291029.639 | 50.768 | 66993.018 | 11.686 | 24809.239 | 4.328 | 60012.014 | 10.469 |
| 3 | 65354.922 | 17.410 | 157658.882 | 41.999 | 59754.667 | 15.918 | 8002.064 | 2.132 | 84617.490 | 22.541 |
| 4 | 123408.053 | 20.761 | 285967.256 | 48.109 | 78557.059 | 13.216 | 20348.866 | 3.423 | 86185.484 | 14.499 |
| 5 | 92000.366 | 19.028 | 289721.447 | 59.923 | 52706.415 | 10.901 | 4779.154 | 0.988 | 44284.668 | 9.159 |
| 6 | 21143.019 | 10.925 | 111292.030 | 57.506 | 32105.821 | 16.589 | 4879.880 | 2.521 | 24111.505 | 12.459 |
| 7 | 65460.748 | 10.550 | 316144.610 | 50.950 | 71481.531 | 11.520 | 13332.577 | 2.149 | 154082.200 | 24.832 |
| 8 | 43865.730 | 17,389 | 147231.396 | 58.365 | 36697.502 | 14.548 | 5266.429 | 2.088 | 19217.769 | 7.618 |
| 9 | 80194.357 | 14.526 | 317183.672 | 57.454 | 87824.631 | 15.908 | 13902.072 | 2.518 | 53119.481 | 9.622 |
| 10 | 49062.855 | 20.764 | 131224.705 | 55.535 | 42676.149 | 18.061 | 3701.005 | 1.566 | 9684.486 | 4.098 |
| 11 | 39604.461 | 23.863 | 58090.595 | 35.002 | 30431.712 | 18.336 | 2441.794 | 1.471 | 35285.626 | 21,261 |
| 12 | 63959.733 | 17.303 | 136637.157 | 36.964 | 43450.997 | 11.755 | 10285.888 | 2.783 | 115329.452 | 31.200 |
| 13 | 150252.468 | 29.757 | 192096.357 | 38.044 | 81350.985 | 16.111 | 32628.822 | 6.462 | 48599.633 | 9.625 |
| 14 | 43037.989 | 5.500 | 178431.243 | 22.804 | 126498.588 | 16.167 | 37678.159 | 4.815 | 396792.220 | 50.711 |
| 15 | 59012.985 | 4.549 | 893786.434 | 68.900 | 223653.886 | 17.241 | 14281.619 | 1.101 | 106814.528 | 8.234 |
| 16 | 112248.529 | 8.286 | 818260.794 | 60.404 | 203637.670 | 15.033 | 34121.739 | 2.519 | 186408.402 | 13.761 |
| 17 | 102092.620 | 5.834 | 692899.041 | 39.595 | 243939.532 | 13.940 | 24585.006 | 1.405 | 687318.565 | 39.276 |
| 18 | 221129.152 | 6.768 | 2370932.896 | 72.564 | 363347.279 | 11.120 | 85385.677 | 2.613 | 226770.831 | 6.940 |

Observing the data in Table 4, there is a significant discrepancy between the area of each structure class, denoting a clear class imbalance. Tubules are by far the most common class that can be observed in Table 5, followed by glomeruli and empty space; however, there is already a large difference between the area occupied by the first class and the area occupied by glomeruli and empty space. The vessel class is highly under-represented, with an average area of 2.96% per WSI.

**Table 5:**
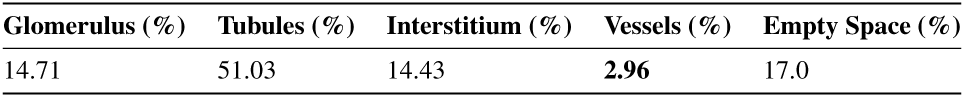
Average area that each class occupied per WSI.

### 3.4. Ethical considerations

This study uses pseudonymised human samples, being conducted in accordance with the Declaration of Helsinki. The research protocol was reviewed and approved by the institutional Health Ethics Commission (reference number CE-04-2023) and Data Protection Officer of Unidade Local de Saúde de São João (ULSSJ), Porto, Portugal. Patients’ samples were assessed only after pseudonymisation by authorised clinicians.

## 4. Setup for the experiments

We considered a semantic segmentation as the main task for all experiments, where each pixel in an image was classified into one of the five renal structure class labels: *glomeruli*, *tubules*, *vessels*, *interstitium*, and the *empty space* inside the structures.

The approach selected to create training, validation, and test subsets was to allocate images extracted from different renal biopsies exclusively to each subset. This strategy guarantees that no subset contains images from the same biopsy as another subset, preventing data leakage and adopting a more robust evaluation of model generalisation.

The original dataset includes many masks that are not fully annotated and therefore have white margins, see example of partially annotated mask in Figure 3.

**Figure 3:**
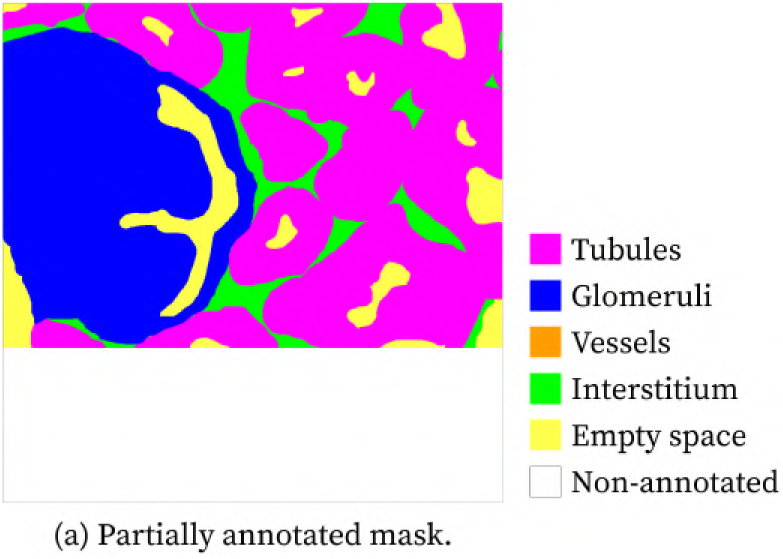
Example of an annotation mask that is not fully annotated using the open-source QuPath software. *Colour code: tubules (magenta), glomeruli (blue), vessels (orange), interstitium (green), empty space (yellow), and non-annotated tissue (white)*.

Upon conversion of the RGB masks to class labels, the masks included in the final sets and that were partially annotated had the white section mapped to an extra label. Throughout model development, this extra label was carefully treated as an *ignore label*. During loss and metrics computations, a “valid mask” was created to filter out the ignored regions, as the model should not learn from these sections.

The segmentation performance metrics we opted for were IoU, DSC, pixel accuracy, precision and recall. In addition, the partition similarity [57], a non-semantic generic metric, was adopted. Unlike class-specific metrics, it evaluates the segmentation as a whole by comparing the partitions formed by clusters of pixels in each mask. The core idea is to compute the distance between segmentation masks, which is defined as the minimum number of pixels that would have to be removed from both the predicted and ground-truth masks, so that the partitions become identical. Consequently, this metric does not evaluate the semantic correctness of the segmentation, but it does capture structural disagreements. Additionally, it can provide insight into over- and under-segmentation behaviour across different models. This metric is particularly suitable for assessing the performance of SAM and SAM2.

The symmetric partition distance is defined as:

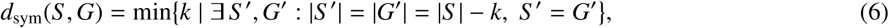

where *S* is the predicted segmentation mask, *G* is the ground-truth mask, and *k* is the minimum number of pixel removals needed for the two partitions to match exactly. A normalised partition similarity score (PS) can be defined as:

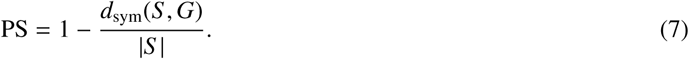

For the computation of all metrics, the *ignore index* present in the partially labelled ground-truth masks was addressed by masking out the regions that contained this label. Therefore, the evaluation was only considered for the ground-truth regions that were annotated. This metric was also calculated for the remaining methods for consistency.

## 5. Preliminary case study using state-of-the-art segmentation methods

In this section, we perform a preliminary study on whether current segmentation strategies, in particular the most promising among the surveyed ones, are sufficient to bypass the challenges of unstained renal samples and successfully segment them. We utilise the state-of-the-art architectures, SAM and U-Net, reviewed in the Related work section.

The code developed for this empirical study is also made publicly available (http://to_be_disclosed_upon_paper_acceptance).

### 5.1. Foundation models

As mentioned in section 2.2.2, SAM was developed with the aim of generalising to different segmentation tasks and achieving strong zero-shot and few-shot performance [36]. Moreover, this model focuses on providing real-time results. Therefore, we found it suitable for our case study and wanted to assess whether these positive outcomes would be reproducible. SAM was designed to be a promptable model. However, since we wanted to achieve fully automatic segmentation, we did not provide any prompt besides the images we wanted to segment in our experiments. A detailed description of SAM’s architecture is provided in section 2.2.2. In addition, the three main components of this architecture – the image encoder, the prompt encoder, and the mask decoder – are summarised in Figure 4.

**Figure 4:**
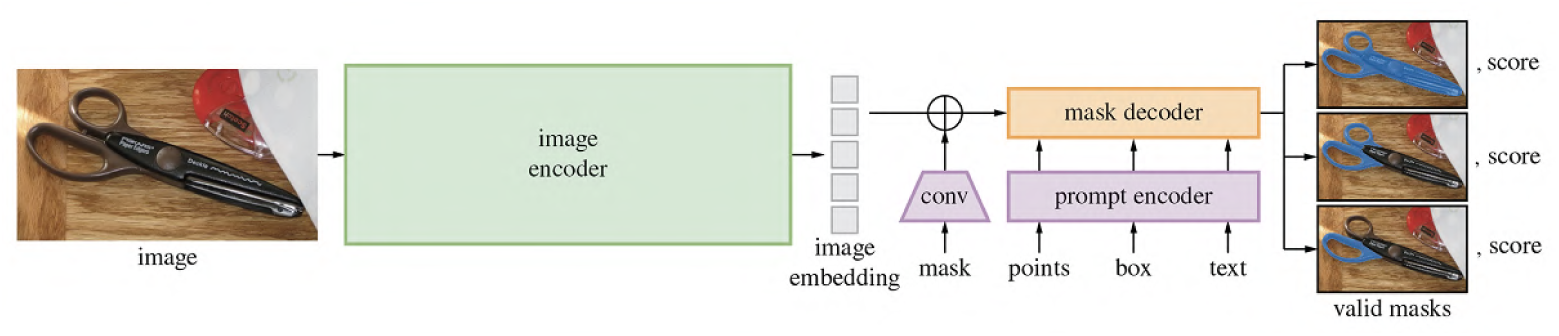
SAM architecture overview from [36], accessed 1 April 2025.

The Segment Anything Model 2 (SAM 2) is an adaptation of SAM that generalises to promptable video segmentation [40]. This model demonstrates improved performance and efficiency in image segmentation compared to SAM, achieving better accuracy and providing results six times faster. For this reason, it was also considered for our experiments.

Figure 5 illustrates the architecture of SAM2, which resembles SAM’s version, but with additional blocks that introduce memory into the network. For a more detailed description of this architecture, refer to section 2.2.2.

**Figure 5:**
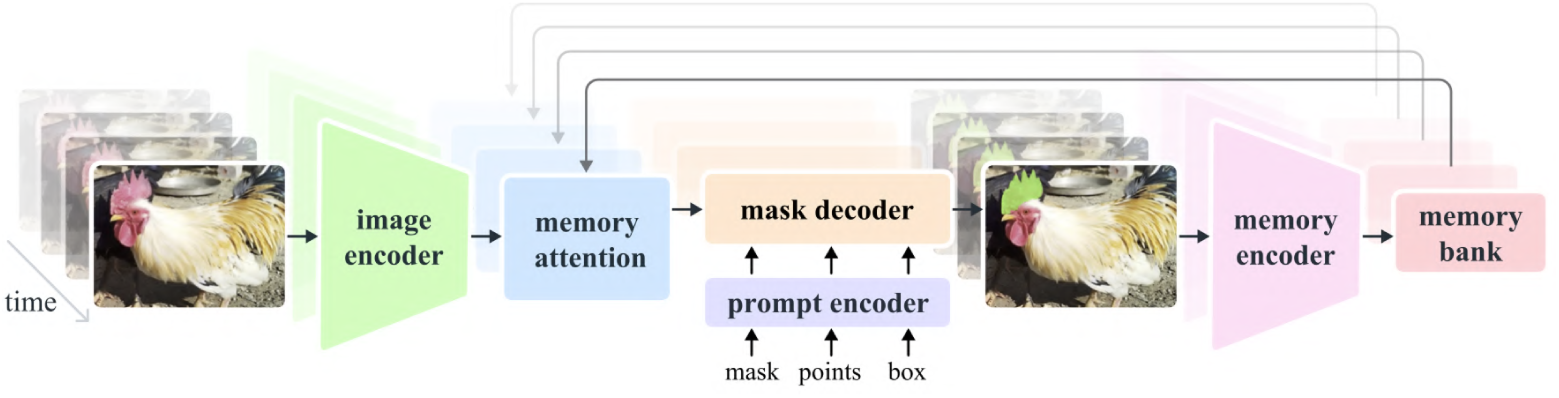
SAM2 architecture overview from [40], accessed 1 April 2025.

#### Implementation details

The SAM experiments were implemented in *python* and were adapted from the public GitHub repository (https://github.com/facebookresearch/segment-anything) [36]. We used the default SAM model with its latest checkpoint. To generate masks in a fully automated fashion, the *SamAutomaticMaskGenerator* class was used. We found that the class parameters in Table 6 generated better results than the default values.

**Table 6:** Configuration parameters for the *SamAutomaticMaskGenerator* class.

| **Parameter** | **Value** |
|---|---|
| points_per_side | 16 |
| pred_iou_thresh | 0.7 |
| stability_score_thresh | 0.85 |
| min_mask_region_area | 100 |
| crop_n_layers | 1 |
| crop_overlap_ratio | 0.6 |

Analogously to the experiments run on SAM, we followed SAM2’s public GitHub repository (https://github.com/facebookresearch/sam2) instructions [40]. This time, we used the *sam2.1_hiera_large* model and its most recent checkpoint. Again, the *SAM2AutomaticMaskGenerator* was selected and set up with the parameter values in Table 7, as it provides fully automatic segmentation.

**Table 7:** Configuration parameters for the *Sam2AutomaticMaskGenerator* class.

| **Parameter** | **Value** |
|---|---|
| points_per_side | 64 |
| pred_iou_thresh | 0.6 |
| stability_score_thresh | 0.85 |
| crop_n_layers | 1 |
| box_nms_thresh | 0.7 |
| crop_n_points_downscale_factor | 3 |
| min_mask_region_area | 100.0 |
| use_m2m | True |

The *generate* class method for both *SAMAutomaticMaskGenerator* and *SAM2AutomaticMaskGenerator* outputs a list of annotation dictionaries, where each annotation is an identified instance in the image. Since these foundation models only segment instances and do not relate them to a class label, it was not reasonable to calculate the previously mentioned performance metrics for the multi-class problem, so only the partition similarity was measured.

### 5.2. U-Net

For this experiment, the goal was to fine-tune a pre-trained deep learning model on our dataset. We chose U-Net as it is foundational to developing more advanced methods, and this network has demonstrated good segmentation results in various medical tasks [13].

U-Net is recognised for its distinctive U-shape architecture, which is detailed in section 2.2.1. For the experiments, ResNet50 [58] was selected as the encoder for U-Net. Figure 6 provides an overview of the network architecture.

**Figure 6:**
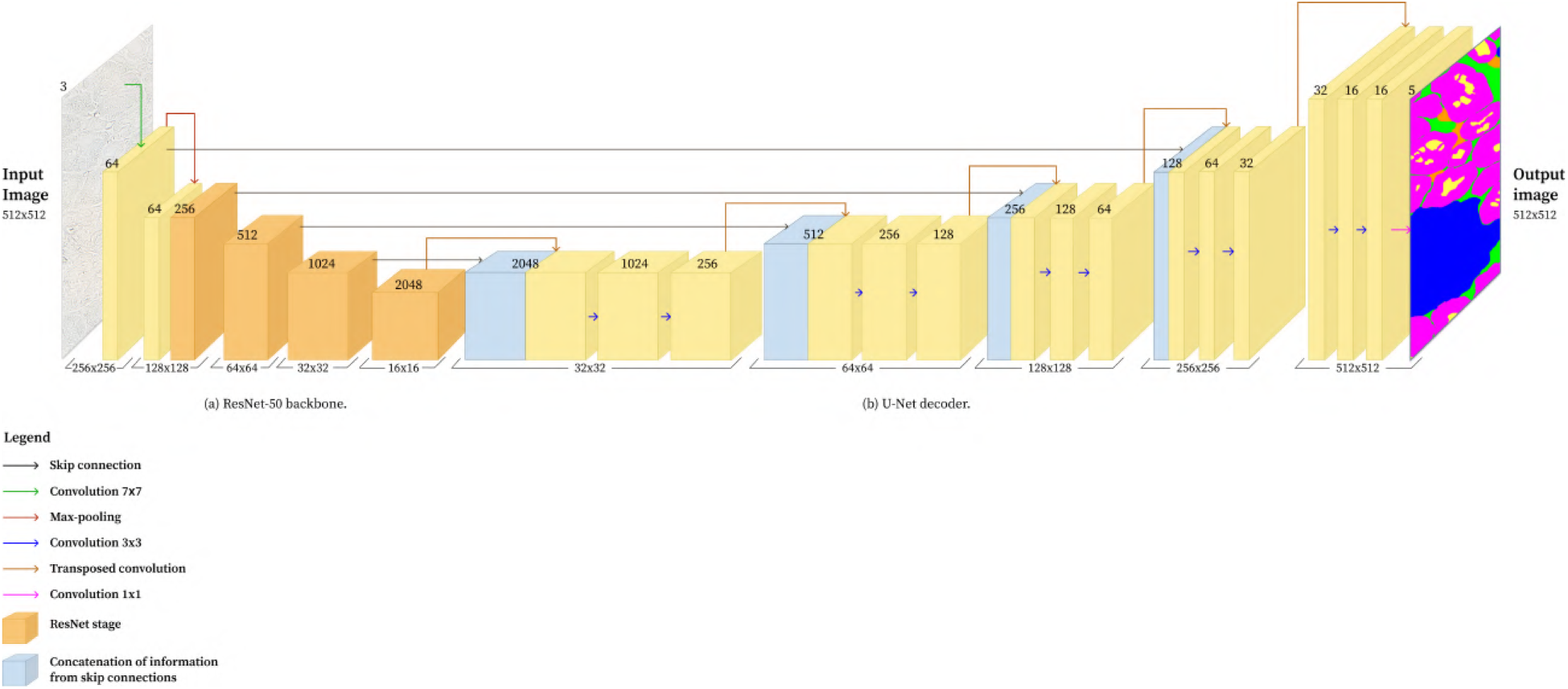
Architecture of U-Net with ResNet-50 backbone. (a) is ResNet-50 backbone architecture; (b) is U-Net decoder architecture. The numbers above each image representation indicate the current number of channels, and the numbers in front correspond to its spatial dimension.

#### Implementation details

These experiments were also implemented in *Python*. The utilised pre-trained U-Net was imported from the *segmentation-models-pytorch Pytorch* package. As mentioned above, we selected the ResNet-50 [58] encoder and the associated weights resultant from pre-training on the ImageNet dataset [59]. A transfer learning approach was adopted such that the encoder’s parameters were frozen, and the decoder was trained on our training set. To improve the model’s robustness for variation in renal structure morphology, a few data augmentation techniques, such as horizontal and vertical flipping, were used.

Our research indicates that a batch size of 6 with the stochastic optimiser Adam [60] and a learning rate of 1 × 10^−4^ produces better results, similar to the Hermsen et al. [14] method. The training occurred over 100 epochs with 91 iterations. The selected loss criterion was Dice Loss, introduced by Sudre et al. as a loss function appropriate for imbalanced class distributions [61]. In our dataset, there was, in fact, a prominent imbalance where the occurrence of a glomerulus or vessels was much rarer than tubules, for instance. We considered the multi-class Dice Loss implementation from the *segmentation-models-pytorch* package:

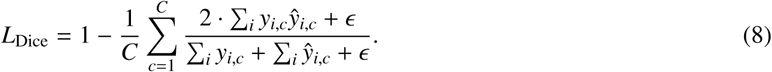

This implementation computes the Dice Loss for a C-class segmentation task using predicted soft probabilities, *y*^, and one-hot encoded ground-truth labels, *y*, where *y_i,c_* = 1 if pixel *i* belongs to class *c*, and zero otherwise.

The DSC is computed per class by calculating the intersection: Σ*_i_ y_i,c_y*^*_i,c_*, and the union approximation: Σ*_i_ y_i,c_* + Σ. *y*^*_i,c_*, where *i* indexes all pixels. Additionally, a small smoothing constant, *ɛ* is added to both the numerator and denominator to avoid division by zero. The per-class scores are averaged into a single scalar value, and the Dice Loss is defined as 1 − Average DSC.

### 5.3. Results

Figure 7 shows two test segmentation masks inferred by the deep learning models and respective manual annotations of renal samples with normal histomorphological appearance.

**Figure 7:**
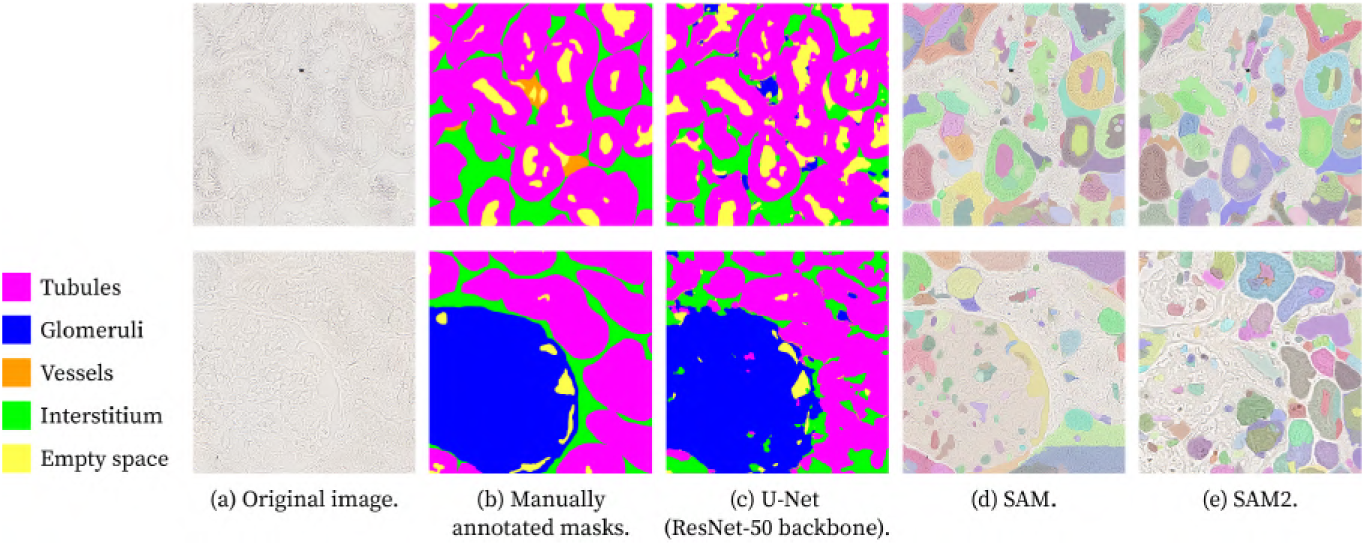
Multi-class segmentation masks output using deep learning models and respective manual annotations, shown for two different example images (top and bottom rows). For each image: (a) Original image, (b) Manually annotated masks, (c) U-Net (ResNet-50 backbone), (d) SAM, and (e) SAM2. *Colour code: tubules (magenta), glomeruli (blue), vessels (orange), interstitium (green), and empty space (yellow)*.

Table 8 reveals that SAM2 is the worst-performing approach from all tested methods. SAM performs surprisingly better than SAM2 but worse than U-Net, with an average partition similarity score of 65.5%.

**Table 8:** Average partition Similarity metric for multi-class segmentation of renal structures.

| **Method** | **Partition Similarity (%)** |
|---|---|
| SAM | 54.1 |
| SAM2 | 44.1 |
| U-Net (ResNet50 encoder) | 65.5 |

Figure 7 clearly shows that U-Net outputs the segmentation masks most similar to the ground-truth annotations. Table 9 shows the performance metric results for this model. Although the accuracy shows high results, it can be justified by its insensitivity to class imbalance, as it does not account for the semantic correctness of the segmented regions. In contrast, metrics such as IoU and DSC fairly evaluate the segmentation by measuring the overlap between predicted and actual object regions within the image, and these results show low performance. Nevertheless, the *tubule* class stands out as the one with better results consistently across all metrics.

**Table 9:** Average performance metrics’ results for multi-class segmentation with U-Net (ResNet-50 backbone).

| **Class** | **IoU** (%) | **DSC** (%) | **Accuracy** (%) | **Precision** (%) | **Recall** (%) |
|---|---|---|---|---|---|
| *Tubule* | 74.2 | 85.0 | 85.6 | 86.0 | 84.6 |
| *Glomerulus* | 56.8 | 67.8 | 91.2 | 67.1 | 71.9 |
| *Vessel* | 1.30 | 2.60 | 97.5 | 13.5 | 1.50 |
| *Interstitium* | 30.9 | 46.2 | 86.0 | 48.5 | 46.2 |
| *Empty space* | 58.2 | 72.6 | 95.0 | 70.1 | 77.1 |

The images in Figure 7, alongside Figure 29, demonstrate that SAM2 struggles to identify the class *glomerulus*. Despite identifying small objects inside the borders of a glomerulus that are not even represented in the manual annotations, it cannot capture the larger object that encapsulates those blobs – the glomeruli.

Across all tested methods for multi-class segmentation, the highest metric scores were achieved for the structure class *tubule*. This result may be due to an imbalanced class distribution, with many instances of this class present in each image. Consequently, the models learned the patterns of these instances and successfully identified them. The structure that models struggled with the most was the *vessel*. For example, Table 9 shows a very weak performance for that class. One could reason that, in contrast to the class *tubule*, there were few objects of the *vessel* class, making it harder for the models to recognise them. Moreover, the manually annotated images in Figure 29 show that vessels are often touching instances of other classes and tend to have smaller sizes than them. This resulted in vessels being frequently classified as instances of different classes, which impacted the respective performance metric values.

### 5.4. Discussion

Foundation segmentation models such as SAM and SAM2 have shown good zero-shot segmentation performance in natural images. However, this preliminary case study reveals that these models are not prepared for zero-shot performance in medical tasks without further fine-tuning on images of this domain. U-Net, a foundational work for image segmentation tasks, exhibited, through this research, that it significantly benefits from fine-tuning. The outputs from these U-Net experiments demonstrate the potential of unstained images, despite their poor contrast. Regardless of these results being considerably better than the outputs of SAM and SAM2, they still show room for improvement. More complex data augmentation and post-processing techniques should be explored to refine the segmentation masks further.

## 6. Advanced techniques for the segmentation of unstained brightfield renal biopsy images

The promising results from our initial study justified further optimisation of U-Net, including the implementation of customised and complex loss functions. In addition to U-Net, we evaluated DeepLabv3+ and TransUNet architectures, which build upon the U-Net framework with more sophisticated design elements. Comparing their performance is essential to determine whether the added architectural complexity translates into meaningful segmentation improvements.

To the best of our knowledge, no prior work has systematically evaluated these advanced models for five-class renal compartment segmentation in unstained brightfield human kidney biopsy images, distinguishing our work from prior glomerulus-focused studies on stained histology and marking a novel contribution to the field.

### 6.1. DeepLabv3+

DeepLab is a family of dilated (atrous) convolutional models. Their primary motivation is to enable kernels to have a larger receptive field without increasing computational cost. This is possible with the introduction of a dilation rate parameter to convolution layers.

DeepLabv3+ is a model that combines cascaded and parallel modules of dilated convolutions and that was proposed by Chen et al. [62]. This architecture builds upon its predecessor, DeepLabv3, by incorporating a decoder module. The resultant architecture resembles the U-Net’s encoder-decoder structure with downsampling and upsampling paths. A key component of this architecture is the Atrous Spatial Pyramid Pooling (ASPP) module. The parallel dilated convolutions with different dilation rates, comprised in ASPP, are concatenated with a 1×1 convolution and global average pooling, as illustrated in Figure 8. This module is an efficient mechanism for capturing multi-scale information without increasing computation complexity.

**Figure 8:**
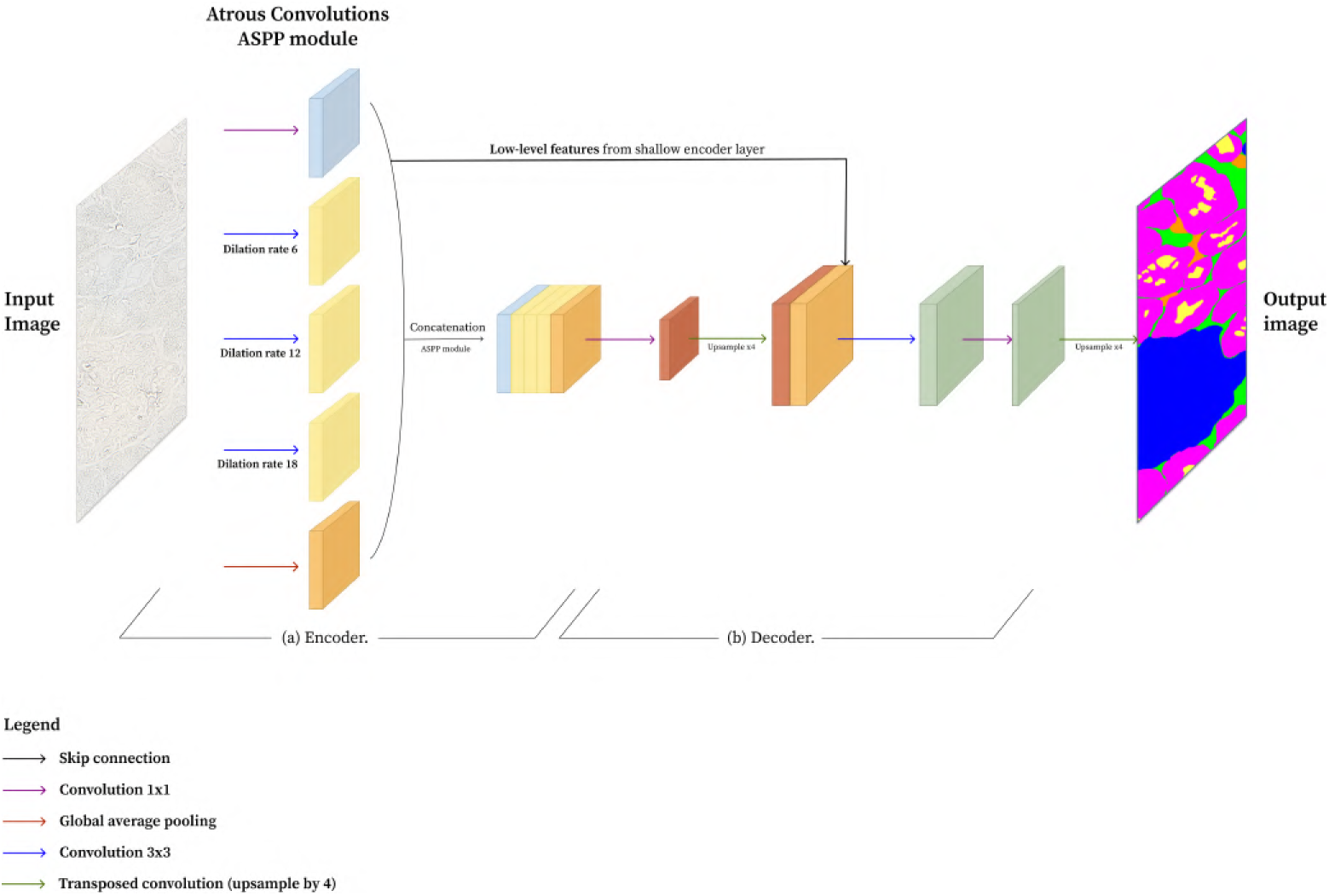
Architecture of DeepLabv3+. (a) Encoder; (b) Decoder. *Adapted from [62]*.

### 6.2. TransUNet

TransUNet is a model that celebrates both the success of U-Net and Transformers, combined to create a strong alternative to U-Net for medical image segmentation [63]. It aims to bypass the limitation of locality, which is inherent to CNN convolutions and may fail to model long-range dependencies. Transformer-based architectures, on the other hand, are designed to support sequence-to-sequence predictions. By replacing the pure CNN encoder with a CNN-Transformer hybrid encoder to extract global context, and by adopting the decoder design of combining upsampled encoded features with high-resolution feature maps to achieve precise localisation, this architecture makes use of the strengths of both Transformers and U-Net.

The TransUNet architecture, depicted in Figure 9, is comprised of twelve Transformer layers composed of Multihead Self-Attention (MSA) and multiple Multi-Layer Perceptron (MLP) blocks. Figure 10 demonstrates the operations that happen in one of these layers.

**Figure 9:**
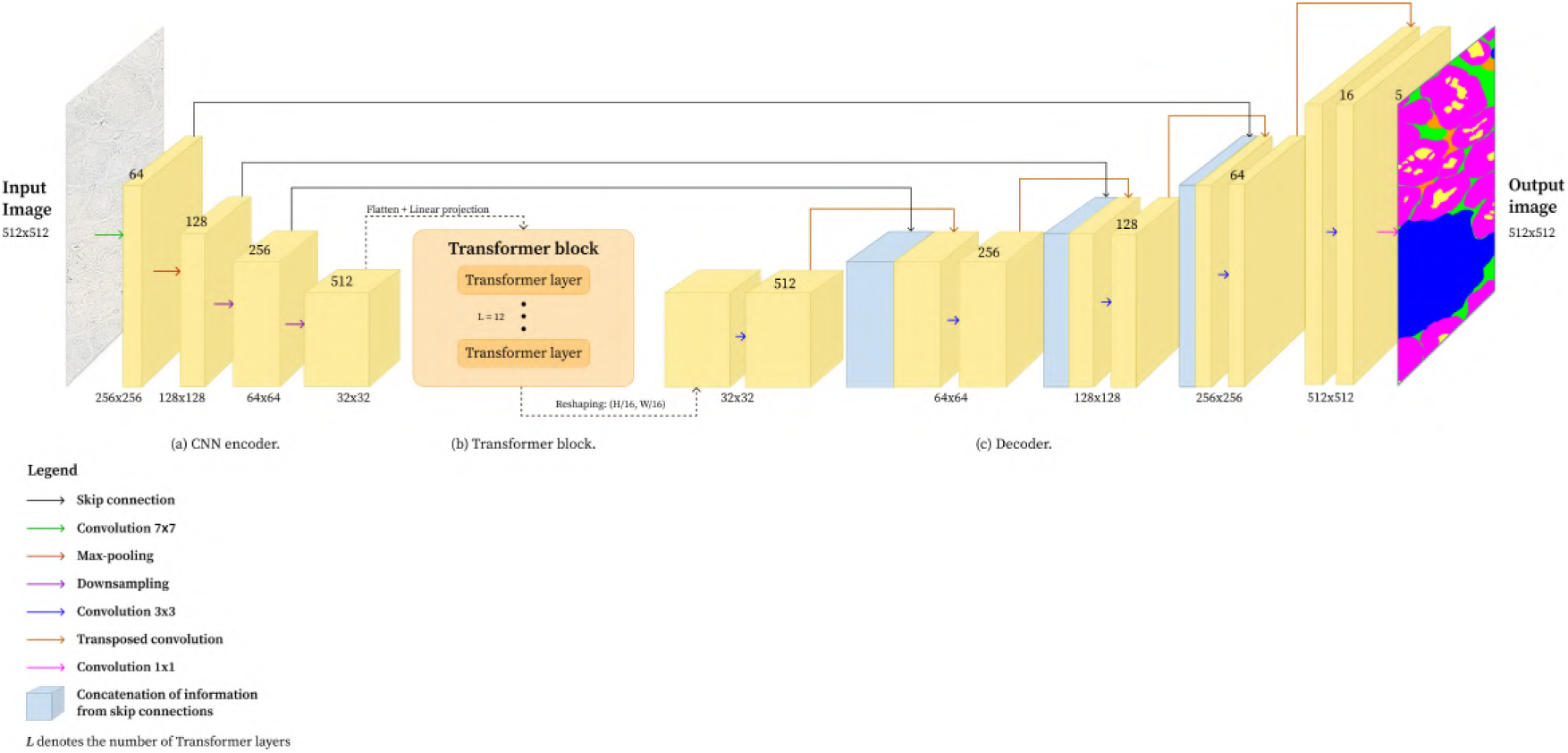
Architecture of TransUNet. (a) CNN encoder; (b) Transformer block; (c) Decoder. The CNN encoder (a) plus the Transformer block (b) compose the full encoder. The numbers above each image representation indicate the current number of channels, and the numbers in front correspond to its spatial dimension. *Adapted from [63]*.

**Figure 10:**
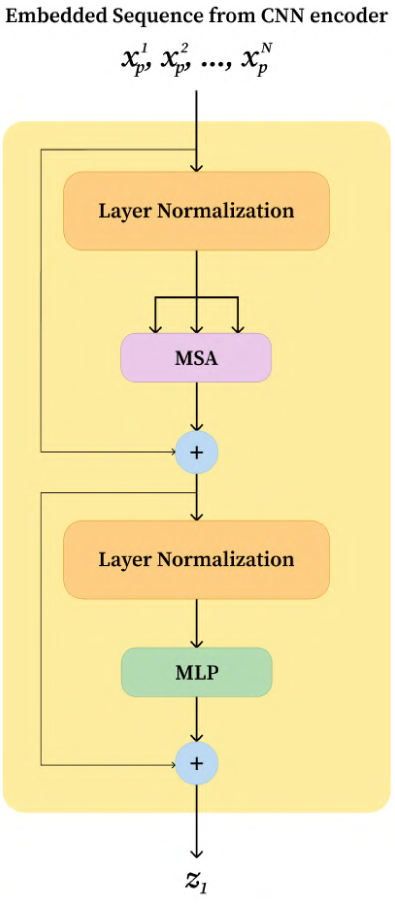
First transformer layer of TransUNet. *Adapted from [63]*.

By integrating transformer-based self-attention with the encoder-decoder architecture of U-Net, TransUNet effectively captures global context but also local spatial details, which are essential for boundary precision.

### 6.3. Loss Function Design

This subsection explores the selected training loss functions. Dice Loss was employed to train the U-Net architecture in the preliminary case study, as it is particularly suitable for imbalanced datasets. Moreover, Dice Loss measures the spatial overlap between prediction and ground-truth masks, making it tailored to segmentation tasks. For this reason, it was considered an essential component of the training objective.

Following the preliminary experiments, hybrid loss functions were explored under the assumption that Dice Loss would remain a contributing factor to the loss, complemented by other terms to improve performance.

#### 6.3.1. Hybrid Loss Functions

Cross-Entropy is a standard loss function for classification tasks. However, it can be effectively integrated with segmentation-tailored losses, such as Dice. In a Dice-Cross-Entropy Loss hybrid, Dice could capture the overlap differences between the prediction and the ground-truth, while Cross-Entropy provides pixel-level control.

For a multi-class problem, Cross-Entropy Loss (CE) becomes Categorical Cross-Entropy Loss, and is defined as:

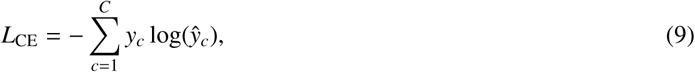

where *C* is the number of classes, *y_c_* is the one-hot encoded ground-truth probabilities for class *c* and *y*^*_c_*the predicted probabilities for class *c*. This loss is computed per pixel and usually averaged across all pixels to produce a scalar loss value.

Focal Loss was introduced by Lin et al. [64] as a version of Cross-Entropy Loss. It was modified to address class imbalance and samples that are hard to classify. It was first proposed for dense object detection tasks, where numerous objects can be overlapped within an image or video; however, it has since been adapted for segmentation purposes. This function penalises more harshly complex or rare pixels, while correctly classified ones contribute less to the total loss.

Focal Loss formulation includes two parameters *α* and *γ*:

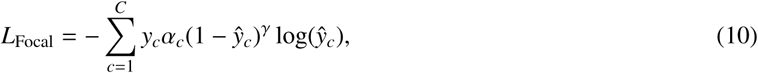

where *y*^*_c_* are the predicted probabilities for class *c*.

Focal and CE loss are mathematically the same if the *γ* parameter is set to zero. Otherwise, Focal Loss will distinguish between confident and wrong predictions, whereas CE will always weigh every pixel loss the same. When a predicted probability *y*^*_c_* is close to 1, the loss loses weight, induced by the (1 − *y*^*_c_*)*^γ^* term. On the contrary, the loss is emphasised if the prediction is very low.

The *α_c_*coefficient is designed to balance class contributions in the loss function, allowing individual weights to be assigned to each class.

The three loss functions discussed – Dice Loss, Cross-Entropy Loss, and Focal Loss – were combined in hybrid loss formulations to guide the model during training and to improve segmentation performance.

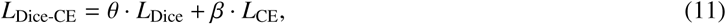

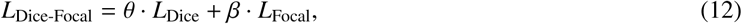

where *θ* and *β* balance the loss terms. Specifically, for the experiments, better results were achieved for the following *θ* and *β* values:

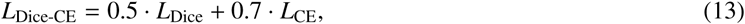

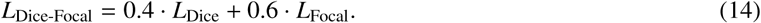

Class-specific weights were applied during loss computation to each loss function. In an experimental phase, the class weights were chosen as the inverse of class frequency throughout the dataset. Before using the weights, they were normalised to avoid extremely small or large weights. In a subsequent experiment, weights were manually chosen, since the previous ones seemed to harm the performance for the most represented classes while gaining minimal improvement for the underrepresented ones.

#### 6.3.2. Boundary Loss

Boundary Loss was introduced to improve segmentation performance, specifically for tasks where precise object boundaries are important. In our study, we adopt the definition established in FixMatchSeg [65]. In that research, Boundary Loss is used as a complement to region-based losses, in particular, Dice Loss. The Boundary Loss function was designed to specifically penalise discrepancies between object contours in predicted and ground-truth masks, which can be overlooked in traditional loss functions.

To compute Boundary Loss following the definition in [65], the first step is to create boundary maps from the predicted and reference maps using morphological operations, which can be implemented via max pooling. Assuming there is a binary ground-truth mask *y_c_* for class *c* and a binary predicted mask *y*^*_c_*, the inverted masks 1 − *y_c_*and 1 − *y*^*_c_*are used to compute the boundaries for the corresponding original masks.

The boundary extraction is performed as follows:

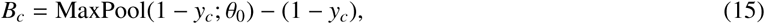

where *θ*_0_ is the kernel size for the initial boundary extraction. The resultant boundary map is dilated with a larger kernel:

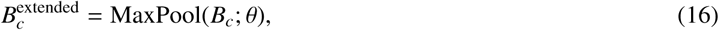

where the parameter *θ* denotes the dilation rate.

The precision and recall for class *c* are:

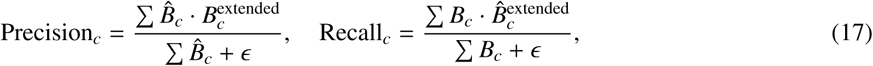

where *ɛ* is a small constant to avoid division by zero.

The Boundary F1-score and respective Boundary Loss can be computed as:

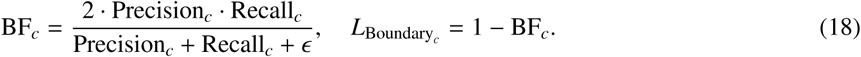

The final reduced loss is the mean over classes:

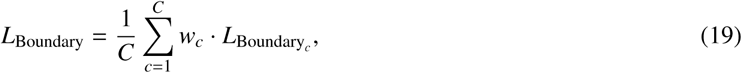

where *w_c_*is a class-specific weight.

Analogously to the methodology proposed in FixMatchSeg research, Boundary Loss can be combined with Dice Loss to create a hybrid loss function that considers region-wise and contour accuracy:

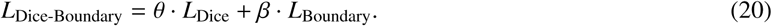

We follow the configuration adopted in [65], where Dice Loss and Boundary Loss have equal loss contribution, *θ* = 0.5 and *β* = 0.5.

#### 6.3.3. Pixel-map for higher frontier penalisation

Building upon the concept of Boundary Loss, we sought to incorporate boundary supervision into additional loss functions. To achieve this, we designed a pixel-wise weight map derived from the ground-truth segmentation masks, structured across C channels corresponding to the number of segmentation classes. In this map, boundary pixels were assigned a weight *w_b_*, while weights for pixels located further from object boundaries decayed exponentially with distance. Integrating this weighted map into the loss computation amplifies the contribution of boundary pixels, thereby directing greater learning focus towards these critical regions and enhancing boundary delineation.

To identify boundary pixels, we applied a 3×3 kernel with a padding of one, excluding the central pixel. This kernel evaluates the immediate neighbourhood of each pixel by counting how many adjacent pixels belong to the same class. A pixel is classified as a boundary pixel if at least one of its neighbours belongs to a different class.

After border pixels are extracted and aggregated across class channels, the distance transform is applied to compute the Euclidean distance from each non-boundary pixel to the closest boundary. The final weight, *w*(*i*) for each pixel *i* is calculated using an exponential decay function:

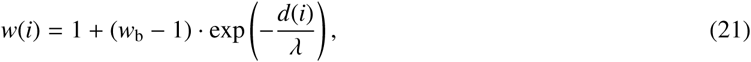

where *w*_b_ is the weight assigned to true boundary pixels, *d*(*i*) is the distance from pixel *i* to the closest boundary, and *λ* is the decay constant that controls how quickly the weight reduces with distance.

Figure 11b illustrates the computed border weight map for the annotated mask in Figure 11a. This heat map visualisation shows the highlighted objects’ frontier, where *w*_b_ is maximal, and the exponential weight decay.

**Figure 11:**
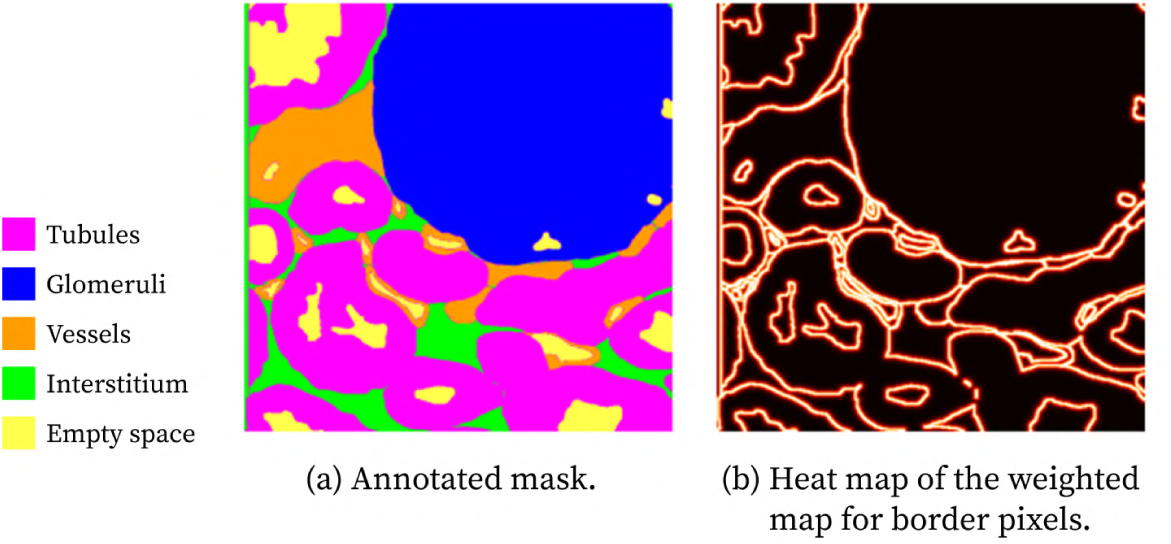
Example of computed weighted border map. Annotation mask (a) and heat map of the computed weighted map for border pixels (b). The heat map, where border pixels with higher weights are brighter, also depicts the exponential weight decay with distance from boundaries. *Colour code: tubules (magenta), glomeruli (blue), vessels (orange), interstitium (green), and empty space (yellow)*.

During experiments, the base weight value that effectively emphasised border pixels without excessively down-weighting interior pixels was *w*_b_ = 3.

This is a simple yet effective approach to employ spatial knowledge in the losses. Additionally, introducing this weighted boundary map into the existing loss functions does not affect their differentiability, which is a requisite for gradient computations and backpropagation. For pixel-wise losses, such as Cross-Entropy Loss and Focal Loss, introducing this weighted map is as simple as:

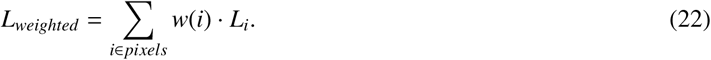

However, to introduce this weighted map into Dice Loss, the loss computation had to be altered slightly:

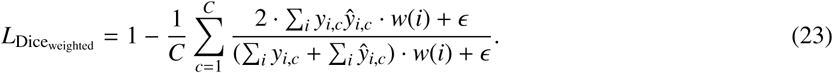

#### 6.3.4. Summary

The customised losses adopted for these advanced experiments are:

- *L*_Dice-CE_ = 0.5 · *L*_Dice_boundary-weights__ + 0.7 · *L*_CE_boundary-weights__
- *L*_Dice-Focal_ = 0.4 · *L*_Dice_boundary-weights__ + 0.6 · *L*_Focal_boundary-weights__
- *L*_Dice-Boundary_ = 0.5 · *L*_Dice_boundary-weights__ + 0.5 · *L*_Boundary_

The design of these hybrid losses is highly relevant to the semantic segmentation objectives of this work. Dice Loss, Cross-Entropy Loss, and Focal Loss each address different challenges in segmentation, such as class imbalance, pixel-wise classification accuracy, and samples that are hard to classify. The renal structures in the brightfield microscopy images demonstrate complex morphology and are spatially close. Therefore, methods that incorporate boundary supervision — such as Boundary Loss or a weighted boundary map — are essential to enhance the model’s ability to delineate object boundaries and achieve accurate segmentation results.

##### Implementation details

These experiments were implemented in a *Python* environment, and conducted on a machine with a NVIDIA GeForce RTX 3060 GPU with 12 GB of VRAM. ResNet-50 was used as the backbone model for every architecture to construct comparable approaches. The pre-trained weights for ResNet-50 from ImageNet were utilised in a transfer learning framework. The U-Net and DeepLabv3+ architectures were imported from the *segmentation-models-pytorch* library, whereas the TransUNet implementation was adapted from the original research GitHub repository (https://github.com/Beckschen/TransUNet).

For every architecture, the encoder’s parameters were first frozen for a 100-epoch training run. Each epoch had 91 iterations and a corresponding validation step. For a second run, the previously frozen encoder parameters were set to update gradients and fine-tune to the renal dataset. For U-Net and DeepLabv3+, the *Pytorch* implementation of stochastic Adam optimiser [60] was initialised with a learning rate of 1 × 10^−3^ for the first run and 1 × 10^−4^ for the second. The optional learning rate decrease upon encoder fine-tuning is to provide a smooth weight transfer from the pre-trained domain to the target domain. Following the TransUNet authors’ original implementation, the *Pytorch* implementation of Stochastic Gradient Descent (SGD) optimiser was used instead, with a learning rate of 0.025, momentum of 0.95, and weight decay of 0.001. To adjust learning rate and help the model escape potential loss plateaus, the *Pytorch ReduceLROnPlateau* method was adopted with a minimum learning rate of 1 × 10^−6^, a patience of 8 epochs, and a reduction factor of 10. It allows for dynamic learning rate reduction based on validation measurements. Consequently, this scheduler was updated after each validation step.

The U-Net architecture was modified to include dropout layers with dropout rates of 0.3 and 0.5 in separate experiments. The performance of the modified version was compared to the performance of the original U-Net architecture. The adopted loss function for this setup was *L*_Dice-CE_. The experimental results of U-Net with dropout didn’t show meaningful improvements from the traditional architecture. Therefore, the original architecture was adopted for the remainder of the work.

The data augmentation mechanisms adopted were Contrast Limited Adaptive Histogram Equalisation (CLAHE) with a probability of 80%, random alterations in blobal brightness and contrast levels within safe interval values and with a probability of 30% per image to avoid undoing the CLAHE enhancement, and square symmetry with a probability of 50%. In subsequent experiments, these mechanisms were incorporated into the training pipelines of U-Net, DeepLabv3+, and TransUNet using the *Albumentations* library. The transformations were dynamically applied to each training data batch.

Besides *L*_Dice-CE_, every model was individually trained with *L*_Dice-Focal_ and *L*_Dice-Boundary_ loss functions. The library https://github.com/BloodAxe/pytorch-toolbeltpytorch-toolbelt was used for both Dice and Focal Loss implementations, whereas *Pytorch* was the source for the utilised Cross-Entropy Loss. In contrast, the Boundary Loss was implemented from scratch.

As mentioned before, some of the ground-truth annotated masks have non-labelled pixels. These pixels were associated with an *ignore label* so that during loss computations, these would not affect the result. Dice, Cross-Entropy, and Focal Loss implementations were designed to support this *ignore label*. To address this specific case in the Boundary Loss custom implementation, masks were created to filter out the pixels that should be ignored.

Each loss function component was reduced by a weighted mean with specific class weights. The weighted map for border pixels was also incorporated into every loss computation, except for the Boundary Loss one.

### 6.4. Results

The following tables and images aggregate the results of these new experiments. The performance metrics’ results were averaged across all the tested samples, and the same image set was utilised to test all model-loss combinations.

Table 10 and Table 11 compare performance results for different custom loss functions using the U-Net architecture with a ResNet-50 backbone. Notably, U-Net trained with a Dice-Cross-Entropy loss hybrid had better prediction power over the other loss functions. Figure 12 illustrates an example of an original tile, the annotated ground-truth, and the U-Net (*L*_Dice-CE_) prediction.

**Figure 12:**
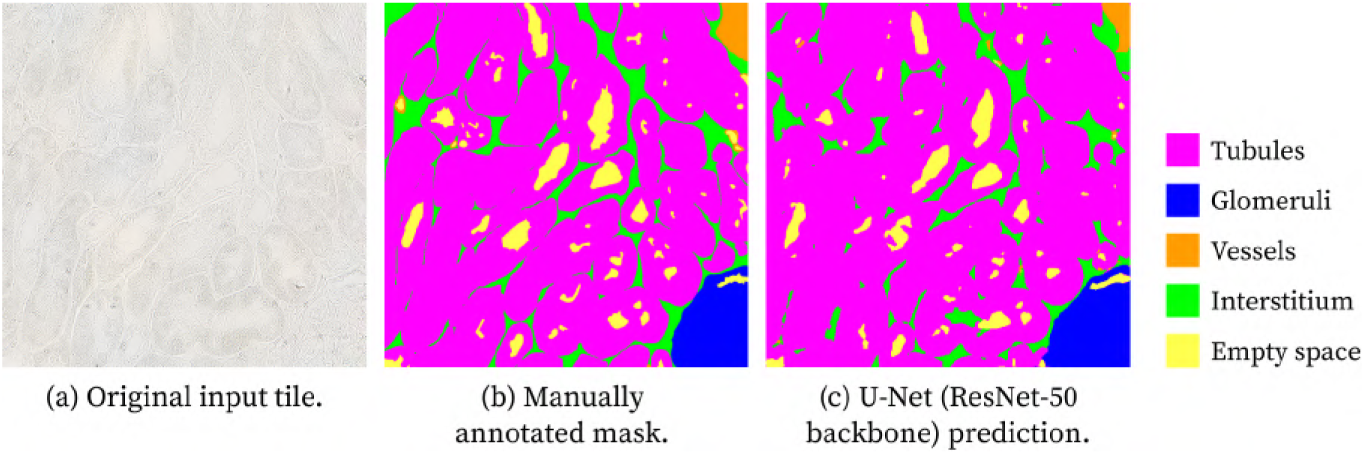
Example of predicted segmentation masks obtained with a **U-Net** (ResNet-50 backbone) trained using the *L*_Dice-CE_ loss function. Panels: (a) Original brightfield image of unstained kidney tissue as input tile, (b) manually annotated mask, and (c) U-Net prediction. *Colour code: tubules (magenta), glomeruli (blue), vessels (orange), interstitium (green), and empty space (yellow)*.

**Table 10:**
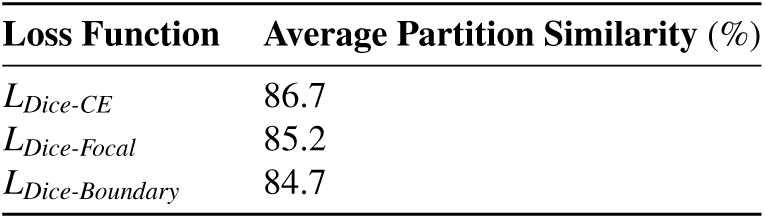
Average Partition Similarity for multi-class segmentation with U-Net (ResNet-50 backbone).

| **Loss Function** | **Average Partition Similarity** (%) |
|---|---|
| *L*<sub>*Dice-CE*</sub> | 86.7 |
| *L*<sub>*Dice-Focal*</sub> | 85.2 |
| *L*<sub>*Dice-Boundary*</sub> | 84.7 |

**Table 11:**
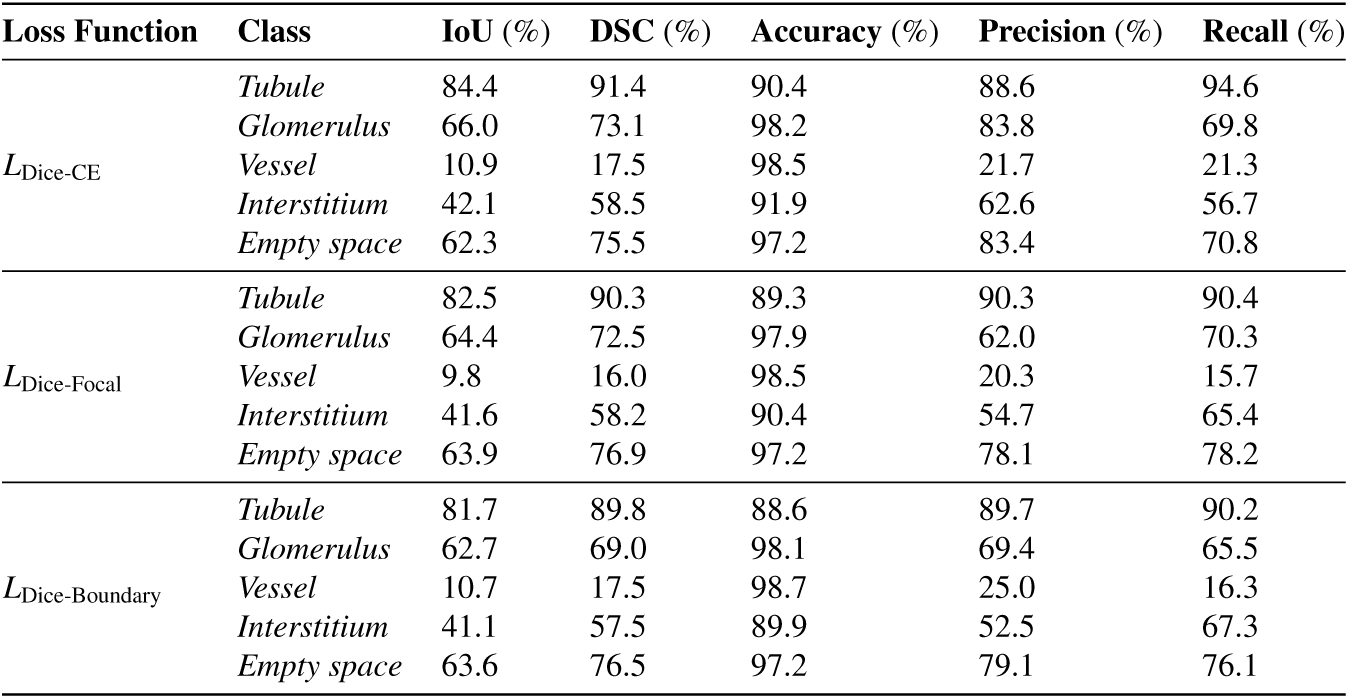
Average performance metrics’ results for multi-class segmentation with U-Net (ResNet-50 backbone) and different loss functions.

| **Loss Function** | **Class** | **IoU** (%) | **DSC** (%) | **Accuracy** (%) | **Precision** (%) | **Recall** (%) |
|---|---|---|---|---|---|---|
| *L*<sub>Dice-CE</sub> | *Tubule* | 84.4 | 91.4 | 90.4 | 88.6 | 94.6 |
|  | *Glomerulus* | 66.0 | 73.1 | 98.2 | 83.8 | 69.8 |
|  | *Vessel* | 10.9 | 17.5 | 98.5 | 21.7 | 21.3 |
|  | *Interstitium* | 42.1 | 58.5 | 91.9 | 62.6 | 56.7 |
|  | *Empty space* | 62.3 | 75.5 | 97.2 | 83.4 | 70.8 |
| *L*<sub>Dice-Focal</sub> | *Tubule* | 82.5 | 90.3 | 89.3 | 90.3 | 90.4 |
|  | *Glomerulus* | 64.4 | 72.5 | 97.9 | 62.0 | 70.3 |
|  | *Vessel* | 9.8 | 16.0 | 98.5 | 20.3 | 15.7 |
|  | *Interstitium* | 41.6 | 58.2 | 90.4 | 54.7 | 65.4 |
|  | *Empty space* | 63.9 | 76.9 | 97.2 | 78.1 | 78.2 |
| *L*<sub>Dice-Boundary</sub> | *Tubule* | 81.7 | 89.8 | 88.6 | 89.7 | 90.2 |
|  | *Glomerulus* | 62.7 | 69.0 | 98.1 | 69.4 | 65.5 |
|  | *Vessel* | 10.7 | 17.5 | 98.7 | 25.0 | 16.3 |
|  | *Interstitium* | 41.1 | 57.5 | 89.9 | 52.5 | 67.3 |
|  | *Empty space* | 63.6 | 76.5 | 97.2 | 79.1 | 76.1 |

Table 12 and Table 13 compare performance results for different custom loss functions using the DeepLabv3+ architecture with a ResNet-50 backbone. In this case, the training loss function that produced higher segmentation performance was the hybrid Dice-Focal Loss. Figure 13 illustrates an example of an original tile, the annotated ground-truth, and the DeepLabv3+ (*L*_Dice-Focal_) prediction.

**Figure 13:**
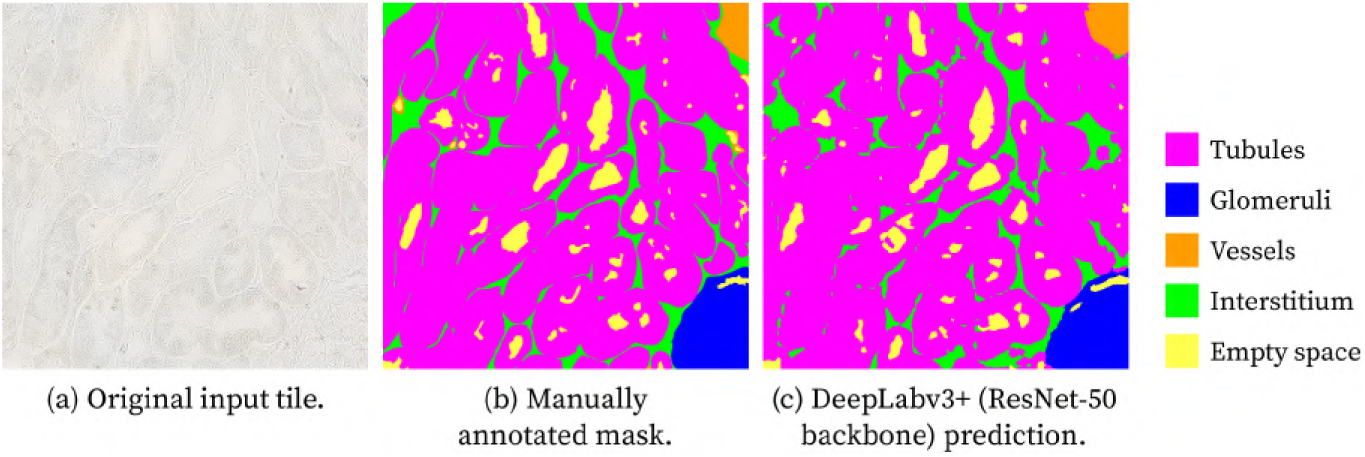
Example of predicted segmentation masks obtained with a **DeepLabv3**+ (ResNet-50 backbone) trained using the *L*_Dice-Focal_ loss function. Panels: (a) Original brightfield image of unstained kidney tissue as input tile, (b) manually annotated mask, and (c) U-Net prediction. *Colour code: tubules (magenta), glomeruli (blue), vessels (orange), interstitium (green), and empty space (yellow)*.

**Table 12:**
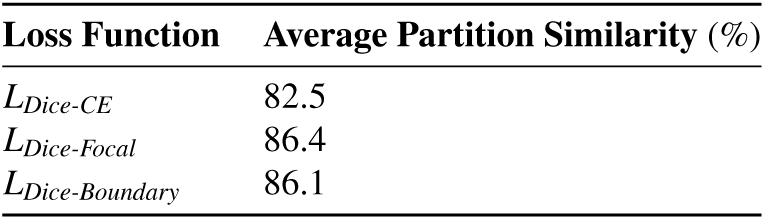
Average Partition Similarity for multi-class segmentation with DeepLabv3+ (ResNet-50 backbone).

| **Loss Function** | **Average Partition Similarity** (%) |
|---|---|
| *L*<sub>*Dice-CE*</sub> | 82.5 |
| *L*<sub>*Dice-Focal*</sub> | 86.4 |
| *L*<sub>*Dice-Boundary*</sub> | 86.1 |

**Table 13:** Average performance metrics’ results for multi-class segmentation with DeepLabv3+ with different loss functions.

| **Loss Function** | **Class** | **IoU** (%) | **DSC** (%) | **Accuracy** (%) | **Precision** (%) | **Recall** (%) |
|---|---|---|---|---|---|---|
| *L*<sub>Dice-CE</sub> | *Tubule* | 79.4 | 88.3 | 86.8 | 89.8 | 87.2 |
|  | *Glomerulus* | 63.9 | 71.3 | 98.0 | 82.2 | 68.9 |
|  | *Vessel* | 10.7 | 16.6 | 98.4 | 17.2 | 20.3 |
|  | *Interstitium* | 38.3 | 54.5 | 88.4 | 47.5 | 67.7 |
|  | *Empty space* | 59.4 | 73.2 | 96.4 | 83.3 | 68.2 |
| *L*<sub>Dice-Focal</sub> | *Tubule* | 84.3 | 91.4 | 90.1 | 91.3 | 91.6 |
|  | *Glomerulus* | 69.5 | 76.2 | 98.3 | 76.2 | 74.9 |
|  | *Vessel* | 11.3 | 17.6 | 98.5 | 20.6 | 20.2 |
|  | *Interstitium* | 42.6 | 59.0 | 91.1 | 57.1 | 63.4 |
|  | *Empty space* | 64.1 | 77.1 | 97.2 | 79.1 | 76.7 |
| *L*<sub>Dice-Boundary</sub> | *Tubule* | 83.2 | 90.7 | 89.4 | 89.7 | 91.9 |
|  | *Glomerulus* | 70.4 | 76.7 | 98.4 | 72.7 | 75.8 |
|  | *Vessel* | 11.7 | 18.2 | 98.6 | 24.0 | 21.7 |
|  | *Interstitium* | 41.2 | 57.5 | 91.0 | 57.2 | 60.6 |
|  | *Empty space* | 62.9 | 75.9 | 97.2 | 79.1 | 74.6 |

Table 14 and Table 15 compare performance results for different custom loss functions using the TransUNet architecture with a ResNet-50 CNN encoder. Despite the *L*_Dice-Focal_ setup reaching performance scores slightly higher in Partition Similarity, and for the prediction of pixels of the *tubule* class, the overall higher performance was achieved by the custom loss function *L*_Dice-CE_, similarly to U-Net. Figure 14 illustrates an example of an original tile, the annotated ground-truth, and the TransUNet (*L*_Dice-CE_) prediction.

**Figure 14:**
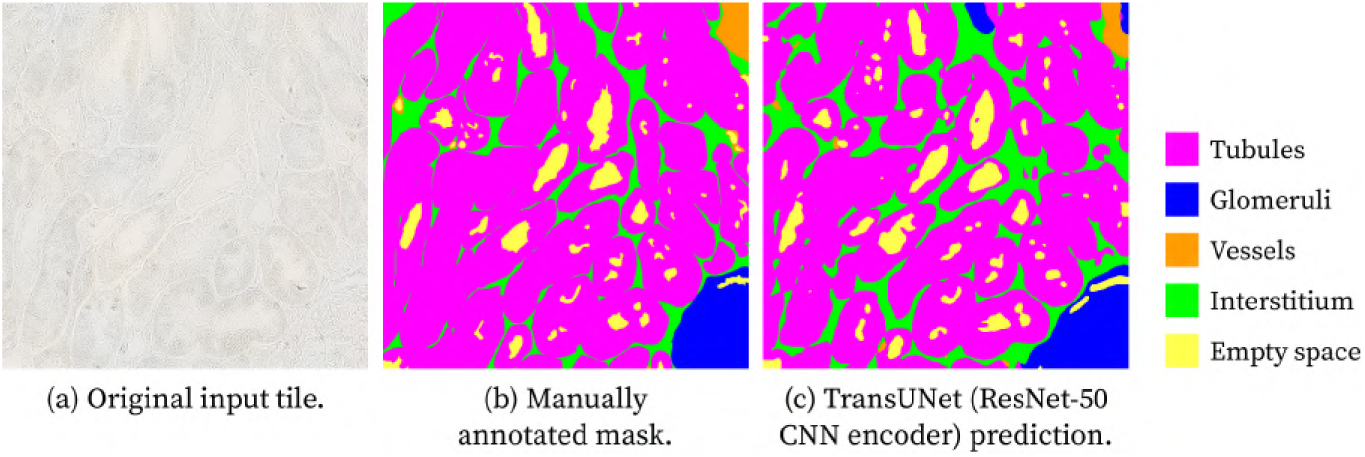
Example of predicted segmentation masks obtained with a **TransUNet** (ResNet-50 CNN encoder) trained using the *L*_Dice-CE_ loss function. Panels: (a) Original brightfield image of unstained kidney tissue as input tile, (b) manually annotated mask, and (c) U-Net prediction. *Colour code: tubules (magenta), glomeruli (blue), vessels (orange), interstitium (green), and empty space (yellow)*.

**Table 14:**
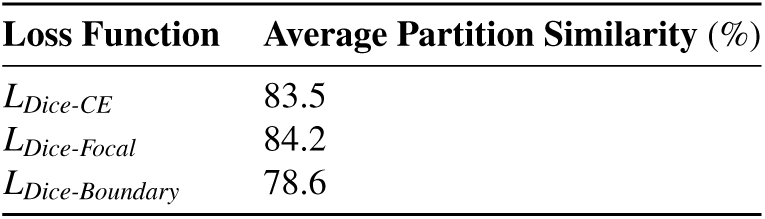
Average Partition Similarity for multi-class segmentation with TransUNet (ResNet-50 CNN encoder).

| **Loss Function** | **Average Partition Similarity** (%) |
|---|---|
| *L*<sub>*Dice-CE*</sub> | 83.5 |
| *L*<sub>*Dice-Focal*</sub> | 84.2 |
| *L*<sub>*Dice-Boundary*</sub> | 78.6 |

**Table 15:** Average performance metrics’ results for multi-class segmentation with TransUNet with different loss functions.

| **Loss Function** | **Class** | **IoU** (%) | **DSC** (%) | **Accuracy** (%) | **Precision** (%) | **Recall** (%) |
|---|---|---|---|---|---|---|
| *L*<sub>Dice-CE</sub> | *Tubule* | 80.6 | 89.2 | 88.1 | 94.9 | 84.3 |
|  | *Glomerulus* | 72.5 | 79.9 | 98.4 | 71.7 | 78.6 |
|  | *Vessel* | 10.0 | 16.6 | 98.0 | 16.0 | 24.5 |
|  | *Interstitium* | 40.5 | 56.9 | 88.6 | 47.9 | 74.1 |
|  | *Empty space* | 62.5 | 76.0 | 96.9 | 70.7 | 84.9 |
| *L*<sub>Dice-Focal</sub> | *Tubule* | 81.5 | 89.7 | 88.8 | 91.7 | 88.2 |
|  | *Glomerulus* | 69.8 | 77.3 | 98.2 | 67.4 | 75.5 |
|  | *Vessel* | 6.9 | 11.3 | 98.3 | 14.6 | 13.2 |
|  | *Interstitium* | 37.4 | 53.6 | 89.7 | 50.7 | 59.8 |
|  | *Empty space* | 60.3 | 74.0 | 96.7 | 71.2 | 80.5 |
| *L*<sub>Dice-Boundary</sub> | *Tubule* | 75.7 | 86.0 | 84.4 | 88.3 | 84.3 |
|  | *Glomerulus* | 55.5 | 66.5 | 94.6 | 49.9 | 75.2 |
|  | *Vessel* | 5.9 | 10.8 | 98.1 | 11.5 | 20.6 |
|  | *Interstitium* | 26.2 | 40.9 | 87.1 | 41.2 | 44.2 |
|  | *Empty space* | 59.3 | 73.2 | 96.7 | 71.5 | 78.9 |

### 6.5. Discussion

The best-performing loss function varied depending on the model’s architecture. For U-Net and TransUNet, the Dice-Cross-Entropy hybrid was the preferred loss function. This is a standard loss function combination, which ha also been proven effective for this dissertation’s research. In contrast, the Dice-Focal loss function worked best for DeepLabv3+. The atrous convolutions incorporated in this architecture, and the captured multi-scale context could be dilating the spatial detail perceived. The Focal Loss component could effectively weigh the loss of harder-to-classify pixels, which impacted the training and resulted in increased segmentation performance across all metrics.

The *glomerulus* and *vessel* classes demonstrate high standard deviation values from the average performance. Due to the heavy imbalance of class distributions, the objects from these classes were harder to identify.

Additionally, vessels are morphologically heterogeneous. For instance, capillaries are very small structures, comparatively smaller than the other types of structures, whereas arteries stand out due to their large size, as depicted in Figure 15. These aspects altogether compromised the learning for this class, which justifies the poor segmentation performance.

**Figure 15:**
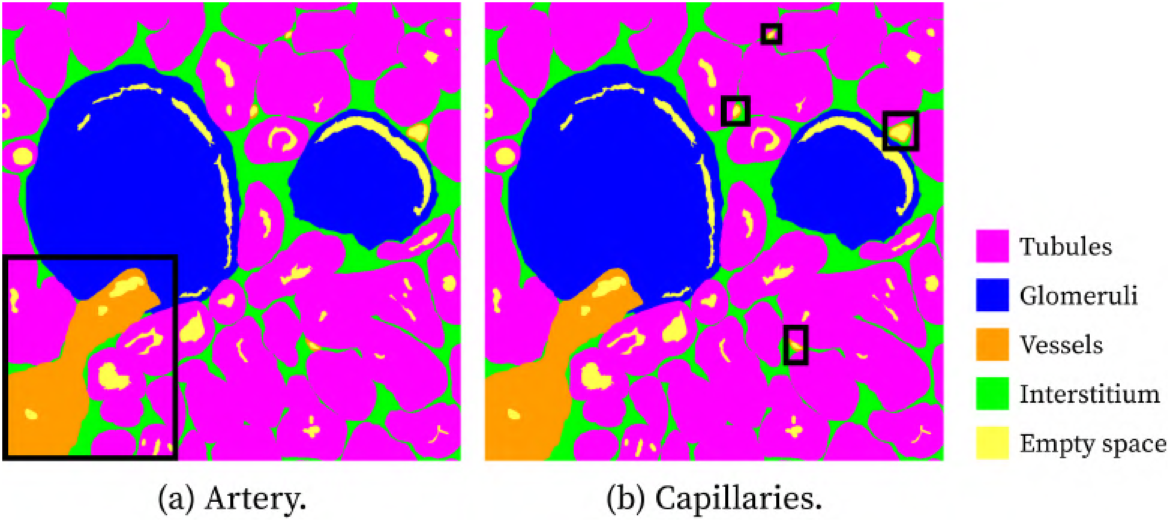
Examples of manually annotated vessel segmentations from the dataset. (a) Artery and (b) capillaries are indicated within the masks. *Colour code: tubules (magenta), glomeruli (blue), vessels (orange), interstitium (green), and empty space (yellow)*.

By analysing confusion matrices relative to these experiments, we gathered that *vessels* are often mistaken for *interstitium*. With this insight, one possibility would be to merge these two classes and have the model predict the pixels for a class “*vessel* + *interstitium*”. Upon using the inferred segmentation masks from this model, this merged class would require an expert in renal structures to distinguish the instances from both structural classes.

The Dice-Boundary Loss Function proved to be less effective compared to Dice-Cross-Entropy and Dice-Focal Loss. This shows that the loss customisation by incorporating the weighted map for the boundary pixels was more effective than the use of a Boundary Loss component.

The three best-performing configurations for each model are as follows:

- U-Net with *L*_Dice-CE_;
- DeepLabv3+ with *L*_Dice-Focal_;
- TransUNet with *L*_Dice-CE_;

These models exhibit comparable performance, making it difficult to determine a clear winner. Figure 16 presents three representative examples of segmentation outputs from U-Net, DeepLabv3+, and TransUNet. For additional segmentation results, refer to Figure 30 in Appendix A.

**Figure 16:**
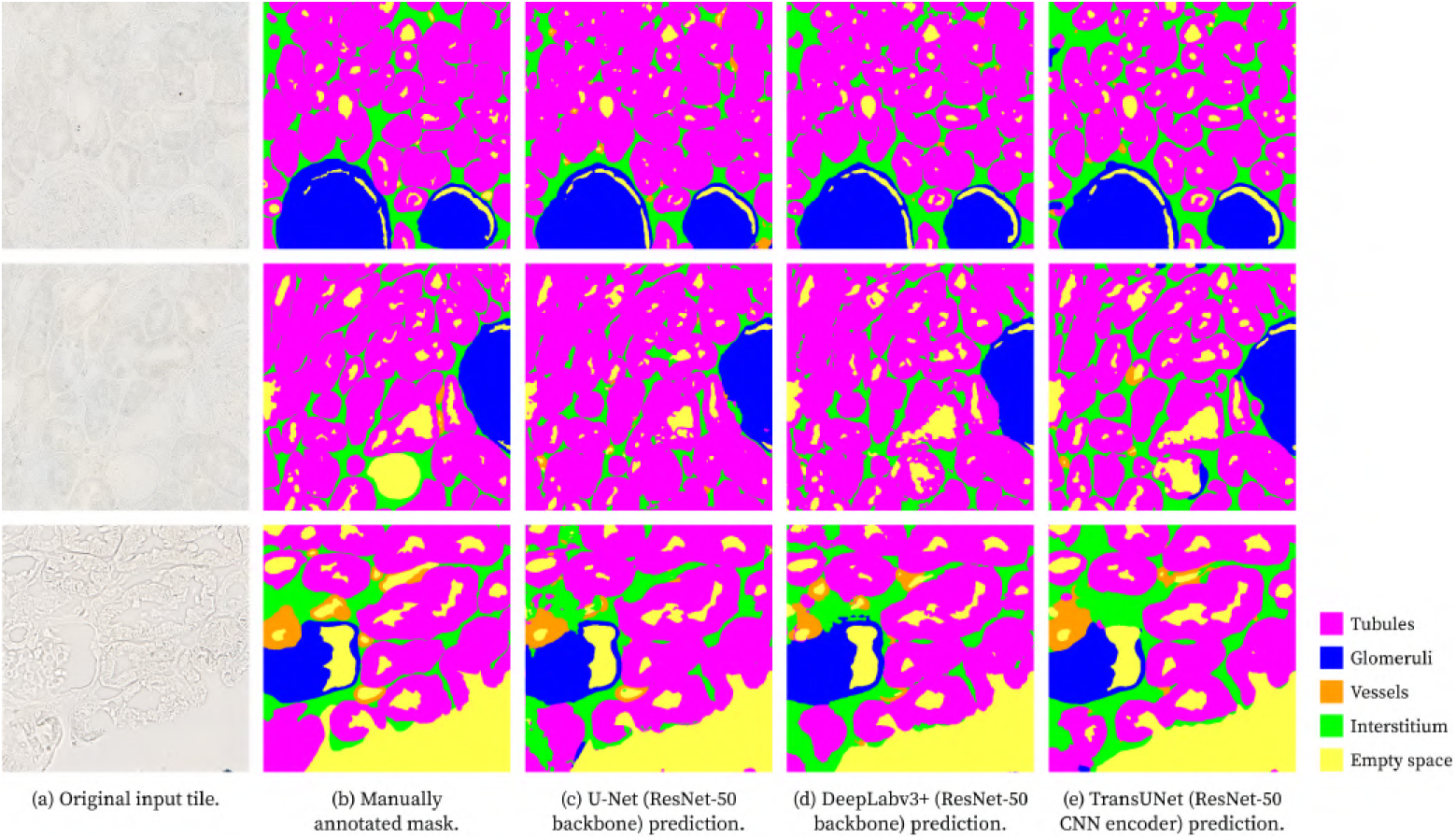
Multi-class segmentation masks output using the supervised deep learning models and respective manual annotations, shown for three different example images (top, middle, and bottom rows). For each image: (a) Original image, (b) Manually annotated masks, (c) U-Net (*L*_Dice-CE_), (d) DeepLabv3+ (*L*_Dice-Focal_), and (e) TransUNet (*L*_Dice-CE_). *Colour code: tubules (magenta), glomeruli (blue), vessels (orange), interstitium (green), and empty space (yellow)*.

All the models’ predictions are visually very similar to the ground-truth masks. The main noticeable limitations are the high misclassification rate for the *vessel* class and object boundary delineation, which could be improved. Nonetheless, the TransUNet predictions Figure 16e show an increased instance separation compared to the other models’ masks. However, that does not precisely align with the true annotations, as the predicted object boundaries seem more eroded than supposed. This resulted in misclassified pixels, or in some cases, entire instances were undersegmented.

Through visual comparison, it is almost impossible to determine which of DeepLabv3+’s and U-Net’s segmentation masks are closest to the true annotations. There are pixel regions where DeepLabv3+ predictions were more accurate, and others where U-Net predicted better. As a result, the best-performing model changes per image.

Figure 17 demonstrates a possible application of the segmentation masks produced by these models. The edges present in Figure 17c were extracted with the same algorithm used for the creation of the weighted boundary map without assigning weights to pixels, and by outputting the boundary map itself.

**Figure 17:**
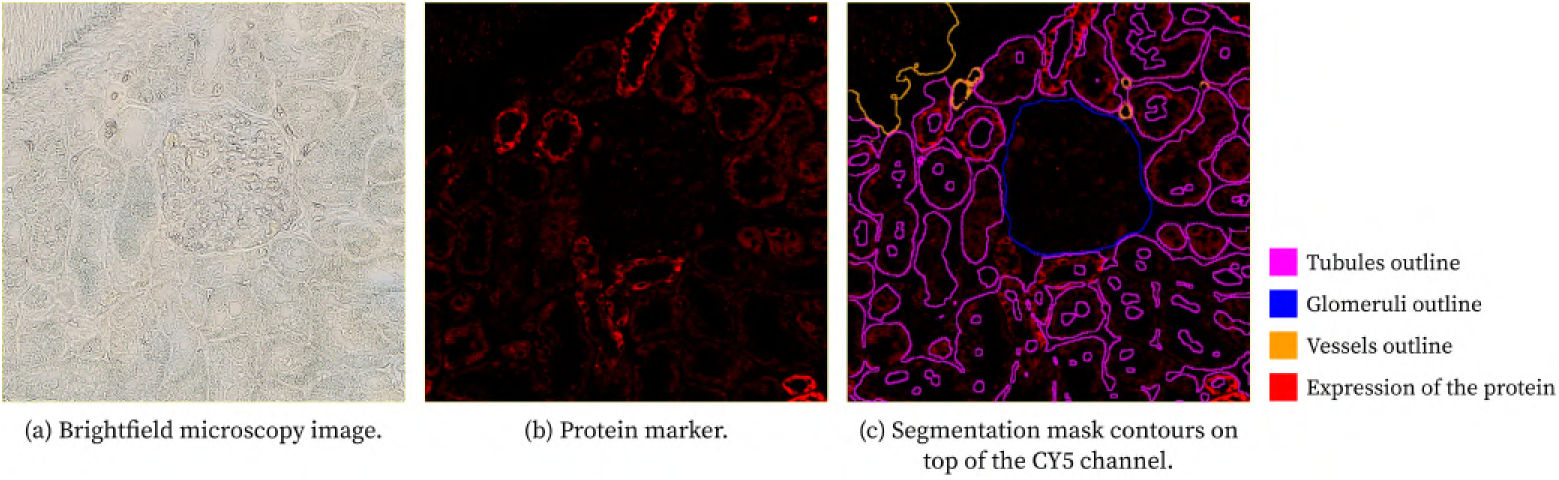
Application of segmentation mask generated by U-Net to unstained brightfield kidney tissue image for the compartment-specific analysis of an immunofluorescent-labelled protein. Brightfield (a) and corresponding fluorescence image of protein-labelling (b) and U-Net segmentation mask (c) for the different kidney tissue compartments. The secondary antibody used for the immunofluorescence staining was Donkey Anti-Rabbit IgG (H+L) Highly Cross-Adsorbed Secondary Antibody, Alexa Fluor™ Plus 647 (Invitrogen, Thermo Fisher Scientific; Waltham, MA, USA; catalog #A32795). The brightfield and fluorescence images were acquired in a NanoZoomer S60 Digital Slide Scanner (Hamamatsu Photonics; Hamamatsu, Japan) (a, b). *Colour code: tubules outline (magenta), glomeruli outline (blue), vessels outline (orange), immunofluorescent-labelled protein (red)*.

### 6.6. Preliminary study of samples that include structures with histomorphological alterations

From the results obtained in the above analysis, it is clear that with further refinement and training, the tested models show promising performance in this elementary segmentation task, where the images show only renal structures with normal histomorphological appearance. However, it is important for clinical applications that the networks can also perform effectively on images containing structures with histomorphological alterations, aiding pathologists in diagnosing.

Consequently, as a preliminary approach, it was relevant to test this model on example tiles that present structural alterations such as glomerulosclerosis, tubular atrophy, and interstitial fibrosis. As opposed to the five classes present in the images used in the previous experiments, the diseased samples have eight different structures: glomeruli, tubules, vessels and interstitium that have a normal histomorphological appearance, as well as fibrotic tubules, fibrotic tubulointerstitium, sclerotic glomeruli, and the empty space in the interior of structures.

As Figure 18 shows, this model’s segmentation of samples with histomorphological alterations obtained the expected results, where the structures with normal histomorphological appearance are mostly correctly identified.

**Figure 18:**
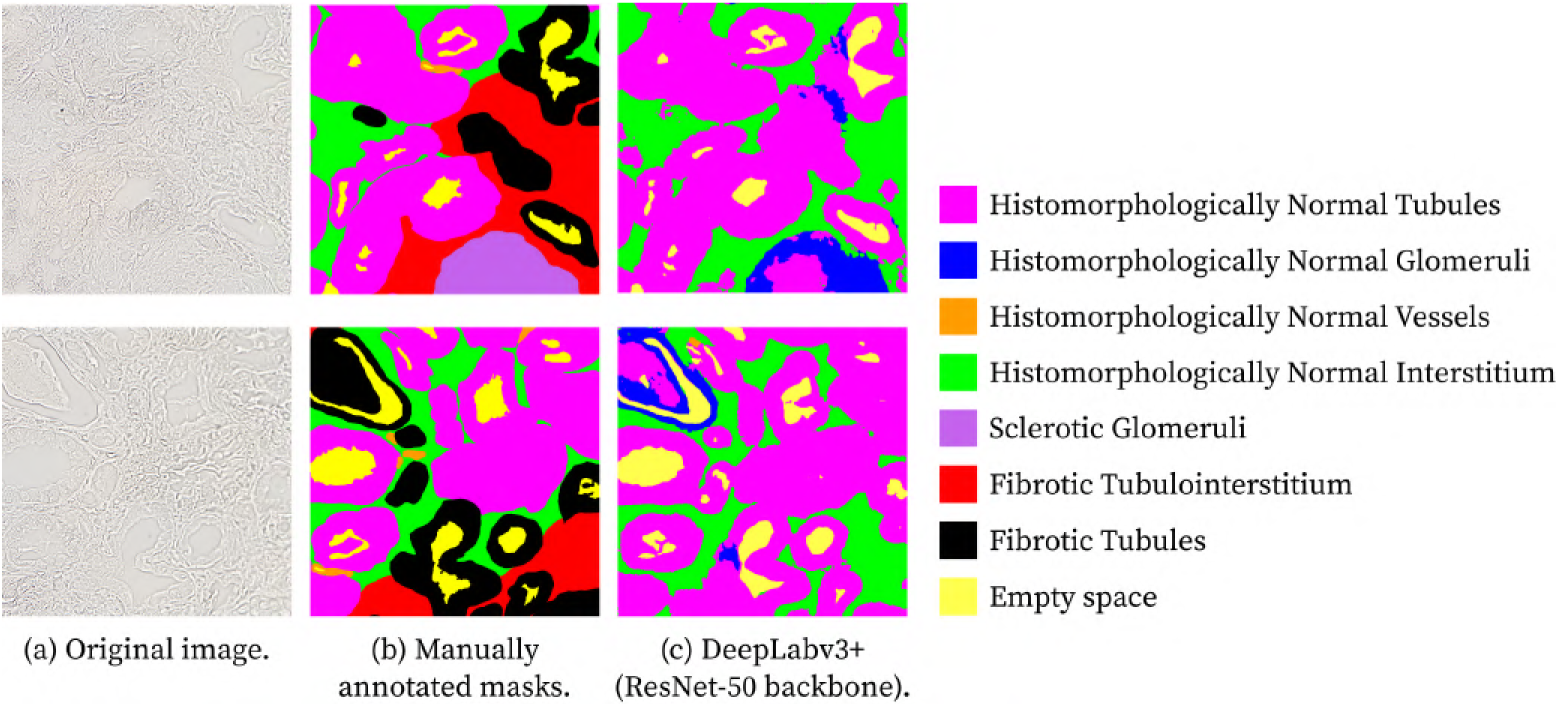
Multi-class segmentation masks output performed on brightfield human kidney tissue presenting histomorphological alterations – glomerulosclerosis, fibrotic tubules, and interstitial fibrosis –, shown for two different example images (top and bottom rows). Brightfield image of human kidney tissue with histomorphological alterations (a) and respective manual (b) and DeepLabv3+ (c) annotations. *Colour code: histomorphologically normal tubules (magenta), histomorphologically normal glomeruli (blue), histomorphologically normal vessels (orange), histomorphologically normal interstitium (green), sclerotic glomeruli (purple), fibrotic tubulointerstitium (red), fibrotic tubules, and empty space (yellow)*.

The fibrotic tubules are often identified as tubules, which is a significant achievement, since with disease, these structures may show changes in their shape. However, it would be relevant to distinguish the instances that have alterations from the ones with a normal appearance.

Future work is motivated to experiment with Open-set Classification methodologies [66] and classify these diseased instances as unknown or include these samples in the network’s training.

The following experiments focus on semi-supervised learning with FixMatchSeg. In the original study [65], a U-Net architecture is trained and supplemented with pseudo-label generation and consistency regularisation. In accordance with the original method, the next section explores the implementation of FixMatchSeg using a U-Net architecture.

## 7. Semi-supervised semantic segmentation

One of the most significant limitations of Computer Vision tasks, especially with biomedical images, is the scarcity of available data, as biomedical images are hard to acquire. For instance, microscopy images of human renal tissue, such as the images used in this research, are typically submitted to extensive steps for image acquisition. First, a biopsy is performed to extract a tissue portion, then the tissue must undergo preparation steps for microscopy, and lastly, the prepared tissue slide must be digitised. All these steps make for a costly and time-consuming process. Another crucial aspect is that medical data must be anonymous, and as a result, most of it is not made publicly available before careful anonymisation. In addition, there is a lack of annotations of the gathered data. Without ground-truth references, a supervised deep learning model is not able to learn how to accurately represent the data. The images used in this work are characterised by high structural complexity and low contrast. In return, annotating these images, which must be done precisely, is extremely time-consuming, and therefore, there is not much of it.

Unsupervised learning is a machine learning framework where models learn from unlabelled data. Since there is no outside guidance to help the model converge and minimise learning error, the model must identify hidden patterns to create representations of the given data. Semi-supervised learning is a middle ground that combines supervised and unsupervised learning. Usually, a small portion of the data, the labelled part, is trained with supervision, while the remaining portion is trained without supervision. This methodology allows for the model to train with larger amounts of data while retaining essential ground-truth context from the supervised component.

The following section delineates the implementation of semi-supervised learning for the semantic segmentation of unstained renal brightfield microscopy images using FixMatchSeg.

### 7.1. FixMatchSeg

FixMatchSeg was introduced by Upretee and Khanal as a semi-supervised approach for semantic segmentation [65]. It adapts the idea of FixMatch [67], originally designed for image classification tasks. FixMatchSeg, depicted in Figure 19, adopts two semi-supervised techniques: consistency regularisation and pseudo-labelling.

**Figure 19:**
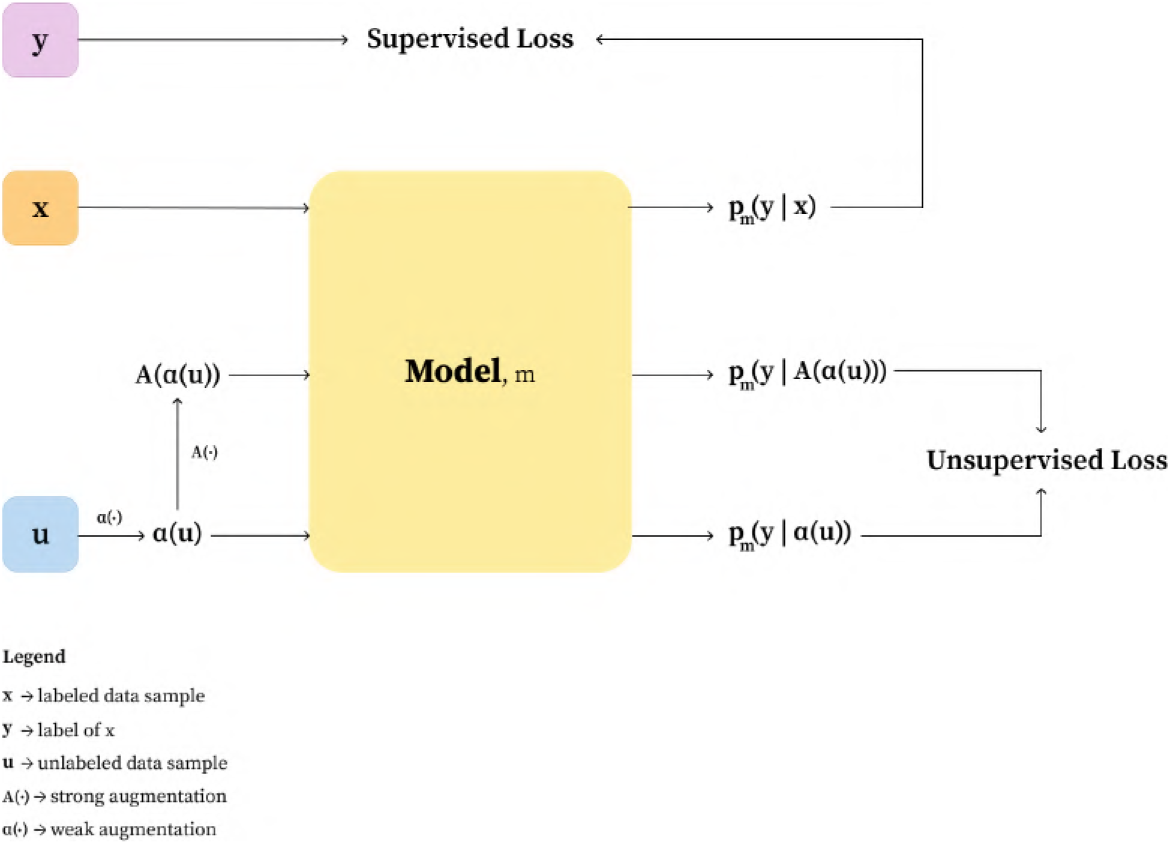
FixMatch training methodology. *Adapted from [65]*.

Consistency regularisation is addressed by using weak and strong data augmentations, such that “for two nearby points in the input space, their labels must be the same. In other words, a robust model should obtain similar predictions for both a point and a locally modified version of it” [68]. Consistency regularisation methods are grounded in the smoothness assumption, which posits that a robust model should produce stable predictions when small perturbations are applied to unlabelled data. These methods enforce prediction consistency by introducing controlled alterations to the input and training the model to remain invariant to such changes. However, FixMatchSeg addresses consistency regularisation without the complex teacher-student network, by assuming two kinds of data augmentation: strong and weak augmentations. The weak augmentations, denoted by *α*, are basic geometric transformations, such as random rotation and elastic distortions. In contrast, strong augmentations, denoted by *A*, should maintain the geometry of the objects in the images and only alter contrast, sharpness, and blurring. The goal is that the strong augmentations are applied to previously weakly augmented images, so that two versions of the same image are obtained by applying perturbations, as can be observed in Figure 20. The unsupervised loss in this approach is therefore calculated between predictions *f_θ_*(*A*(*α*(*x*))) and *f_θ_*(*α*(*x*)) from model *f_θ_* and image *x*.

**Figure 20:**
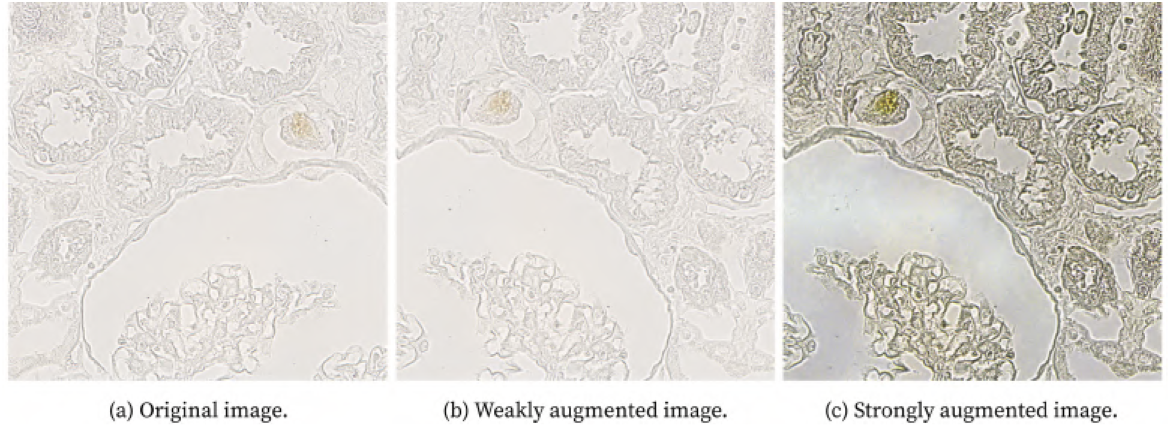
Augmentations applied to a brightfield microscopy renal image. These augmentations were adopted in the implementation of Fix-MatchSeg. The original image is image (a), (b) shows the weakly augmented image that was geometrically perturbed, and (c) shows the strongly augmented image, where colour, contrast, and blurring perturbations were applied to the output of the weak augmentation.

In FixMatchSeg, to obtain pseudo-labels, the model must predict a class distribution map for an unlabelled image. In a C-class problem, the model prediction for each pixel contains C softmax-normalised output probabilities. To obtain the pseudo-labels (*y*^), *argmax* is computed pixel-wise. However, to decide if the pseudo-label is accepted for further use, the average confidence *y* of the predicted class distribution map is computed, and if it is higher than a threshold *τ*, then the pseudo-labels are accepted. In other words, the maximum probability per pixel corresponds to the confidence of the model in predicting that label, and if the average maximum confidence per image is higher than a threshold, then those pseudo-labels will be used for loss computation and discarded otherwise.

The unsupervised loss computation is defined as follows:

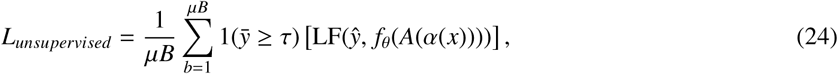

where *µ* is the ratio of unlabelled to labelled data, *B* is the labelled dataset size, LF is an arbitrary loss function, and *f_θ_*(*A*(*α*(*x*))) is the predicted class distribution map for the strongly augmented version of image *x*.

The total loss in FixMatchSeg is:

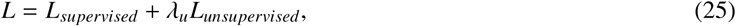

where *λ_u_*is a non-negative scalar.

### 7.2. Weak and strong data augmentations

To address consistency regularisation, weak and strong augmentations are applied to the unlabelled data. These transformations generate two similar versions of the same data sample. The weak augmentation focuses on geometric variations, while the strong augmentation alters brightness, contrast, hue saturation, and blurring. The strong augmentation is only applied to weakly augmented data, so that the objects in these two versions are precisely aligned. This is crucial because the unsupervised loss is computed between the accepted pseudo-labels of the weakly augmented sample and the model prediction of the strongly augmented version.

The chosen transformations for the weak augmentation were:

- Square symmetry with a probability of 0.5;
- Rotation between −20 and 20 degrees, with linear interpolation to “fill” the image, and a probability of 0.3;
- Elastic transform, which applies random elastic distortions to data with a probability of 0.3. This transformation is very useful because it maintains the relative positions of the features while introducing realistic deformations.

These geometric transformations were applied to an unlabelled data instance. Then, the following strong augmentations were applied to the weakly augmented image:

- CLAHE with a probability of 0.8;
- Global brightness and contrast levels were randomly altered with a probability of 0.3;
- Hue saturation was altered with a probability of 0.3;
- Gaussian blurring with probability of 0.5.

These transformations are slightly more intense than the ones adopted for the experiments of the previous section. However, to avoid excessive transformations and potentially produce unrealistic data, all the transformation-defining parameters were selected carefully and tested to make sure they corresponded with expectations.

### 7.3. Implementation details

Similarly to the previous section, these experiments were implemented in a *Python* environment, and were conducted on a machine with a NVIDIA GeForce RTX 3060 GPU with 12 GB of VRAM.

In the FixMatchSeg research, an EfficientNet-b4 model [69] was used as the U-Net’s backbone. Therefore, Fix-MatchSeg was implemented with a U-Net architecture. However, there were three different setups:

- ResNet-50 backbone with initialised pre-trained ImageNet weights;
- ResNet-50 backbone with initialised pre-trained weights from the experiments in the last section, which we will denote as RenalDataset weights;
- EfficientNet-b4 with initialised pre-trained ImageNet weights.

Each of these setups was trained with losses *L*_Dice-CE_, *L*_Dice-Focal_, and *L*_Dice-Boundary_. Again, each loss component was initialised to address the *ignore label*, to incorporate the weighted map for border pixels into computations, except for Boundary Loss, and to perform weighted mean reduction with predefined class weights.

The ratio of unlabelled to labelled data was eight, meaning that there was eight times more unlabelled data than labelled in the training dataset. The adopted batch size was nine, so that in each iteration, one labelled image and eight unlabelled images were processed. The total unsupervised loss for each batch of images was computed as in Equation 24, where LF is one of the previously mentioned hybrid loss functions, *µ* = 8, and *B* = 1, and therefore, it corresponds to the average of the unsupervised losses.

The total semi-supervised loss was computed as in Equation 25, where *λ_u_*= 0.5. After initially experimenting with a *λ_u_*= 1.0, we found that it caused unstable learning, particularly in the beginning, when pseudo-labels are not that reliable. Therefore, the *λ_u_* = 0.5 was adopted to ensure that the supervised loss brings a dominant contribution to the total loss, while the unsupervised loss still contributes, but less significantly. In addition, each experiment was repeated for the modified pseudo-label acceptance algorithm, resulting in a total of eighteen different training setups. With a transfer learning strategy for ResNet-50 and EfficientNet-b4 with ImageNet weights, the Adam optimiser [60] was initialised with a learning rate of 1 × 10^−3^ for the decoder training, and 1 × 10^−4^ for the encoder fine-tuning. For the fine-tuning of the RenalDataset weights, the Adam optimiser was initialised with a learning rate of 1 × 10^−4^. To adjust learning rate and escape potential loss plateaus, *ReduceLROnPlateau* method was adopted with a minimum learning rate of 1 × 10^−6^, a patience of 8 epochs, and a reduction factor of 10.

The data augmentation mechanisms were incorporated into all training pipelines using the *Albumentations* library.

### 7.4. Results

The following tables and images aggregate the experimental results of this section on semi-supervised learning for semantic segmentation. The performance metrics’ results were averaged across all the tested samples, and the same image set was utilised to test all model-loss configurations.

Table 16 and Table 17 compare performance results for the FixMatchSeg method and different custom loss functions. Here, a U-Net architecture was used with an EfficientNet-b4 backbone. The results obtained with the loss function *L*_Dice-CE_ distinguish themselves by improved performance. This strategy achieved evaluation results up to par with the best configurations of the tested supervised methods for the *tubules*, *interstitium*, and *empty space* classes. However, it did not outperform for the remainder of the structure classes. Figure 21 illustrates an example of an original tile, the annotated ground-truth, and the inference of the FixMatchSeg method implemented with U-Net (EfficientNet-b4 backbone, initialised with ImageNet weights) (*L*_Dice-CE_).

**Figure 21:**
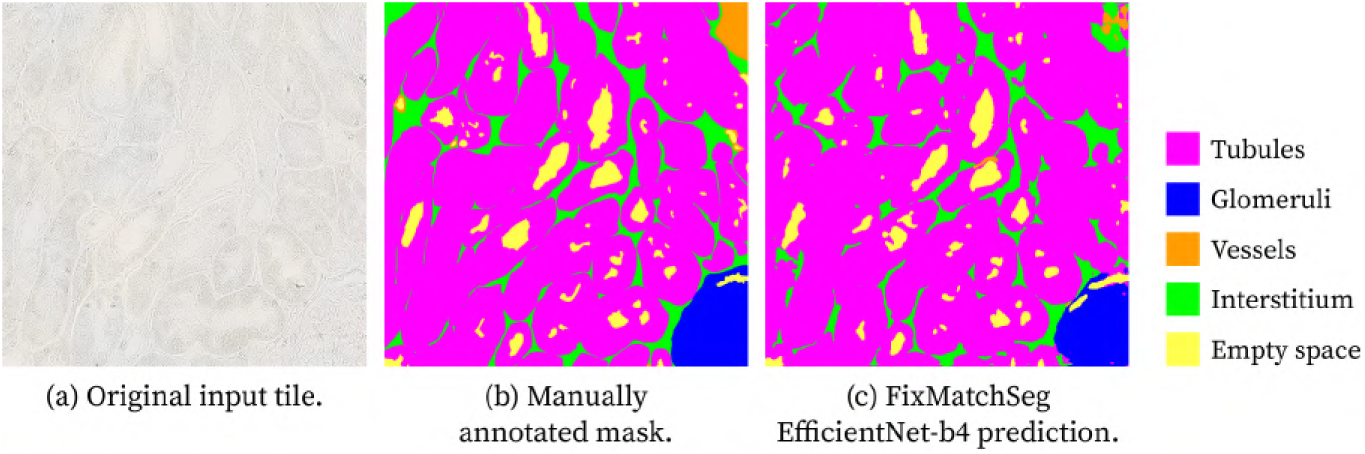
Example of predicted segmentation masks obtained with **FixMatchSeg** (U-Net with **E**ffi**cientNet-b4 backbone, initialised with ImageNet weights**) and *L*_Dice-CE_ loss function. Panels: (a) Original brightfield image of unstained kidney tissue as input tile, (b) manually annotated mask, and (c) FixMatchSeg prediction. *Colour code: tubules (magenta), glomeruli (blue), vessels (orange), interstitium (green), and empty space (yellow)*.

**Table 16:** Average Partition Similarity for multi-class segmentation with FixMatchSeg (U-Net with EfficientNet-b4 backbone, initialised with ImageNet weights).

| **Loss Function** | **Average Partition Similarity** (%) |
|---|---|
| *L*<sub>*Dice-CE*</sub> | 84.3 |
| *L*<sub>*Dice-Focal*</sub> | 83.0 |
| *L*<sub>*Dice-Boundary*</sub> | 81.6 |

**Table 17:**
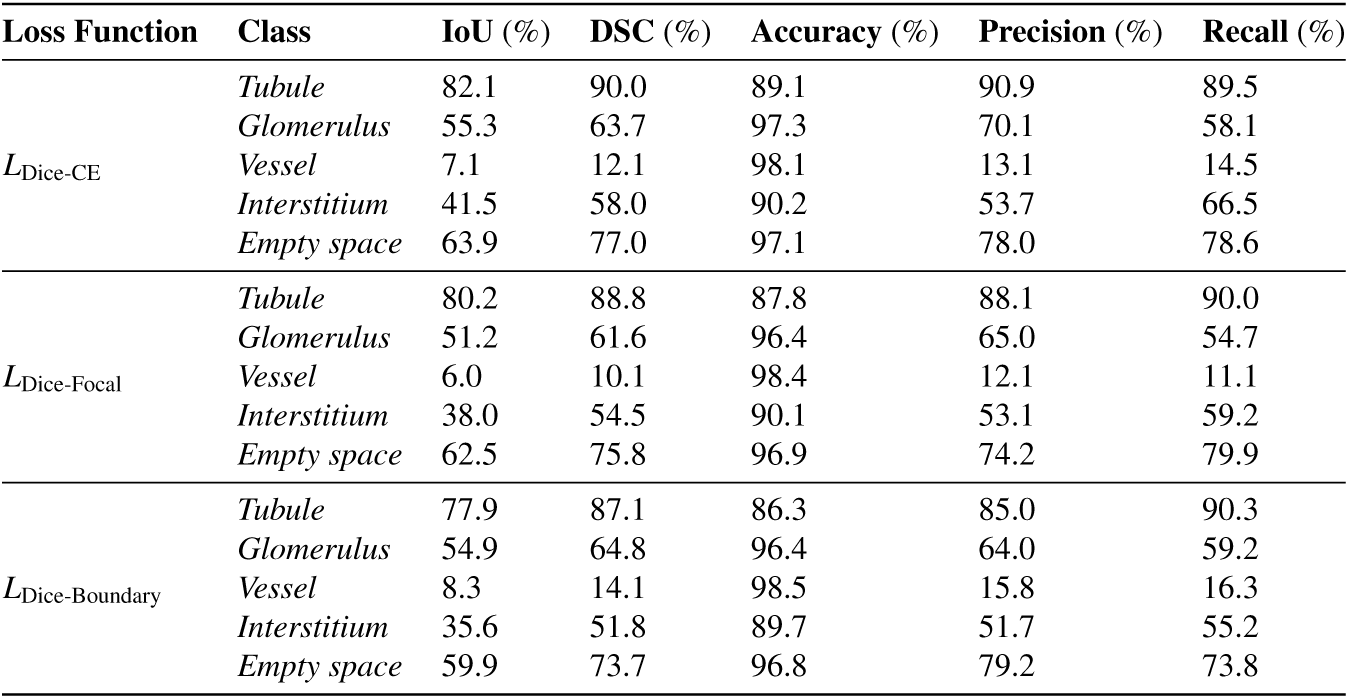
Average performance metrics results for multi-class segmentation with FixMatchSeg (U-Net with EfficientNet-b4 backbone, initialised with ImageNet weights) with different loss functions.

Table 18 and Table 19 compare performance results for the semi-supervised strategy that uses a U-Net with a ResNet-50 backbone initialised with pre-trained ImageNet weights. For the first time, the hybrid loss function *L*_Dice-Boundary_ produced the best segmentation results. It was the loss configuration that worked best at predicting *glomerulus*. However, despite this semi-supervised method using eight times more images (with the unlabelled samples) for training than its supervised version (Table 11), it does not achieve better segmentation performance. Figure 22 illustrates an example of an original tile, the annotated ground-truth, and the inference of the FixMatchSeg method implemented with U-Net (with ResNet-50 backbone, initialised with ImageNet weights) (*L*_Dice-Boundary_).

**Figure 22:**
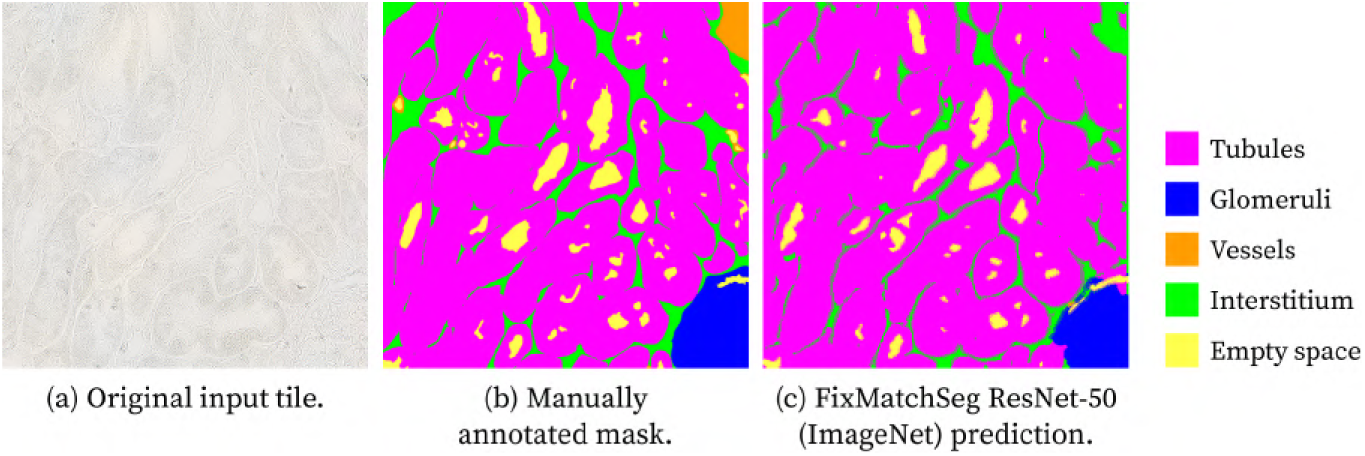
Example of predicted segmentation masks obtained with **FixMatchSeg** (U-Net with **ResNet50 backbone, initialised with ImageNet weights**) and *L*_Dice-Boundary_ loss function. Panels: (a) Original brightfield image of unstained kidney tissue as input tile, (b) manually annotated mask, and (c) FixMatchSeg prediction. *Colour code: tubules (magenta), glomeruli (blue), vessels (orange), interstitium (green), and empty space (yellow)*.

**Table 18:**
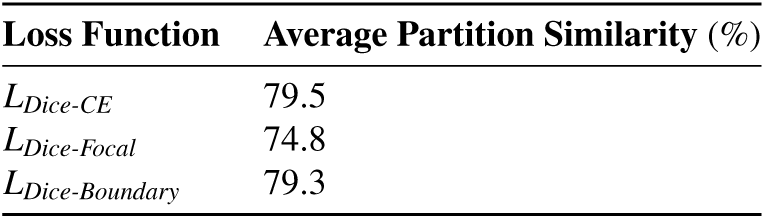
Average Partition Similarity for multi-class segmentation with FixMatchSeg (U-Net with ResNet-50 backbone, initialised with ImageNet weights).

**Table 19:** Average performance metrics results for multi-class segmentation with FixMatchSeg (U-Net with ResNet-50 backbone, initialised with ImageNet weights) with different loss functions.

| **Loss Function** | **Class** | **IoU** (%) | **DSC** (%) | **Accuracy** (%) | **Precision** (%) | **Recall** (%) |
|---|---|---|---|---|---|---|
| *L*<sub>Dice-CE</sub> | *Tubule* | 76.7 | 86.5 | 85.8 | 90.0 | 84.0 |
|  | *Glomerulus* | 39.8 | 48.9 | 95.2 | 71.4 | 43.0 |
|  | *Vessel* | 4.1 | 7.3 | 97.9 | 8.2 | 9.9 |
|  | *Interstitium* | 36.6 | 52.7 | 86.4 | 42.8 | 72.7 |
|  | *Empty space* | 56.9 | 71.1 | 96.8 | 84.8 | 65.1 |
| *L*<sub>Dice-Focal</sub> | *Tubule* | 65.9 | 77.1 | 86.8 | 83.9 | 73.5 |
|  | *Glomerulus* | 42.6 | 52.9 | 95.4 | 42.7 | 56.2 |
|  | *Vessel* | 4.2 | 7.1 | 98.9 | 16.7 | 5.7 |
|  | *Interstitium* | 32.4 | 47.4 | 90.8 | 43.5 | 58.7 |
|  | *Empty space* | 53.4 | 67.6 | 95.4 | 58.1 | 80.8 |
| *L*<sub>Dice-Boundary</sub> | *Tubule* | 74.4 | 84.4 | 85.0 | 90.1 | 81.6 |
|  | *Glomerulus* | 47.2 | 56.3 | 93.3 | 48.7 | 62.3 |
|  | *Vessel* | 5.5 | 9.8 | 98.7 | 19.1 | 9.7 |
|  | *Interstitium* | 33.5 | 49.5 | 89.0 | 48.1 | 53.8 |
|  | *Empty space* | 60.4 | 74.0 | 96.8 | 78.9 | 74.6 |

Table 20 and Table 21 compare performance results for the FixMatchSeg approach that uses a U-Net with a ResNet-50 backbone initialised with pre-trained weights from the research in the previous section. The best-performing loss configuration was *L*_Dice-Focal_, which is curious because it was not the loss that achieved the highest performance for the supervised version of the same model. Figure 23 illustrates an example of a segmentation mask whose labels were predicted by this FixMatchSeg approach. This was the best training setup between the FixMatch-Seg variants, which was expected since the model was pre-trained on this specific medical domain, in contrast with the natural domain of the ImageNet dataset. However, this model did not outperform the U-Net trained under a supervised setting. This indicates that the generated pseudo-labels didn’t have enough quality to further improve the model’s segmentation.

**Figure 23:**
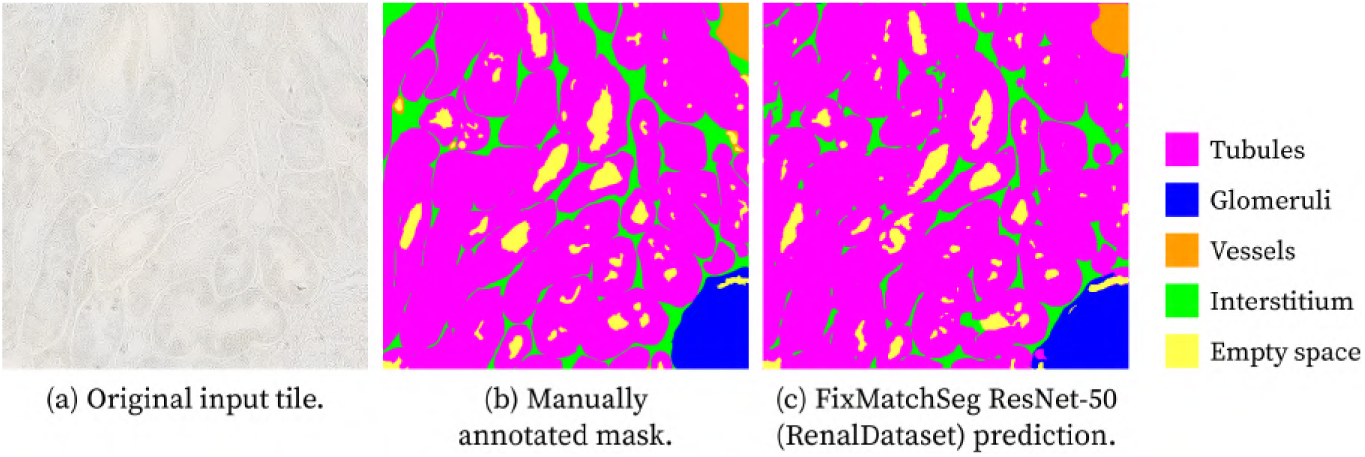
Example of predicted segmentation masks obtained with **FixMatchSeg** (U-Net with **ResNet50 backbone, initialised with Renal-Dataset weights**) and *L*_Dice-Focal_ loss function. Panels: (a) Original brightfield image of unstained kidney tissue as input tile, (b) manually annotated mask, and (c) FixMatchSeg prediction. *Colour code: tubules (magenta), glomeruli (blue), vessels (orange), interstitium (green), and empty space (yellow)*.

**Table 20:**
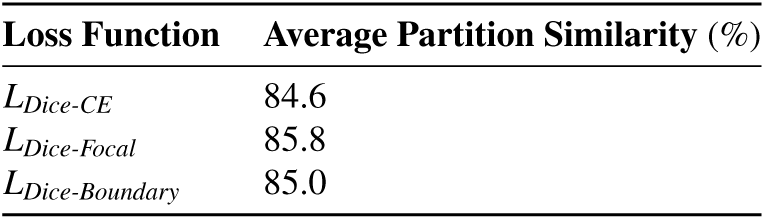
Average Partition Similarity for multi-class segmentation with FixMatchSeg (U-Net with ResNet-50 backbone, initialised with Renal-Dataset weights).

| **Loss Function** | **Average Partition Similarity** (%) |
|---|---|
| *L<sub>Dice-CE</sub>* | 84.6 |
| *L<sub>Dice-Focal</sub>* | 85.8 |
| *L<sub>Dice-Boundary</sub>* | 85.0 |

**Table 21:**
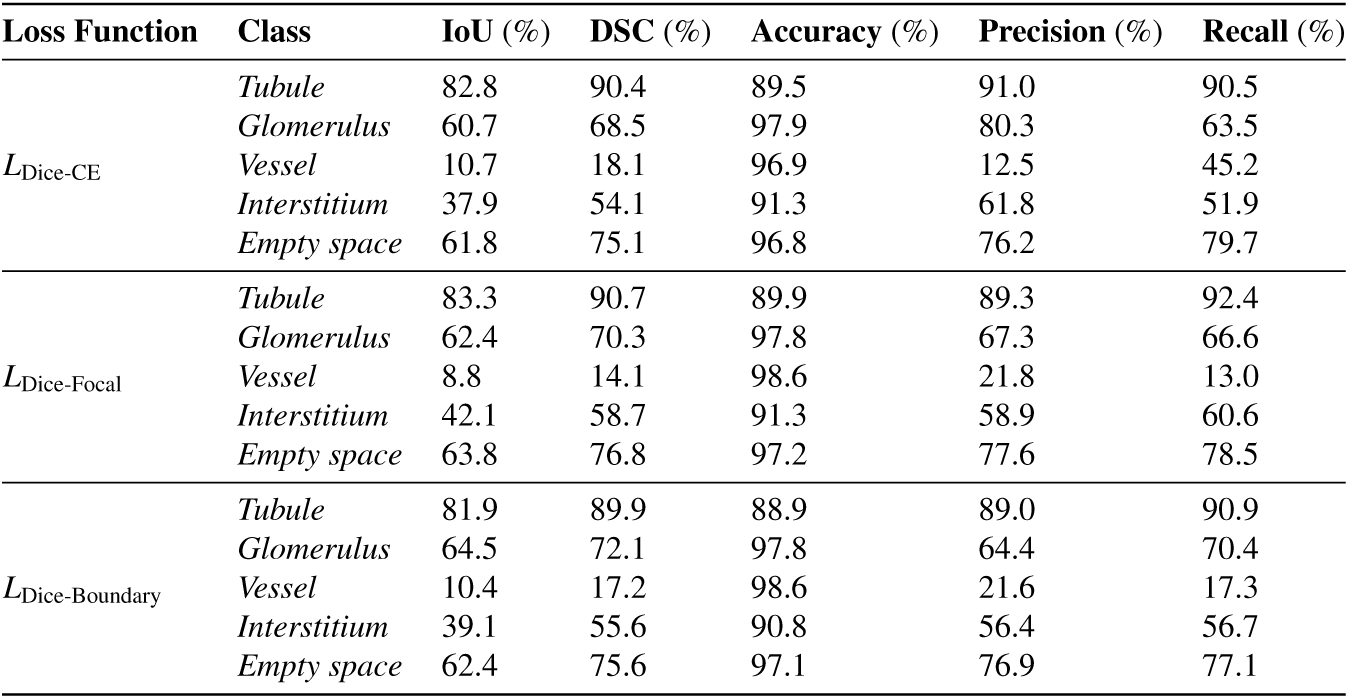
Average performance metrics results for multi-class segmentation with FixMatchSeg (U-Net with ResNet-50 backbone, initialised with RenalDataset weights) using different loss functions.

### 7.5. Discussion

The three best-performing FixMatchSeg model-loss combinations were:

- U-Net with EfficientNet-b4 backbone initialised with ImageNet weights using the *L*_Dice-CE_ loss function;
- U-Net with ResNet50 backbone initialised with ImageNet weights using the *L*_Dice-Boundary_ loss function;
- U-Net with ResNet50 backbone initialised with RenalDataset weights using the *L*_Dice-Focal_ loss function.

Among the FixMatchSeg variants, the model initialised with pre-trained weights from the RenalDataset achieved the most accurate segmentation. Figure 24 illustrates three examples with the best-performing configurations. Additional results are provided in Figures 31 in the Appendix.

**Figure 24:**
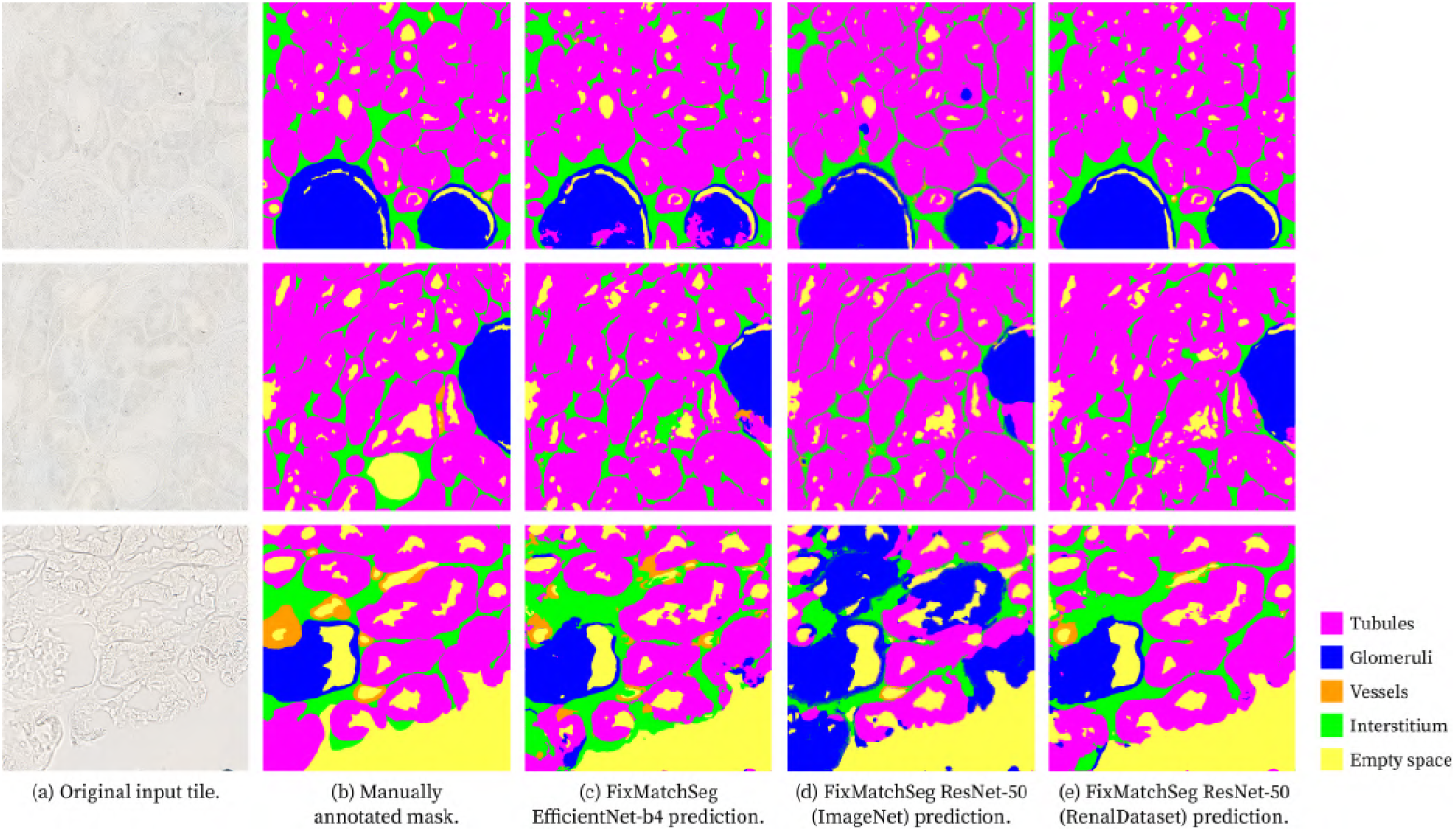
Multi-class segmentation masks output using the FixMatchSeg methods and respective manual annotations, shown for three different example images (top, middle, and bottom rows). For each image: (a) Original image, (b) Manually annotated masks, (c) FixMatchSeg with EfficientNet-b4 backbone, (d) FixMatchSeg with ResNet-50 backbone initialised with ImageNet weights, and (e) FixMatchSeg with ResNet-50 backbone initialised with RenalDataset weights. *Colour code: tubules (magenta), glomeruli (blue), vessels (orange), interstitium (green), and empty space (yellow)*.

The segmentation masks in Figure 24, particularly those corresponding to EfficientNet-b4 and to ResNet-50 (ImageNet) models, are visibly different from the ground-truth, with some artefacts and misclassified pixels. In contrast, the predictions generated by U-Net with a ResNet-50 backbone initialised with RenalDataset weights are closer to the true annotations, and visually similar to the supervised segmentation outputs shown in Figure 16 from the advanced models’ implementation section.

Overall, the supervised models in the previous section – U-Net, DeepLabv3+, and TransUNet – outperformed the semi-supervised FixMatchSeg methods. However, since those models were trained on fewer data instances, even with the use of data augmentation, they may be less generalizable. On the contrary, the FixMatchSeg models were trained on eight times more data, and were additionally submitted to weak and strong augmentations to improve generalisation. The more robust training setup could explain the slightly lower performance of these models on the evaluation set, although they may be better prepared to segment previously unseen data.

## 8. Post-processing of the segmentation masks

Post-processing techniques can be very beneficial to remove small artefacts in the segmentation masks, to improve boundaries, and fill small holes inside larger objects. However, excessive post-processing is extremely hard to generalise, and therefore, should be avoided and kept to a minimum.

Since the segmentation performance for the *vessel* class was poor, no post-processing techniques were applied to it. In most cases, few pixels were assigned the *vessel* label, and often, only small portions of whole vessel instances were identified. Therefore, no further processing was applied to ensure that these already small regions were not affected or removed.

The class of glomeruli instances was the main target of this post-processing stage, due to their distinct shape compared to the other classes. The instances from this class are characterised by their round shape and are usually larger than other objects. Nevertheless, segmenting glomeruli is not straightforward, as their appearance can vary due to different slicing angles upon the biopsy extraction. The glomerulus can exhibit a less or more uniformly round shape depending on the cut. In reality, this aspect can affect all structures and not only glomeruli; however, their singular shape makes it more noticeable to the naked eye. Figure 25 demonstrates tiles extracted from different biopsies and the irregularity in the glomeruli’s shape.

**Figure 25:**
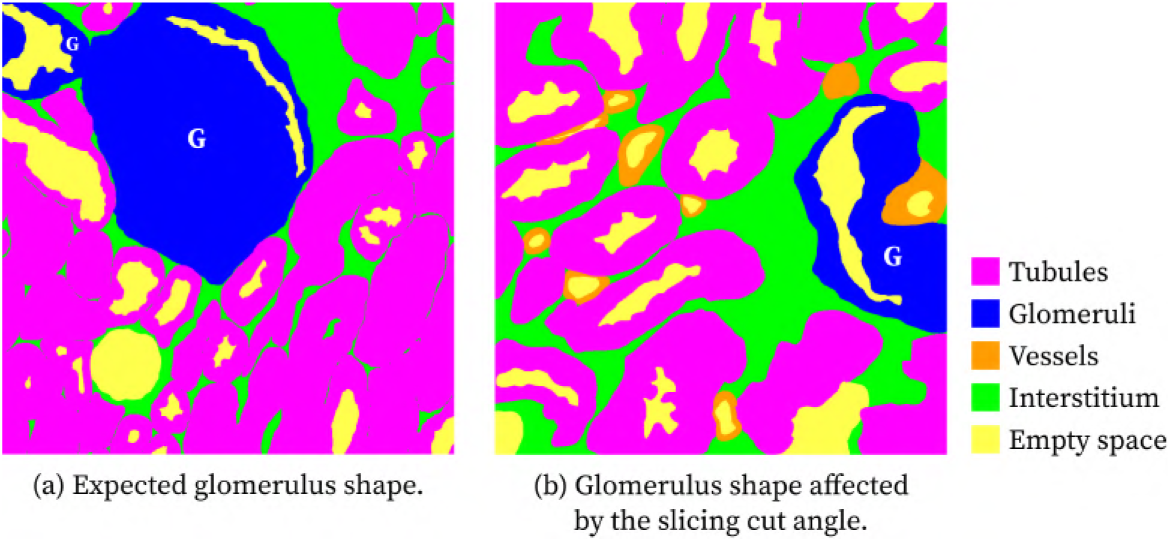
Example of annotation masks from different biopsy images presenting glomeruli (in blue, annotated by G) with different silhouettes. *Colour code: tubules (magenta), glomeruli (blue), vessels (orange), interstitium (green), and empty space (yellow)*.

The main rules that defined the post-processing techniques were derived from domain-specific knowledge:

1. In this segmentation setup, only empty space should be distinguished inside glomeruli, and all other structure classes should not be found inside the borders of a glomerulus.
2. Typically, glomeruli are larger than other structures, so small predicted objects that were classified as *glomerulus* should be removed.
3. Small *tubules* artefacts should be removed.
4. There is no occurrence of small *empty space* objects in the interstitium tissue, so appearances of small *empty space* pixel groupings bounded by interstitium should be removed.
5. Large empty areas, characterised by the absence of tissue, should not have sporadic instances of tissue.

### 8.1. Post-processing of the glomeruli segmentation class predictions

Based on the first rule, no other renal structure, except *empty space*, should be identified inside a glomerulus. Consequently, the first post-processing step was to fill small “holes” inside these structures. These holes are pixel groupings of class labels different from *glomerulus*. To address these internal holes, a binary mask *glomerulus* was created, so that *glomeruli* pixels were set to one, and the rest were set to zero. Then, an elliptical kernel of size 19×19 was chosen due to the round morphology of glomeruli. This kernel was used to close the holes in the binary mask with the *morphologyEx* method configured to perform the morphological closing operation from the *OpenCV* library. After this step, the newly assigned *glomeruli* pixels were updated in the original segmentation mask, and the previously assigned *empty space* pixels were restored.

To address the second rule, bounding boxes were computed for a new *glomerulus* binary mask, and the ones whose area was below a predefined threshold or that had low pixel density were flagged for removal. After experimenting with different threshold values, the chosen threshold was 6000 pixels. Additionally, bounding boxes were considered sparse if the number of *glomeruli* pixels inside them was lower than 35% of the bounding box area. These bounding boxes were extracted from the binary mask by first applying the Canny Edge algorithm [70] to identify object contours. The edges were dilated for refinement, and the *findContours* method from the *OpenCV* library was used to retrieve the outermost contours. Finally, the *boundingRect* function from the same library returned the list of bounding boxes. The pixels within each bounding box that were flagged for removal were reclassified using the most common class label from the frontier pixels outside the bounding box.

### 8.2. Post-processing of the tubules, interstitium, and empty-space segmentation classes predictions

For the *tubules*, *interstitium*, and *empty-space* segmentation classes, no morphological closing operation was performed. Achieving a generalised closing operation was nearly impossible, since instances from these classes, in particular the tubules, are adjacent or overlapping, and the closing operation often led to instances being merged together when they should be distinct objects. Nevertheless, other post-processing techniques were applied.

According to the third rule, *tubules* artefacts should be removed. To eliminate these artefacts, a binary *tubule* mask was created, so that *tubules* pixels were set to one, and the rest were set to zero. Then, *tubules* bounding boxes were extracted from the binary mask, and the ones with areas smaller than 2000 pixels were considered artefacts and were removed by replacing their pixels with the most common class label outside the border of each bounding box.

For the last two rules, bounding boxes were extracted from *interstitium* and *empty space* masks separately. Bounding boxes were flagged for removal only if they were bounded by a specific class and their area was smaller than a threshold. Specifically, *interstitium* boxes of less than 1000 pixels that were surrounded by empty space were removed, while *empty space* bounding boxes with areas smaller than 600 pixels surrounded by interstitium were also removed. In these scenarios, the pixels inside the flagged bounding boxes were directly replaced by the surrounding tissue class.

Further processing of the *intertitium* or *empty space* classes was avoided. For instance, the tubules can be close together, which creates very small interstitial portions. However, since the interstitium is the tissue that resides between other tissue structures, there are vast interstitial areas in other regions. Analogously, tiny empty space regions can be found inside glomeruli or tubules, while other tiles can contain a lot of empty space due to tissue absence. These domain-specific characteristics make it difficult to create post-processing rules that generalise well across different tissue slides.

### 8.3. Results and Discussion

The performance metrics results are displayed in the tables below. These results correspond to the predictions of the best model and loss function configurations after post-processing.

Table 22 demonstrates that the U-Net *L*_Dice-CE_ configuration has the highest Partition Similarity evaluation score.

**Table 22:** Average Partition Similarity after post-processing for multi-class segmentation with the best model and training loss configurations.

| **Configuration** | **Average Partition Similarity** (%) |
|---|---|
| *U-Net L<sub>Dice-CE</sub>* | 87.0 |
| *DeepLabv3+ L<sub>Dice-Focal</sub>* | 86.3 |
| *TransUNet L<sub>Dice-CE</sub>* | 83.3 |
| *FixMatchSeg U-Net EfficientNet (ImageNet) L<sub>Dice-CE</sub>* | 84.7 |
| *FixMatchSeg U-Net ResNet-50 (ImageNet) L<sub>Dice-Boundary</sub>* | 79.8 |
| *FixMatchSeg U-Net ResNet-50 (RenalDataset) L<sub>Dice-Focal</sub>* | 86.1 |

Tables 23, 27, and 28 reveal the segmentation evaluation results of the processed prediction masks of the supervised U-Net, DeepLabv3+, and TransUNet. Only the U-Net results were improved by the post-processing. DeepLabv3+ and TransUNet didn’t particularly benefit from the additional processing. One could argue that the segmentation masks inferred by these models had minimal noise, and consequently, there was no distinct enhancement of the masks. Figure 26 illustrates an example of an original segmentation mask and the refined version for the U-Net model. In this particular case, the *tubules* artefacts were noticeably eliminated.

**Figure 26:**
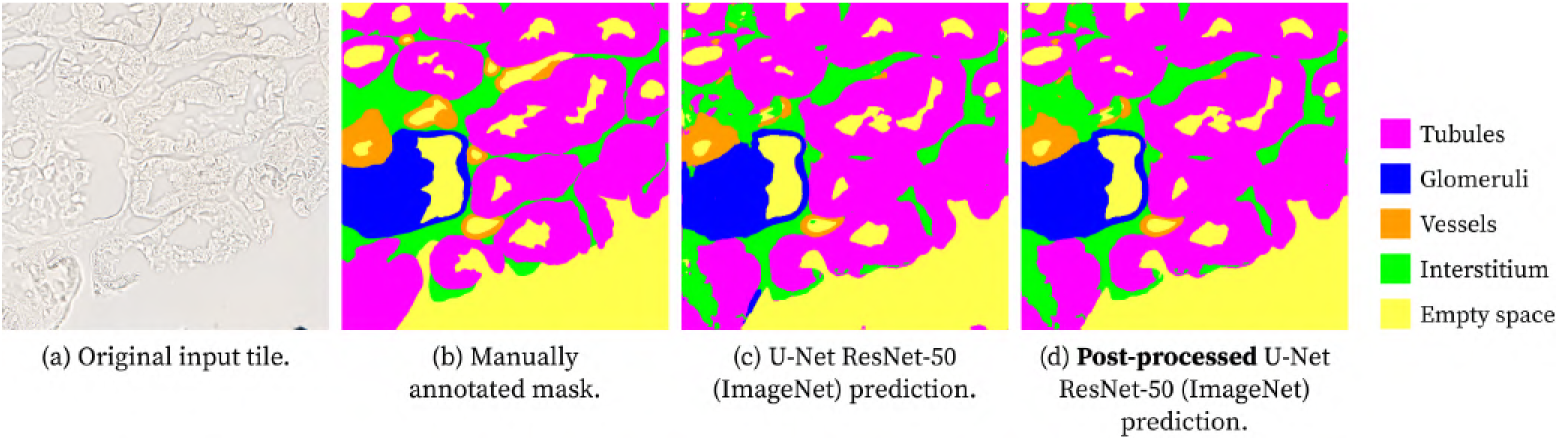
Example of predicted segmentation masks before and after post-processing, obtained using **U-Net** with a ResNet-50 backbone and *L*_Dice-CE_ loss function. Panels: (a) Original brightfield image of unstained kidney tissue as input tile, (b) manually annotated mask, (c) U-Net original prediction, and (d) U-Net’s segmentation mask after post-processing. *Colour code: tubules (magenta), glomeruli (blue), vessels (orange), interstitium (green), and empty space (yellow)*.

**Table 23:** Average performance metrics results after post-processing for multi-class segmentation with U-Net (ResNet-50 backbone) and *L*_Dice-CE_ loss function.

| **Class** | **IoU** (%) | **DSC** (%) | **Accuracy** (%) | **Precision** (%) | **Recall** (%) |
|---|---|---|---|---|---|
| *Tubule* | 84.7 | 91.6 | 90.5 | 89.0 | 94.6 |
| *Glomerulus* | 65.9 | 71.6 | 98.4 | 92.6 | 69.4 |
| *Vessel* | 11.0 | 17.6 | 98.5 | 21.8 | 21.4 |
| *Interstitium* | 42.7 | 59.1 | 92.0 | 62.9 | 57.6 |
| *Empty space* | 62.1 | 75.4 | 97.2 | 83.1 | 70.9 |

In contrast to the other models, the FixMatchSeg methods greatly benefited from the post-processing and their results are showcased in Tables 24, 25, and 26. This could be expected, since their predictions were noisier and had more artefacts, which were successfully processed and removed with the added techniques. The corresponding evaluation results improved, and the visual impact is significant, as can be seen in Figures 27 and 28.

**Figure 27:**
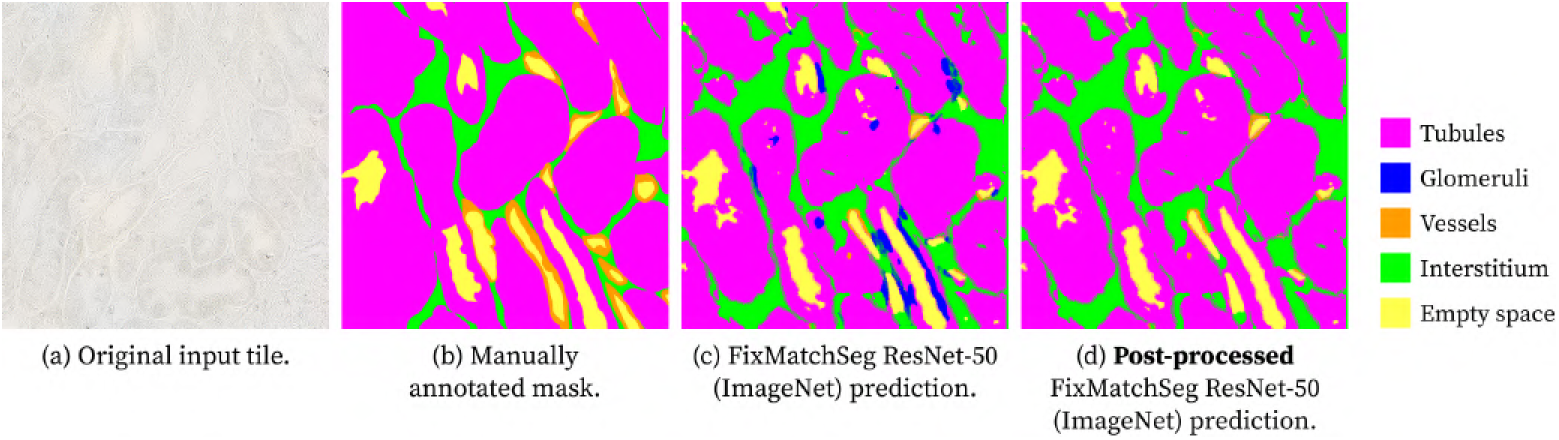
Example of predicted segmentation masks before and after post-processing, obtained using **FixMatchSeg** (U-Net with **ResNet50 backbone, initialised with ImageNet weights**) and *L*_Dice-Boundary_ loss function. Panels: (a) Original brightfield image of unstained kidney tissue as input tile, (b) manually annotated mask, (c) FixMatchSeg original prediction, and (d) FixMatchSeg’s segmentation mask after post-processing. *Colour code: tubules (magenta), glomeruli (blue), vessels (orange), interstitium (green), and empty space (yellow)*.

**Figure 28:**
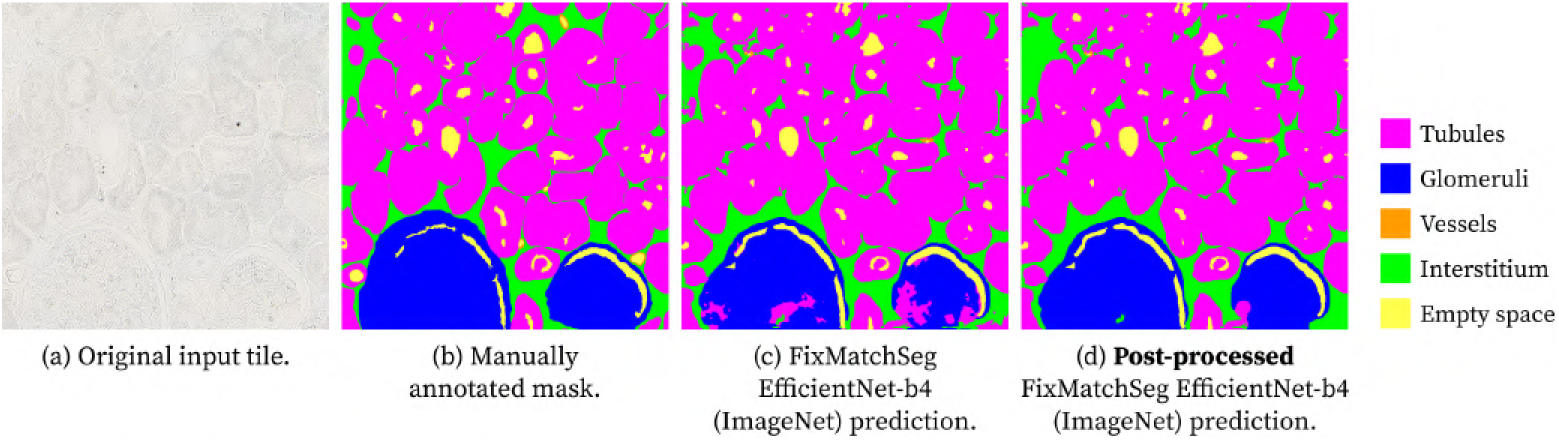
Example of predicted segmentation masks before and after post-processing, obtained using **FixMatchSeg** (U-Net with **E**ffi**cientNet-b4 backbone, initialised with ImageNet weights**) and *L*_Dice-Boundary_ loss function. Panels: (a) Original brightfield image of unstained kidney tissue as input tile, (b) manually annotated mask, (c) FixMatchSeg original prediction, and (d) FixMatchSeg’s segmentation mask after post-processing. *Colour code: tubules (magenta), glomeruli (blue), vessels (orange), interstitium (green), and empty space (yellow)*.

**Table 24:** Average performance metrics results after post-processing for multi-class segmentation with FixMatchSeg (U-Net with EfficientNet-b4 backbone, initialised with ImageNet weights) and *L*_Dice-CE_ loss function.

| **Class** | **IoU** (%) | **DSC** (%) | **Accuracy** (%) | **Precision** (%) | **Recall** (%) |
|---|---|---|---|---|---|
| *Tubule* | 82.6 | 90.3 | 89.4 | 91.8 | 89.4 |
| *Glomerulus* | 57.0 | 63.4 | 97.8 | 88.3 | 59.0 |
| *Vessel* | 7.1 | 12.1 | 98.1 | 13.1 | 14.6 |
| *Interstitium* | 41.5 | 58.1 | 90.2 | 53.5 | 66.9 |
| *Empty space* | 63.6 | 76.8 | 97.0 | 77.5 | 78.8 |

**Table 25:** Average performance metrics results after post-processing for multi-class segmentation with FixMatchSeg (U-Net with ResNet-50 backbone, initialised with ImageNet weights) and *L*_Dice-Boundary_ loss function.

| **Class** | **IoU** (%) | **DSC** (%) | **Accuracy** (%) | **Precision** (%) | **Recall** (%) |
|---|---|---|---|---|---|
| *Tubule* | 74.7 | 84.6 | 85.2 | 90.1 | 82.0 |
| *Glomerulus* | 48.7 | 56.3 | 93.9 | 69.3 | 62.4 |
| *Vessel* | 5.4 | 9.6 | 98.7 | 18.9 | 9.5 |
| *Interstitium* | 33.2 | 49.0 | 89.1 | 48.7 | 52.4 |
| *Empty space* | 59.6 | 73.5 | 96.6 | 77.6 | 75.1 |

**Table 26:** Average performance metrics results after post-processing for multi-class segmentation with FixMatchSeg (U-Net with ResNet-50 backbone, initialised with RenalDataset weights) and *L*_Dice-Focal_ loss function.

| **Class** | **IoU** (%) | **DSC** (%) | **Accuracy** (%) | **Precision** (%) | **Recall** (%) |
|---|---|---|---|---|---|
| *Tubule* | 83.7 | 90.9 | 90.1 | 89.8 | 92.3 |
| *Glomerulus* | 63.2 | 69.8 | 98.0 | 88.2 | 67.2 |
| *Vessel* | 8.9 | 14.2 | 98.6 | 21.8 | 13.1 |
| *Interstitium* | 42.4 | 58.9 | 91.3 | 59.0 | 61.0 |
| *Empty space* | 63.3 | 76.5 | 97.1 | 77.0 | 78.6 |

## 9. Discussion

Renal biopsies are a crucial tool for gaining knowledge of the state of the nephron. Nowadays, a pathologist manually analyses the tissue to understand the amount of damage to the kidney and, in necessary cases, to diagnose renal disease. The largest-sized needle used to perform biopsies produces a sample with a diameter of approximately 2 mm [3]. Therefore, pathologists must meticulously study digitised images of the microscopic samples, looking for histomorphological alterations in the renal structures. That said, automatically segmenting these structures would become an essential tool for pathologists to perform a more efficient and faster examination.

In the current standard workflow of biopsy collection, the samples go through a staining process that is costly and lengthy. Moreover, artefacts can affect the quality of the sample, and, on occasion, these can be introduced during the staining process. To our knowledge, no publications have studied the segmentation of unstained kidney biopsy images. We are interested in analysing the feasibility of skipping the staining step from the biopsy collection process and achieving favourable results in automatic segmentation. This would create a cheaper and faster workflow for the analysis of renal tissue.

The unstained renal tiles dataset that the team has gathered so far could be complemented by other renal datasets. However, due to the lack of public datasets of this kind, it would be relevant in future work to use a dataset with regular stained images and artificially remove the staining or, in the opposite direction, to use colourisation models to stain our images. This would produce synthetic input images for the segmentation network.

Foundation segmentation models such as SAM and SAM2 have shown good zero-shot segmentation performance in natural images. However, our experiments reveal that these models are not prepared for zero-shot performance in medical tasks without further fine-tuning. While MedSAM presents a promising development as a fine-tuned version of SAM for medical tasks, its direct use does not fit our requirements for a fully automated approach.

The outputs from our U-Net, DeepLabv3+, and TransUNet experiments demonstrate the potential of unstained images, despite their original poor contrast. FixMatchSeg, a semi-supervised learning approach, was proposed with the goal of increasing the training data size by including unlabelled data. While this approach aimed to achieve better generalisation to unseen data, it did not produce better results than the previous supervised methods, which can be attributed to the use of pseudo-labels. Consequently, we should explore other semi-supervised approaches with stricter methods than pseudo-labelling algorithms, capable of avoiding confirmation bias, such as Generative Adversarial Networks.

Post-processing techniques did, in fact, improve the evaluation metrics of the FixMatchSeg variants. However, they did not significantly refine the U-Net, DeepLabv3+, and TransUNet outputs. This shows that when the segmentation masks already closely align with the ground-truth, post-processing can even introduce new artefacts or under-segmentation. This means that a significant improvement in segmentation performance should not rely on post-processing techniques, but rather be a result of a well-trained model.

## 10. Conclusion

This work surveyed state-of-the-art segmentation methodologies in the medical imaging field, particularly focusing on renal histopathological images. We examined both semantic and instance segmentation methods, as well as the architecture of the networks used in these approaches. In recent years, researchers have been adopting instance-based methods. Ronneberger et al. definitely impacted this research field when they introduced U-Net’s famous encoder-decoder architecture [13]. It still inspires and is the base of most approaches. Some foundation models, such as SAM and SAM2, have also been demonstrating great results and impacting the progress in the segmentation field. However, since they were trained on open-world images, they are inadequate for segmenting medical images without prior fine-tuning. A different path that we could take in the future, in comparison to the task-specific segmentation architectures, would be to fine-tune these foundation models in our dataset.

The findings, according to the test data, show that both U-Net and DeepLabv3+ already provide reliable segmentation masks that can be introduced into real-world pathology applications and are suitable for use in downstream tasks. A partner medical pathologist at the Anatomical Pathology Service of ULSSJ, and a partner biomedical engineer and annotator specialist at i3S, have both approved the reliability and utility of these results.

One notable application is that it is possible to establish direct structure mapping between the produced segmentation masks and the immunofluorescence slides with fluorescent peroxidasin protein markers. After the mapping is complete, the assessment of potential histomorphological alterations can be easily identified by varying expression levels of the protein in each renal structure. This application is truly beneficial to pathologists and biomedical researchers since it reduces complexity and time spent on analysis while ensuring accuracy in locating renal structures.

The reviewed datasets used in these studies demonstrate gaps in the Computational Pathology field regarding the scarcity of public renal brightfield microscopy image datasets, particularly the non-existence of unstained collections. With the contribution of our dataset, we aim to promote advancements in this area.

Future work should aim to distinguish between distal and proximal tubules, given their distinct roles in renal physiology. Additionally, a more in-depth investigation is needed into the automatic segmentation of samples exhibiting histomorphological alterations. This task is inherently more complex than segmenting structures with normal morphology, as pathological changes can distort shape and obscure boundaries, leading to greater intra-class variability.

The subtle performance differences observed among the evaluated methods may also stem from limitations in the adopted evaluation metrics, which may fail to capture clinically relevant segmentation characteristics. In particular, metrics aligned with downstream tasks, such as biomarker quantification and disease diagnosis, are essential for assessing the practical utility of segmentation models.

Finally, exploring the complementary approach of virtual staining [71] may offer a promising avenue for segmenting unstained samples. Virtual staining techniques aim to synthetically generate stained representations from unstained tissue, potentially enabling the integration of both stained and unstained datasets to train more robust segmentation models. Although this approach presents its own challenges, successful implementation could significantly enhance stain-independent analysis in computational pathology.

## Ethics approval

This study was conducted in accordance with the Declaration of Helsinki. The research protocol was reviewed and approved by the institutional Health Ethics Commission (reference number CE-04-2023) and the Data Protection Officer of Unidade Local de Saúde de São João (ULSSJ), Porto, Portugal.

## Data Availability

The dataset generated and analysed during the current study is available in the HUKID (*HUman KIDney*) repository, http://to_be_disclosed_upon_paper_acceptance.

## Code Availability

The underlying code, training, testing, and validation datasets for this study are available in the HUKID-Seg repository and can be accessed via this link http://to_be_disclosed_upon_paper_acceptance.

## Acknowledgements

The kidney tissue samples used in this study were kindly provided by the Department of Anatomic Pathology of Unidade Local de Saúde de São João (ULSSJ), Porto, Portugal.

## Authors Contribution

M.C., I.S.A., and J.S.C. conceptualised the systematic review; J.P.O., R.S, and I.B. were responsible for primary data collection; I.B. was responsible for primary data preparation and annotation; M.C. and J.S.C. designed the experiments; M.C. conducted the experiments and the analysis of the results; M.C. drafted the manuscript; All authors revised the manuscript; J.S.C. supervised the project. All authors read and approved the final manuscript.

## 11. Appendix

**Figure 29:**
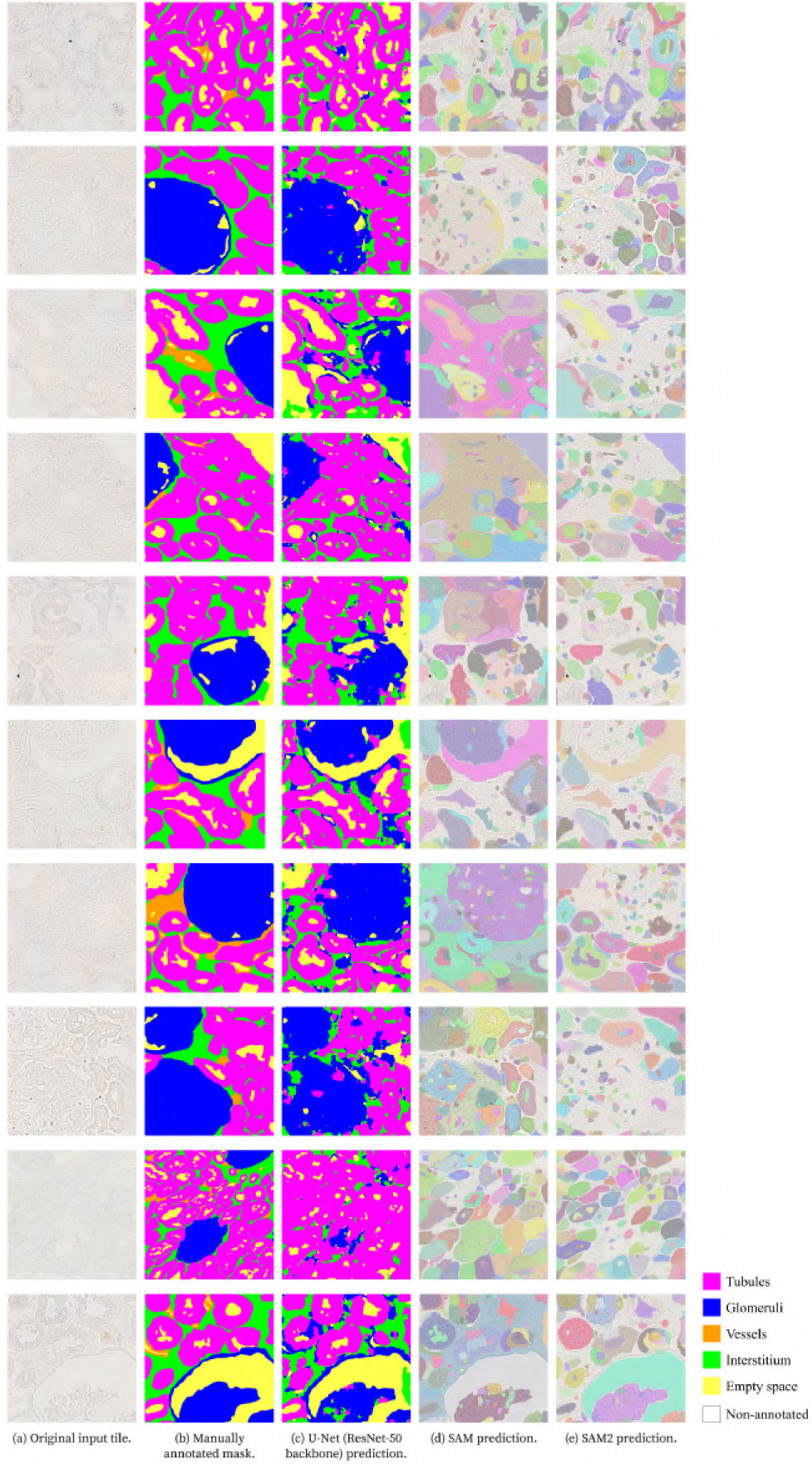
Multi-class segmentation masks output using deep learning models and respective manual annotations, shown for ten different example images (rows). For each image: (a) Original image, (b) Manually annotated masks, (c) U-Net (ResNet-50 backbone), (d) SAM, and (e) SAM2. *Colour code: tubules (magenta), glomeruli (blue), vessels (orange), interstitium (green), empty space (yellow), and non-annotated tissue (white)*.

**Figure 30:**
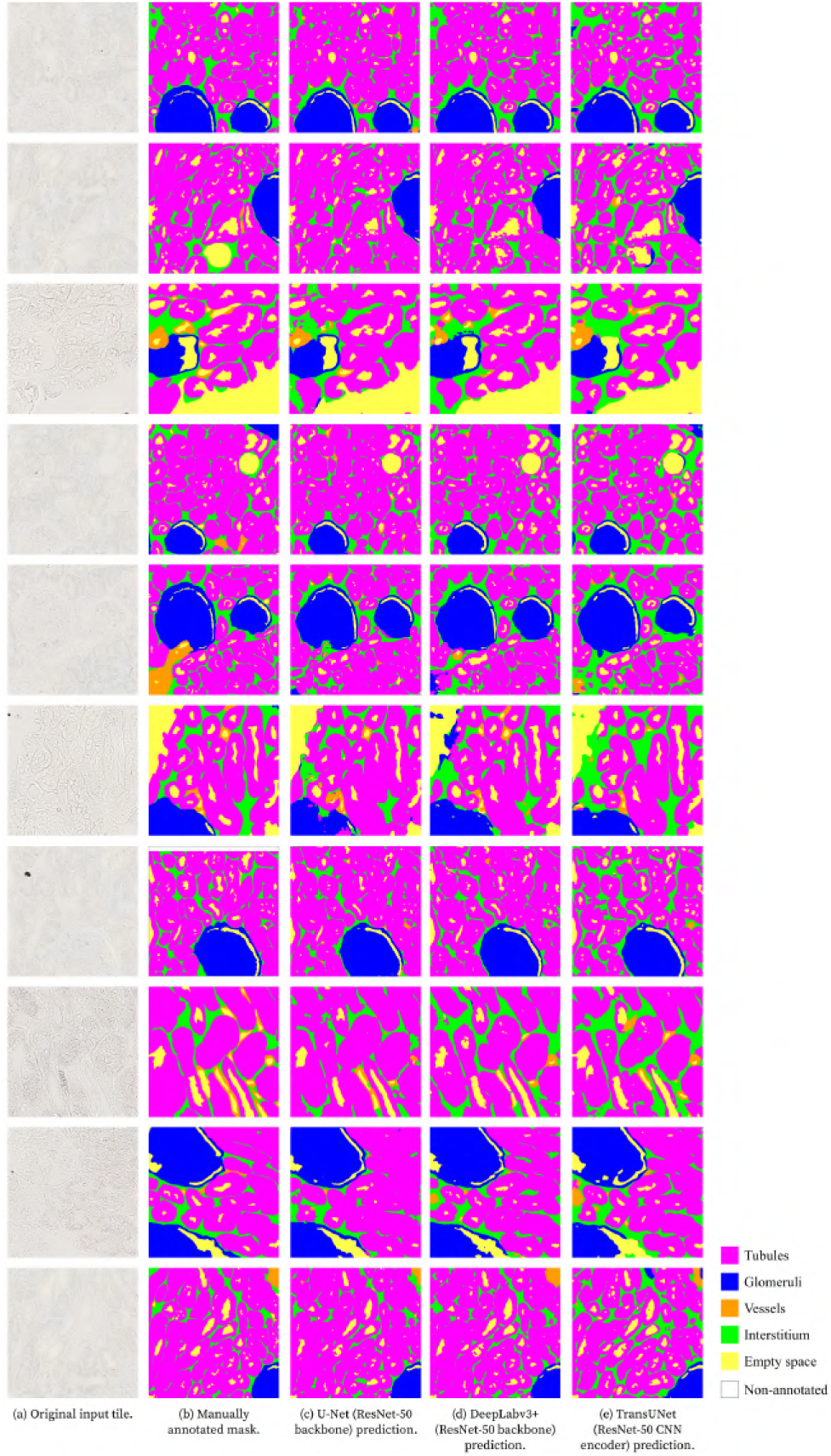
Multi-class segmentation masks output using the supervised deep learning models and respective manual annotations, shown for ten different example images (top, middle, and bottom rows). For each image: (a) Original image, (b) Manually annotated masks, (c) U-Net (*L*_Dice-CE_), (d) DeepLabv3+ (*L*_Dice-Focal_), and (e) TransUNet (*L*_Dice-CE_). *Colour code: tubules (magenta), glomeruli (blue), vessels (orange), interstitium (green), empty space (yellow), and non-annotated tissue (white)*.

**Table 27:** Average performance metrics results after post-processing for multi-class segmentation with DeepLabv3+ (ResNet-50 backbone) and *L*_Dice-Focal_ loss function.

| **Class** | **IoU** (%) | **DSC** (%) | **Accuracy** (%) | **Precision** (%) | **Recall** (%) |
|---|---|---|---|---|---|
| *Tubule* | 84.2 | 91.3 | 90.1 | 91.4 | 91.4 |
| *Glomerulus* | 67.4 | 73.2 | 98.4 | 92.2 | 72.6 |
| *Vessel* | 11.5 | 17.8 | 98.5 | 20.6 | 20.4 |
| *Interstitium* | 42.4 | 58.8 | 91.0 | 56.6 | 63.7 |
| *Empty space* | 63.7 | 76.8 | 97.2 | 78.5 | 76.8 |

**Figure 31:**
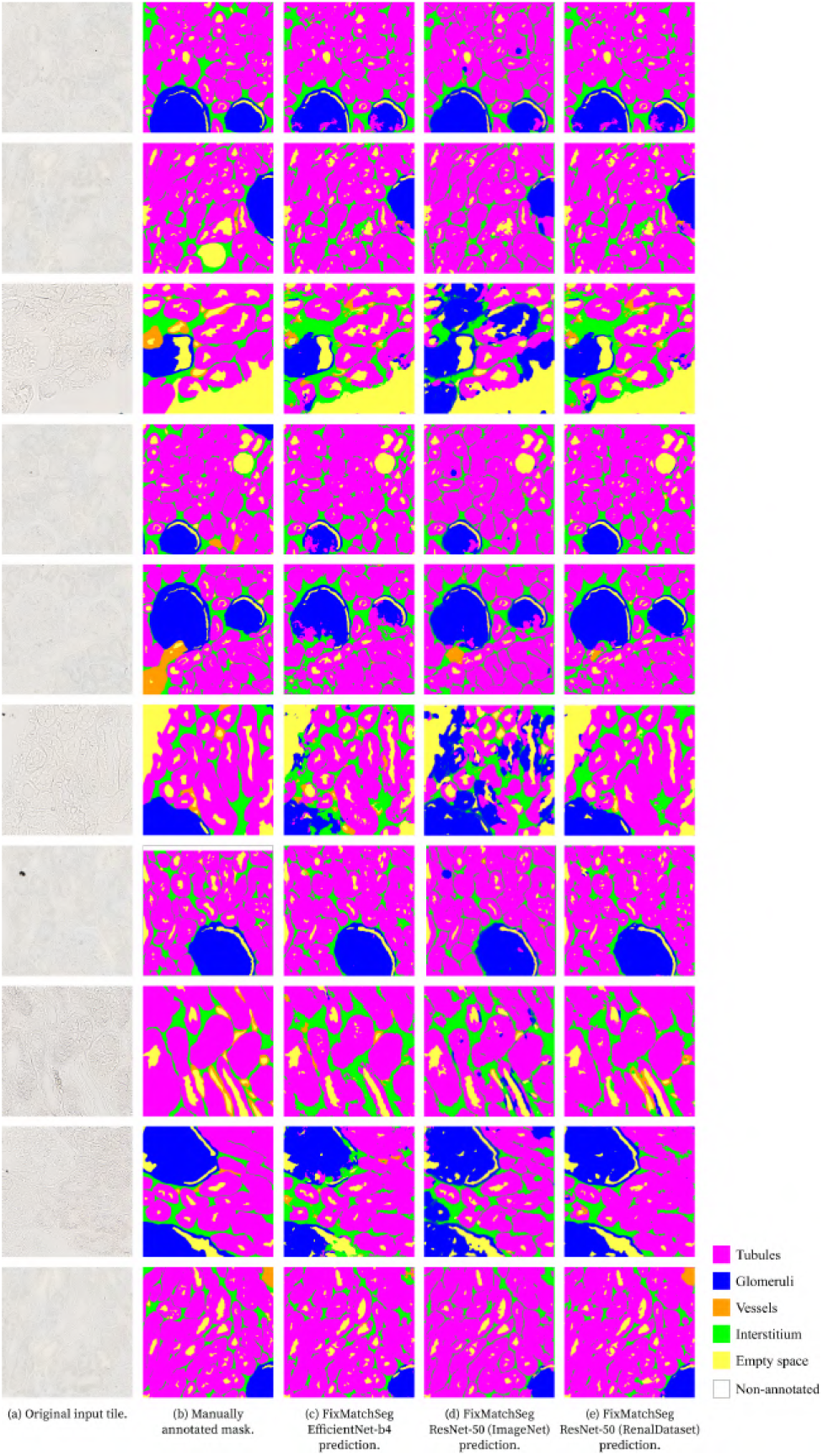
Multi-class segmentation masks output using the FixMatchSeg methods and respective manual annotations, shown for ten different example images (rows). For each image: (a) Original image, (b) Manually annotated masks, (c) FixMatchSeg with EfficientNet-b4 backbone, (d) FixMatchSeg with ResNet-50 backbone initialised with ImageNet weights, and (e) FixMatchSeg with ResNet-50 backbone initialised with RenalDataset weights. *Colour code: tubules (magenta), glomeruli (blue), vessels (orange), interstitium (green), empty space (yellow), and non-annotated tissue (white)*.

**Table 28:** Average performance metrics results after post-processing for multi-class segmentation with TransUNet and *L*_Dice-CE_ loss function.

| **Class** | **IoU** (%) | **DSC** (%) | **Accuracy** (%) | **Precision** (%) | **Recall** (%) |
|---|---|---|---|---|---|
| *Tubule* | 80.3 | 89.0 | 87.9 | 95.0 | 84.0 |
| *Glomerulus* | 68.8 | 74.4 | 98.5 | 90.7 | 72.6 |
| *Vessel* | 10.5 | 17.1 | 98.0 | 16.2 | 25.3 |
| *Interstitium* | 40.4 | 56.7 | 88.3 | 47.3 | 75.0 |
| *Empty space* | 62.4 | 75.9 | 96.9 | 70.4 | 84.9 |

